# Redox-controlled dimerisation regulates ethylene biosynthesis

**DOI:** 10.64898/2025.12.04.692314

**Authors:** Dona M. Gunawardana, Francis Kuang, Simranjeet Kaur, Jordan A. P. McIvor, Xiaoxu Chen, Yuliana Yosaatmadja, Ashish Sethi, Henry Y. H. Tang, Martin J. Middleditch, Davide Mercadante, Michael J. Haydon, Christopher J. Squire, Ivanhoe K. H. Leung

## Abstract

Ethylene is a central plant hormone that orchestrates growth, development, senescence, and stress responses. Because it is gaseous, ethylene must be synthesised on demand, yet the catalytic and regulatory mechanisms of its biosynthetic enzyme, 1-aminocyclopropane-1-carboxylic acid oxidase (ACO), remain poorly understood. Here, using structural, biophysical, and computational analyses, we uncovered two principles: ACO catalysis relies on an induced-fit mechanism, and disulfide-mediated dimerisation via a conserved cysteine acts as a redox switch toggling ACO between active monomer and inactive dimer. This previously unrecognised regulatory layer positions ACO as a redox sensor in plant cells, revealing a fundamental control point in ethylene biosynthesis. Given ethylene’s pivotal role in crop productivity and stress resilience, these findings open new opportunities for precise manipulation of hormone signalling in agriculture and biotechnology.

## Introduction

Ethylene (C_2_H_4_) is a gaseous phytohormone that integrates developmental programs with stress signalling in plants^1–3^. Because it cannot be stored, ethylene biosynthesis is tightly regulated, both temporally and spatially, to meet physiological demands^4–6^. The final step of this pathway is catalysed by 1-aminocyclopropane-1-carboxylic acid (ACC) oxidase (ACO) (Figure 1a)^7,8^, which emerging evidence identifies as a rate-limiting enzyme during specific developmental stages and under stress conditions^6,8^. Modulating ACO abundance or activity, and thus ethylene availability, can enhance tolerance to abiotic stresses such as salinity, drought, flooding, and extreme temperatures^8–11^, while ACO inhibition has been exploited postharvest to delay ripening and extend shelf life^12^.

**Figure 1.**
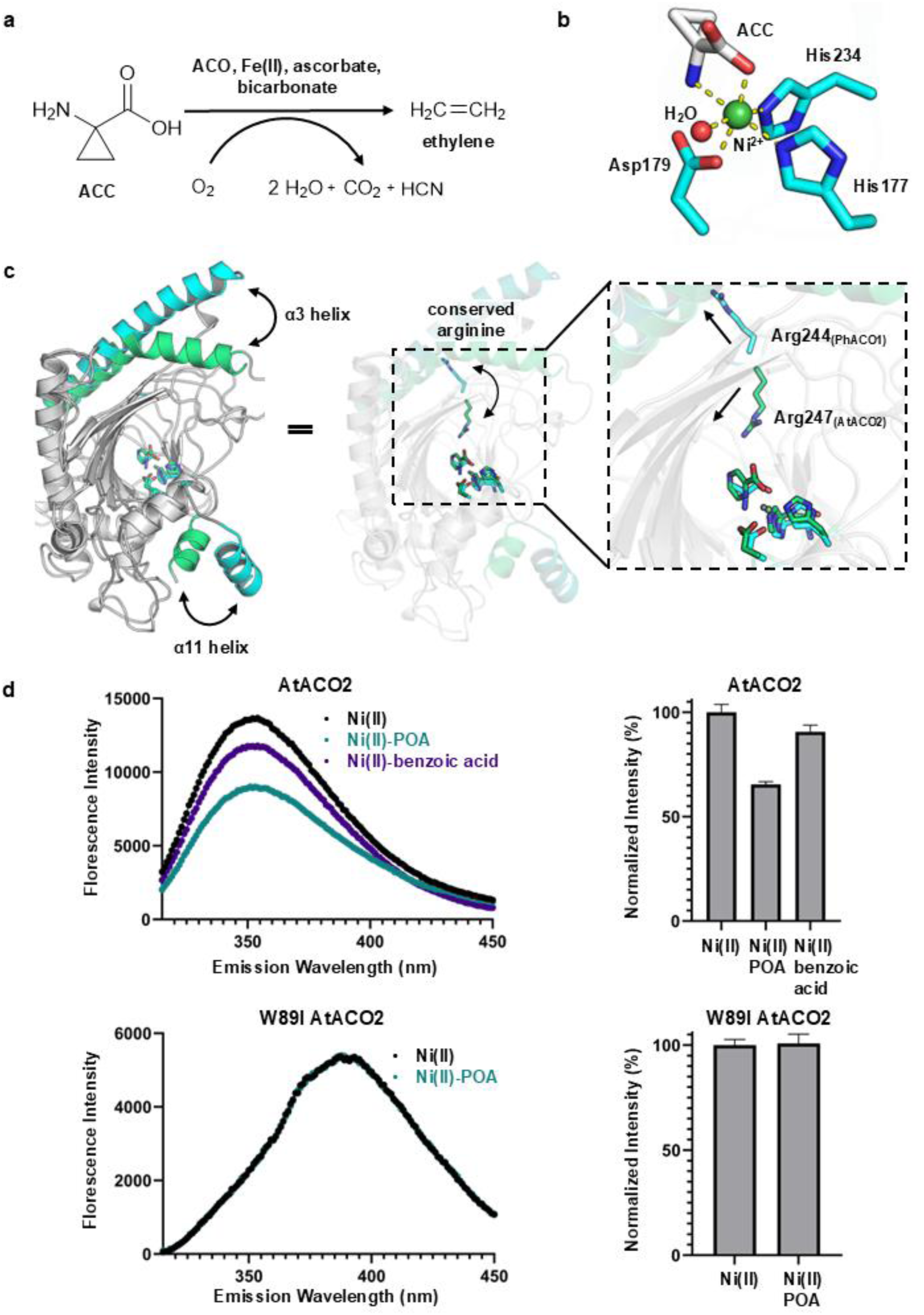
Ligand-induced conformational changes enable ACO-mediated ethylene synthesis. (a) ACO converts ACC to ethylene using molecular oxygen, producing water, carbon dioxide, and cyanide as by-products. The reaction requires Fe(II), ascorbate, and bicarbonate cofactors. (b) ACC binds the active-site metal ion in a bidentate fashion, with a water molecule occupying the sixth coordination site. (c) Structural overlay of PhACO1–Ni(II)–ACC (PDB: 5TCV, cyan) and AtACO2–Zn(II)–POA (PDB: 5GJ9, green) highlights differences in helices α3 and α11 and the conserved arginine within the RXS motif. (d) Intrinsic tryptophan fluorescence of AtACO2 upon ligand binding. POA induces marked quenching in wild-type AtACO2, whereas benzoic acid (non-binder) shows minimal effect. The W89I mutant exhibits no fluorescence change with POA. Data represent mean ± s.d. from three independent experiments (5 µM AtACO2, 50 µM Ni(II), 500 µM POA where applicable).

ACO catalyses the two-electron oxidation of ACC to produce ethylene, carbon dioxide, and cyanide, using molecular oxygen as a co-substrate and Fe(II), ascorbate, and bicarbonate as co-factors (Figure 1a)^13–16^. Structurally, ACO belongs to the 2-oxoglutarate (2OG) and Fe(II)-dependent dioxygenase (2OGD) superfamily^7,13^, and adopts the characteristic distorted double-stranded β-helix (DSBH) fold with the conserved 2-His-1-carboxylate Fe(II) binding motif common to all 2OGDs^13^. However, ACO is an atypical member of this superfamily: unlike canonical 2OGDs, which require 2OG as a co-substrate, ACO directly utilises ACC and molecular oxygen, and uniquely requires bicarbonate (Figure 1a and Supplementary Figure S1)^13–16,17,18^.

Despite ACO’s central role in plant biology, its molecular mechanism and regulation remain poorly defined. Crystal structures of *Petunia hybrida* ACO1 (PhACO1) and *Arabidopsis thaliana* ACO2 (AtACO2) suggested multiple enzyme conformations^13,19^, yet subsequent computational and spectroscopic failed to corroborate these observations^20^. ACO has also been reported to exist in multiple oligomeric states^13,16,19^, but the biological significance of this heterogeneity remains unclear, limiting rational strategies to manipulate ethylene output for crop improvement.

Here, we uncover a previously uncharacterised regulatory mechanism of ACO at the molecular level. Using X-ray crystallography, we solved the structure of PhACO1 bound to its substrate ACC. Supported by biophysical assays, mutagenesis, and computational modelling, we show that ACO undergoes large-scale, ligand-induced conformational change essential for catalysis, with redox-mediated dimerisation acting as a molecular switch between active monomer and inactive dimer. This unprecedented mode of regulation positions ACO as a cellular redox sensor, revealing a fundamental control point in ethylene biosynthesis and providing new molecular targets for improving crop resilience and postharvest quality.

## Results

### ACC binds to ACO by chelating the active site metal

To understand how ACO engages its substrate, we first determined the substrate-free structure of *Petunia hybrida* ACO1 (PhACO1) bound to Ni^2^⁺ (Supplementary Table S1; PDB: 5TCW). Substituting Fe^2^⁺ with Ni^2^⁺ is a standard strategy for 2OGD enzymes, as Ni^2^⁺ preserves coordination geometry while remaining redox-inactive, preventing oxidative turnover during crystallography^21^. Ni^2^⁺ also provided the greatest structural stability among tested metals (Supplementary Fig. S2). PhACO1 crystallised as an apparent tetramer, consistent with previous reports (Supplementary Fig. S3). In our structure, Ni^2^⁺ adopts a distorted octahedral geometry coordinated by His177, Asp179, and His234 in the canonical facial arrangement, plus a monodentate acetate from the crystallisation buffer; two sites remain vacant (Supplementary Fig. S4). This configuration matches the Fe^2^⁺-bound PhACO1 structure (PDB: 1WA6; Supplementary Fig. S4)^13^, validating Ni^2^⁺ substitution.

We next visualised ACC binding by soaking pre-formed PhACO1–Ni^2^⁺ crystals with ACC, yielding a 2.6 Å structure (Supplementary Table S2; PDB: 5TCV). ACC replaces acetate and chelates Ni^2^⁺ in a bidentate mode via its amino and carboxylate groups (Fig. 1b). The metal centre retains a distorted octahedral geometry, with a single water molecule completing the coordination sphere, presumably in the oxygen-binding site required for catalysis. This model provides a refined catalytic mechanism whereby the formation of a highly oxidising Fe(IV)-oxo intermediate drives the oxidative decarboxylation of ACC (Supplementary Fig. S1). This arrangement contrasts with some earlier models proposing ACC binding equatorially with the two histidine ligands and oxygen trans to the aspartate (Supplementary Fig. S1)^14^; instead, ACC binds equatorially alongside His177 and Asp179 (Fig. 1b). These data define the substrate-binding mode and set the stage for mechanistic analysis of ACO catalysis.

### ACO undergoes ligand-induced conformational change

Our PhACO1-Ni(II)-ACC structure represents the first ACO structure with its native substrate. However, ACO structures with other ligands have been reported. Sun et al. solved an *Arabidopsis thaliana* (AtACO2) structure in complex with Zn^2+^ and inhibitors pyrazinecarboxylic acid (POA) or 2-picolinic acid (PDB: 5GJ9, 5GJA) (Supplementary Figure S5)^19^. Despite 82% sequence similarity between PhACO1 and AtACO2 (Supplementary Figure S6), their structures differ markedly in three regions: the α3 helix, the C-terminal α11 helix, and the conserved RXS motif (Figure 1c). In PhACO1, α3 and α11 adopt extended conformations, whereas in AtACO2 they are more compact. Strikingly, the conserved arginine in the RXS motif (Arg 244 in PhACO1; Arg247 in AtACO2) points away from the active site in PhACO1 but toward the ligand-bound metal centre in AtACO2 (Figure 1c). In most ligand-bound 2OGD structures, this basic residue interacts with the terminal carboxylate group of cosubstrate 2OG or inhibitors such as NOG (Supplementary Figure S7)^21^. Large ligand-induced rearrangements are well documented in other 2OGDs^22–25^, suggesting that ACO similarly toggles between open and closed states. We propose that our PhACO1 crystal structures capture an open, substrate-accessible conformation, whereas AtACO2-ligand complexes represent a closed, catalytically competent state. This conformational plasticity likely underpins ACO’s induced-fit mechanism.

To test whether ligand binding triggers structural rearrangement, we used protein NMR spectroscopy^26^ on AtACO2 with Zn^2+^ and POA as model ligands. Comparison of ^1^H-^15^N transverse relaxation-optimised spectroscopy (TROSY) spectra of ^2^H,^15^N-labelled AtACO2-Zn^2+^ in the presence and absence of POA revealed a global shift in backbone amide resonances (Supplementary Figure S8), consistent with a conformational change upon ligand binding. However, detailed mapping was precluded by broad resonance arising from the high molecular mass of the protein (∼38 kDa)^27^.

To further probe conformational dynamics, we used intrinsic tryptophan fluorescence, a sensitive reporter of local environment changes^28,29^. Both PhACO1 and AtACO2 contain three tryptophans (Trp31, Trp86 and Trp203 in PhACO1; Trp34, Trp89 and Trp207 in AtACO2), all >10 Å from the active site (Supplementary Figure S9). These tryptophan residues are located at least 10 Å away from the active site (Supplementary Figure S9). Two residues (Trp31/Trp203 and Trp34/Trp207) lie buried in the hydrophobic core and align well between PhACO1-Ni^2+^ and AtACO2-Zn^2+^-POA structures. In contrast, Trp86 (PhACO1) and Trp89 (AtACO2) reside on the α3 helix, which shifts markedly between open and closed states. Notably, Trp89 in AtACO2-POA sits within 3.6 Å of Phe68, a potential fluorescence quencher, whereas Trp86 in PhACO1 is >8 Å from any aromatic residue. This suggested Trp86/Trp89 could serve as reporters for ligand-induced conformational change.

Consistent with this, both enzymes displayed strong fluorescence in the presence of Ni^2^⁺ (used as an Fe^2^⁺ surrogate). Addition of POA (20-fold molar excess) quenched fluorescence in PhACO1 and AtACO2 without shifting the emission maximum (Fig. 1d; Supplementary Fig. S10), consistent with quenching rather than solvent exposure changes. These data support a ligand-induced rearrangement involving the α3 helix.

To exclude non-specific surface binding of POA as the source of fluorescence quenching, we tested benzoic acid, a POA analogue lacking the ring nitrogen required for metal chelation (Supplementary Fig. S11). Isothermal titration calorimetry confirmed no binding to ACO (Supplementary Fig. S12), and benzoic acid did not quench fluorescence (Fig. 1d). Finally, we generated an AtACO2 W89I mutant to probe the role of Trp89. Addition of POA to W89I-Ni^2^⁺ caused no fluorescence change (Fig. 1d). Together, these controls indicate that quenching in wild-type enzymes reflects environmental changes around Trp86(PhACO1)/Trp89(AtACO2) driven by ligand-induced conformational rearrangement.

To test whether POA binding alters global shape, we performed small-angle X-ray scattering (SAXS) on PhACO1 with and without POA. Scattering profiles differed markedly between PhACO1-Ni^2^⁺ and PhACO1-Ni^2+^-POA (Supplementary Fig. S13). Both the radius of gyration (*R*_g_) and maximum dimension (D_max_) decreased upon POA binding (*R*_g_: 29.81 → 27.59 Å; *D*_max_: 116.61 → 107.87 Å), indicating a more compact structure. This agrees with solvent-accessible surface area and volume analyses ^30^ of PhACO1-Ni^2^⁺ (PDB ID: 5TCW; open), AtACO2–Zn^2^⁺–POA (PDB ID: 5GJ9; closed), and an AlphaFold model of closed PhACO1 (volumes: 264 Å^3^, 200 Å^3^, and 227 Å^3^, respectively; Supplementary Fig. S14). To further capture the conformational change of PhACO1 in solution, we employed the Ensemble Optimisation Method (EOM)^31^ to generate a pool of possible conformation (Supplementary Fig. S13) following POA binding. In the absence of POA, PhACO1 existed in an opened conformation with higher D_max_ (Supplementary Fig. S13). Upon POA binding, PhACO1 transitioned to a closed or compact conformation with lower D_max_ (Supplementary Fig. S13). These findings align to our intrinsic tryptophan fluorescence quenching assay that addition of POA induces a closed conformation. Moreover, both samples showed a consistent peak with higher D_max_ (Supplementary Fig. S13), which is highly correlated to the flexibility of the C-terminal. Together, these data support our hypothesis that ligand binding drives an open-to-closed transition.

### Molecular dynamics simulations reveal a ligand stabilised salt-bridge network

To probe how POA binding promotes the closed conformation, we performed molecular dynamics (MD) simulations on AtACO2-Ni^2^⁺ in apo and POA-bound states (PDB: 5GJ9) ^19^. Analysis of key salt bridges (Glu53–Lys194, Glu90–Lys161, Asp182–Lys291, Lys161–Asp182, Lys75–Asp86) revealed that removing POA weakens several interactions, increasing inter-residue distances (Supplementary Fig. S15). Notably, Lys161–Asp182, which are critical for metal coordination, and Asp182–Lys291 near the C-terminal helix showed the largest shifts, consistent with structural comparisons between PhACO1 and AtACO2 and prior observations by Fournier et al.^20^. Conversely, Glu90–Lys161, linking the α3 helix loop to the active site, strengthened in the apo state, suggesting a rearrangement of the salt-bridge network upon ligand binding. These changes highlight a stabilising role for POA and again implicate α3 in the induced-fit mechanism (Fig. 1F).

### α3 helix movement links ligand binding to catalysis

Our analyses suggested that bending of the α3 helix is central to ACO’s induced-fit mechanism. In POA-bound AtACO2, Arg247 of the RXS motif occupies the position of Tyr162 in PhACO1 and forms a hydrogen bond with Ser249 (Fig. 2a and b). This rearrangement displaces Tyr165, which hydrogen bonds to Glu90 on the α3 loop. Glu90 is further stabilised by a charge–charge interaction with Lys161, which also forms a salt bridge to POA, and by analogy, to ACC (Fig. 2c). Collectively, these interactions create a tether from the ligand-binding site, through the central β-sheet, to α3, bending the helix and pulling its tip up to 17 Å toward the active site (Fig. 2a–c). This coupling provides a plausible route by which substrate binding drives closure and positions catalytic residues for reaction.

**Figure 2.**
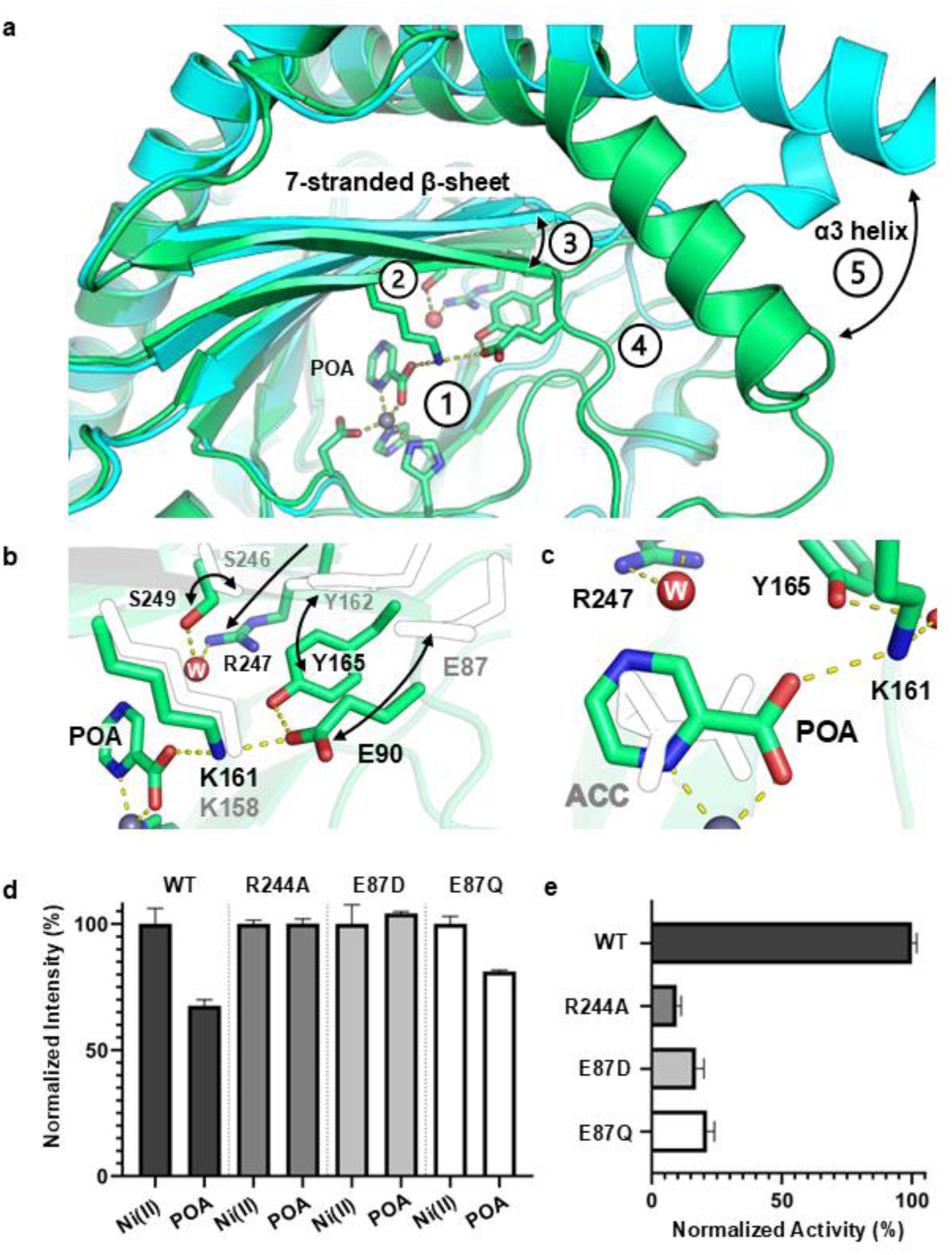
Rearrangement of active-site residues underlies ligand-induced conformational change. (a) Overlay of PhACO1–Ni(II)–ACC (cyan) and AtACO2–Zn(II)–POA (green) structures reveals five key changed on ligand binding (labelled 1-5). POA makes new contacts at location ➀ that rearrange the central β-sheet at ➂ and the loop region between the active site and α3 at location ➃. The largest movement reorients Arg247 toward the active site at ➁, displacing Tyr165 and altering α3 helix orientation relative to the β-sheet. Collectively, the rearrangements bend α3 helix as it is pulled in a large-scale movement at ➄ towards the active site. (b) A zoomed view showing the residue movements (indicated by arrows) associated with POA binding. (c) Comparison of ACC and POA binding shows Arg247, Tyr165, and Lys161 positioned to either ligand. (d) Intrinsic tryptophan fluorescence quenching of PhACO1 variants upon POA addition. Wild-type shows significant quenching, as does E87Q although reduced. R244A and E87D show no response and indicate the elimination of critical important ligand binding elements. Samples included 5 µM PhACO1, 50 µM Ni(II) and 200 µM POA (where applicable). (e) Single-point activity assay of PhACO1 mutants relative to wild-type. Substitution of Arg244 or Glu87 markedly decreases catalytic activity. Reactions contained 2 µM enzyme, 20 µM Fe(II), 250 µM ACC, 12.5 mM ascorbic acid, 30 mM bicarbonate, and 250 µg catalase in Tris-D11 buffer (pH 7.5) with 10% H_2_O and 90% D_2_O.Data represent mean ± s.d. from three independent experiments.

To test whether this network controls conformational change, we engineered three PhACO1 variants targeting its key nodes: R244A, E87D, and E87Q (Arg244 and Glu87 in PhACO1 correspond to Arg247 and Glu90 in AtACO2). R244A is predicted to disrupt the Arg–Ser contact (Arg244–Ser246). E87Q neutralises the Glu87 side chain, abolishing the charge-charge interaction with Lys158 (AtACO2 Lys161 equivalent), while E87D shortens the side chain, weakening that same interaction (Fig. 2a and b).

We then assayed ligand-induced closure using intrinsic tryptophan fluorescence. Unlike wild-type PhACO1, which shows robust quenching upon POA addition in the presence of Ni^2+^, none of the mutants exhibited significant quenching (Fig. 2d), indicating loss of the conformational response. To confirm that this effect reflects impaired closure rather than loss of ligand binding, we performed NMR-based binding experiments, which showed POA binds to wild-type and all three mutants (Supplementary Fig. S16). ITC further demonstrated similar affinities for wild-type and R244A (*K*_D_ = 69.5 µM and 104 µM, respectively; Supplementary Figs. S17 and S18).

Finally, we quantified catalysis using a ^1^H NMR assay monitoring ACC consumption under standard cofactor conditions (Fe^2^⁺, bicarbonate, ascorbate). Wild-type PhACO1 readily converted ACC to ethylene, whereas all three mutants displayed ≥80% loss of activity (Fig. 2e; Supplementary Fig. S19). Together, these data show that the α3-linked interaction network is required for ligand-induced closure, and that closure is directly coupled to ACO catalysis.

### PhACO1 undergoes redox-mediated dimerisation

Several ACOs from diverse plant species, including PhACO1, are reported to exist in solution as mixtures of oligomeric states^13,16,19^. Consistent with these observations, recombinant PhACO1 consistently resolved as two species during size-exclusion chromatography purification, despite eluting as a single peak in IMAC (Fig. 3a; Supplementary Fig. S20). SDS-PAGE confirmed both fractions contained PhACO1 alone. Because oligomerisation is a common mechanism of protein-level regulation^34,35^, we decided to investigate this phenomenon in detail.

**Figure 3.**
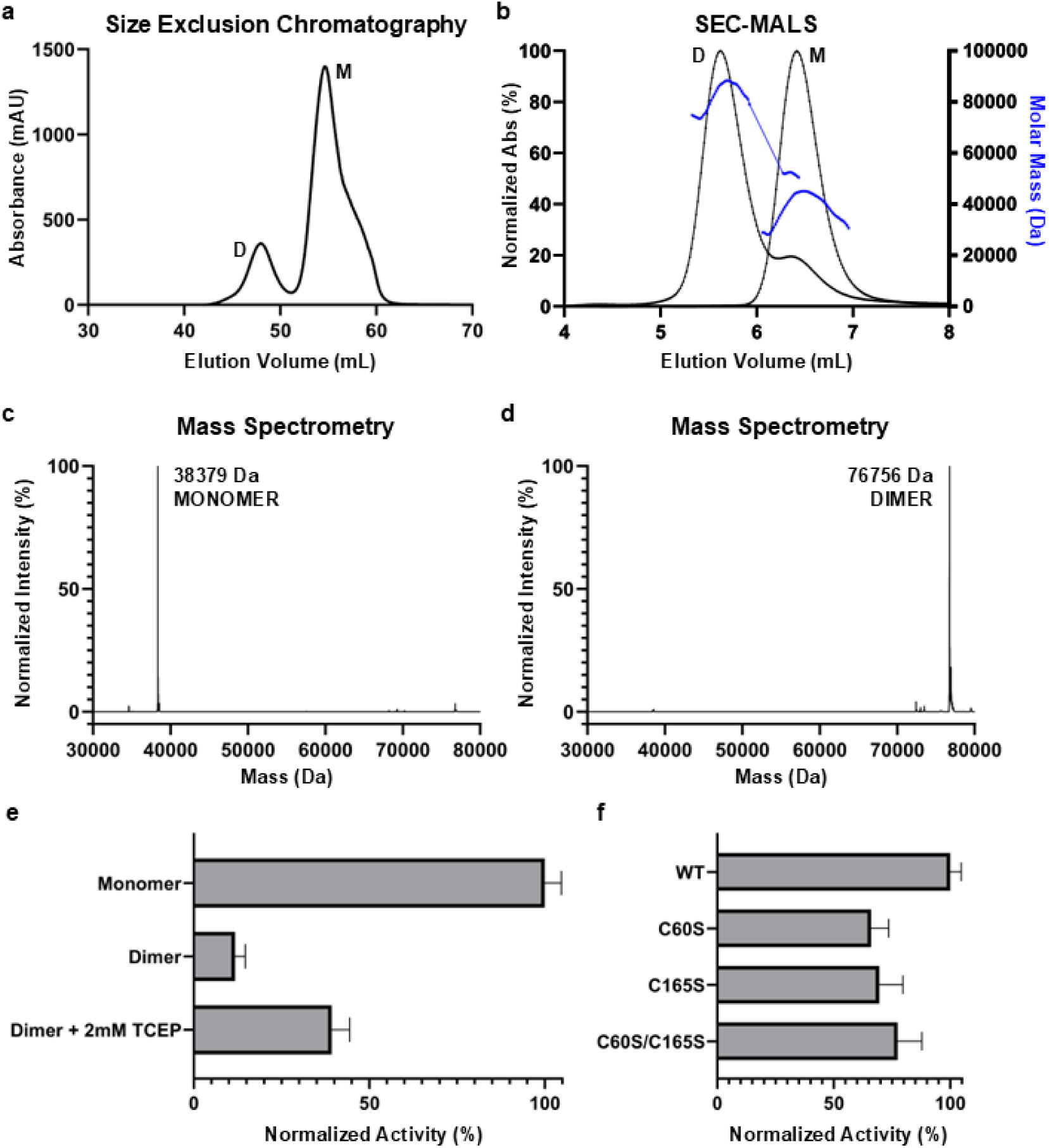
Redox-mediated dimerisation modulates PhACO1 activity. (a) Size-exclusion chromatography of PhACO1 revealed two oligomeric states: a major monomer peak (M) and a minor dimer peak (D). (b) Size exclusion chromatography coupled with multi-angle light scattering (SEC-MALS) confirmed the species identities: monomer (∼40 kDa) and dimer (∼80 kDa) eluting at distinct volumes. (c) Intact protein mass spectrometry of the monomer species shows a single peak at 38,379 Da, matching the calculated mass after N-terminal methionine removal (38,377 Da). (d) Intact protein mass spectrometry of the dimer species shows a peak at 76,756 Da, consistent with two monomers linked by disulfide bond. (e) Normalised enzyme activity of monomer and dimer with or without 2 mM TCEP. Activity was measured using a single-concentration assay monitoring ACC consumption our 2 hours in a mixture containing 2 µM enzyme, 20 µM Fe(II), 250 µM ACC, 12.5 mM ascorbic acid, 30 mM bicarbonate, and 250 µg catalase in Tris-D11 buffer (pH 7.5) in 10%/90% H_2_O/D_2_O. Dimer exhibits massively reduced activity, which is partially restored upon reduction. (f) Normalised enzyme activity of cysteine mutants (C60S, C165S, and C60S/C165S) compared to wild-type. Cysteine substitution significantly diminishes dimerisation-dependent regulation. Data represent mean ± s.d. from three independent experiments; wild-type monomer activity defined as 100%.

Using SEC-MALS, the two species from size exclusion chromatography were assigned as monomer (∼40 kDa) and dimer (∼80 kDa) (Fig. 3b). Intact protein MS under denaturing conditions gave masses of 38,379 Da and 76,756 Da, matching a monomer lacking the N-terminal methionine and a dimer 2 Da lighter than expected. The mass difference in the dimer is consistent with a disulfide linkage having formed (Fig. 3c,d). Thus, PhACO1 exists as a mixture of monomers and disulfide-linked dimers in solution.

Treatment with tris(2-carboxyethyl)phosphine (TCEP), a selective reducing agent for disulphide bonds^37^, converted dimers to monomers (Supplementary Figures S21 and S22), and was unaffected by cofactors or substrates (Supplementary Figure S23). Glutathione partially reduced dimers and generated a glutathionylated monomer peak (Supplementary Fig. S24), whereas ascorbate, despite being an ACO cofactor, failed to reduce dimers even at 10 mM (Supplementary Fig. S25), a condition that has been reported to reduce disulphides in model systems^38–40^. This suggests that the dimer is stable under physiological ascorbate levels and likely requires dedicated cellular reductants such as glutathione to effect change *in planta*. Conversely, incubation with H_2_O_2_, promoted dimer formation (Supplementary Fig. S26). Hydrogen peroxide is widely used to mimic oxidising conditions for studying redox-active enzymes^41–43^, is a biologically relevant oxidant and signalling molecule in plants^44,45^, and has been implicated in regulating ethylene biosynthesis as well as other ethylene-related processes^46–48^. Together, these data demonstrate that PhACO1 oligomeric state is redox-sensitive, with oxidative conditions favouring disulphide-linked dimers and reducing environments favouring monomers.

### Cys165 and Cys60 mediate redox-controlled dimerisation

To identify the cysteines responsible for dimerisation, we performed tandem MS/MS on PhACO1 dimers. Two disulfide-linked species were detected: a predominant homodimer involving Cys165 from each monomer, and a less abundant heterodimer (∼7-fold lower) linking Cys165 to Cys60 (Supplementary Fig. S27). Both residues are conserved among ACOs (Supplementary Fig. S28), and structural analysis confirmed their solvent accessibility (Supplementary Fig. S29).

To validate their roles, we generated C60S, C165S, and C60S/C165S mutants. Size-exclusion chromatography without reducing agents showed that C165S and the double mutant existed exclusively as monomers, whereas C60S retained ∼20% dimer (Supplementary Fig. S30). Under oxidising conditions (e.g., in the presence of H_2_O_2_), both single mutants could still dimerise, but dimerisation was abolished in the double mutant (Supplementary Fig. S31). These data indicate that Cys165 is the primary residue mediating dimerisation, with Cys60 contributing under certain conditions.

### PhACO1 monomer is the active species

To assess functional relevance, we separated monomeric and dimeric PhACO1 and measured activity. The monomer readily catalysed ACC conversion, whereas the dimer was inactive (Fig. 3e). Reducing the dimer back to monomer restored ∼50% activity, likely reflecting incomplete reduction or partial denaturation. All cysteine mutants displayed activity comparable to monomeric wild type (Fig. 3f). Thus, PhACO1 activity is gated by redox-controlled dimerisation, a regulatory mechanism not previously observed in 2OG oxygenases.

### Dimerisation locks ACO in an open, inactive state

Crystallisation of purified dimers was unsuccessful, so we modelled the dimer using protein-protein docking, while satisfying a 4 Å constraint between Cys165 residues (Supplementary Fig. S32), reflective of a disulfide bond formation. The docking procedure could only satisfy the constraint for the open-state monomer (PhACO1-Ni^2+^; PDB: 5TCV), even when the S-S distance was relaxed to 10 Å. The relaxation of the docked pose with the lowest energy score, via MD simulations, revealed an electrostatic interface involving residues that normally undergo conformational rearrangement during catalysis, including Glu87 (forming an intermolecular salt bridge with Lys167) and Arg244 (hydrogen bonding to Thr82) (Supplementary Fig. S33). The α3 helix remained rigid in the dimer, suggesting that these interactions restrict the conformational flexibility required for induced fit, explaining the inactivity of the dimer.

### Glutathione and cysteinyl-glycine modify ACO in plant lysate

To test physiological relevance, we incubated monomeric PhACO1 in *Arabidopsis* ET free mutant cell lysate lacking functional ACOs^50^. Both wild type and C60S formed dimers, whereas C165S and the double mutant remained monomeric (Fig. 4a). Dimer levels were lower than *in vitro* (Figure 4a and Supplementary Figures S26 and S34) and adding H_2_O_2_ did not increase dimerisation. Surprisingly, PhACO1 recovered and purified from the plant cell lysate resisted dimerisation in buffer, suggesting post-translational modification. Intact protein mass spectrometry revealed mass shifts (+178, +306, +354, +484 m/z) consistent with glutathione or cysteinyl-glycine adducts (Fig. 4b), which were reversible by application of TCEP (Supplementary Fig. S35). Glutathione and cysteinyl-glycine are both biologically important redox molecules in plants cells^51^ and they have been implied in functions such as fruit ripening^52^. Purification of recombinant protein with oxidised glutathione or cysteinyl-glycine confirmed these modifications (Supplementary Fig. S36), and mutagenesis showed Cys165 as the primary site (Supplementary Fig. S37). Importantly, these modifications did not impair activity (Fig. 4c), indicating that glutathione/cysteinyl-glycine locks ACO in its monomeric, active form and that Cys165 is a key redox-sensitive switch in PhACO1, through which dimerisation inhibits protein activity (Figure 5),

**Figure 4.**
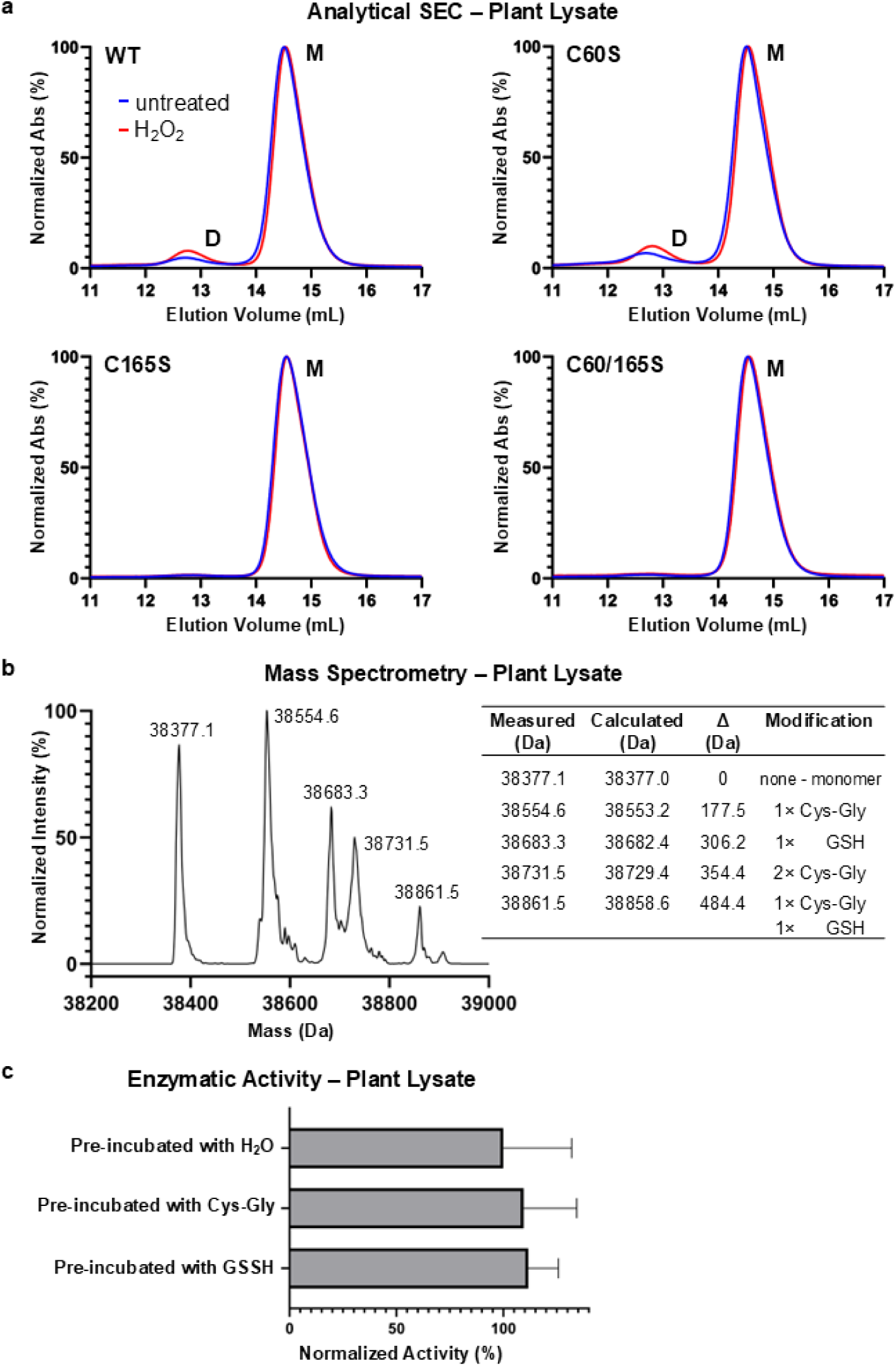
Plant-derived oxidative conditions modify catalytic cysteines in PhACO1. (a) Analytical size-exclusion chromatography of PhACO1 variants after incubation in Arabidopsis plant lysate with or without 2 mM H₂O₂. The wild-type and C60S show a shift toward dimer formation under oxidative conditions, whereas C165S, and C60S/C165S mutants remain monomeric. (b) Intact protein mass spectrometry of PhACO1 recovered from plant lysate reveals multiple mass shifts corresponding to cysteine modifications: +177.5 Da (Cys-Gly), +305.4 Da (GSH), and combinations thereof. (c) Normalised enzymatic activity of modified PhACO1 compared to unmodified enzyme (carried out in the same conditions as Fig. 3e). Pre-incubation with Cys-Gly or GSH significantly increases activity relative to untreated control. Data represent mean ± s.d. from three independent experiments; unmodified PhACO1 activity is defined as 100%.

**Figure 5.**
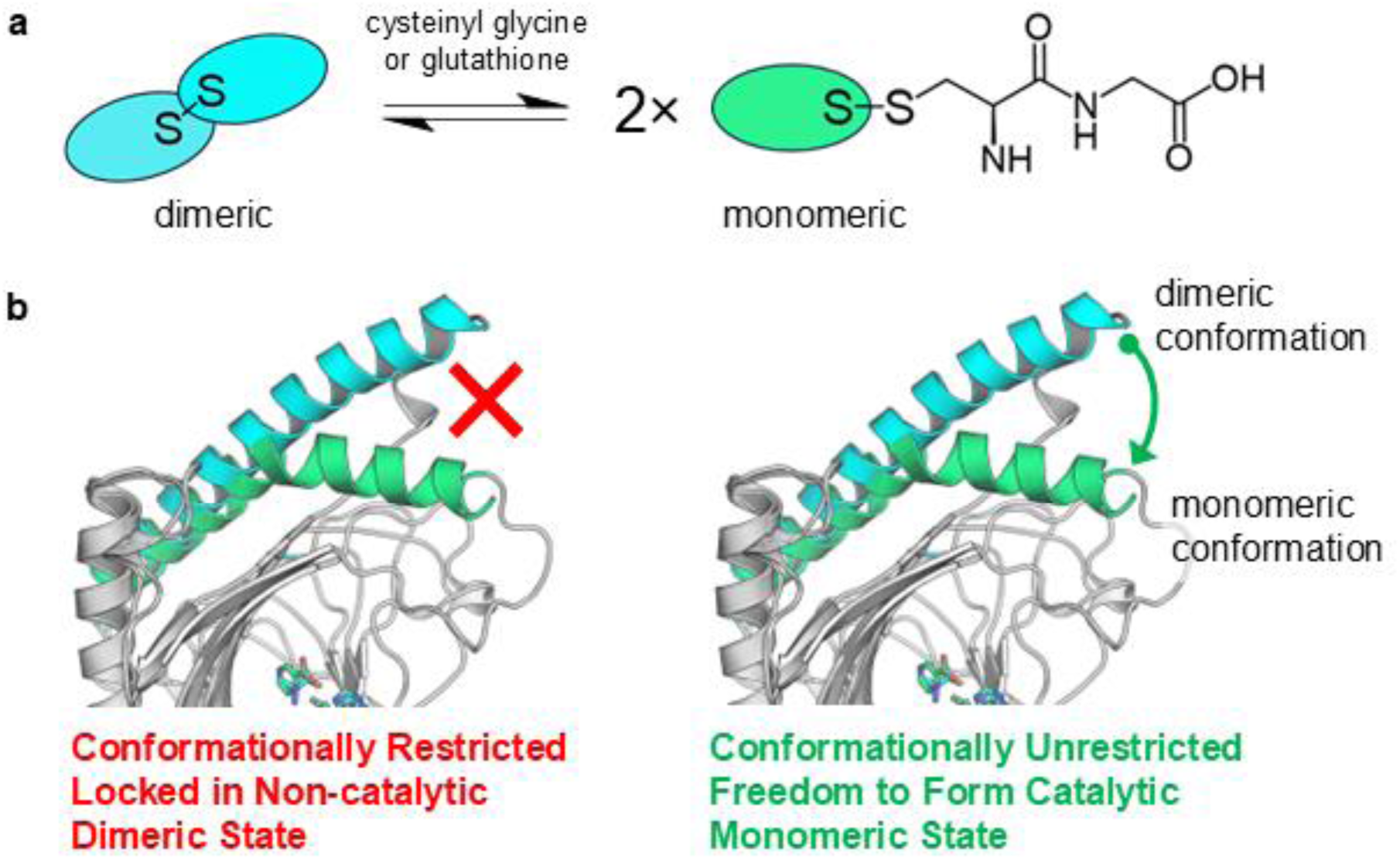
Proposed protein-level regulatory mechanism of PhACO1. (a) Oxidative conditions promote disulfide-linked dimer formation, whereas reducing agents or thiol modifications (cysteinyl glycine or glutathione) favour the monomeric state. (b) Structural representation of conformational states related to redox status: oxidative-dimerisation locked the α3 helix and the enzyme in a non-catalytic conformation (left), while the monomer retains the conformational freedom required for catalysis on ligand binding (right).

### PhACO1 dimerisation occurs in planta

Finally, we tested PhACO1 dimerisation in *Nicotiana benthamiana* using split-luciferase complementation^54^. In this assay, if the protein dimerises, it will bring the split luciferase components together, producing luminescence. Luminescence was detected for wild type and all cysteine mutants, confirming dimerisation *in planta* (Supplementary Fig. S38). Signals were lower for cysteine mutants than wild type, consistent with reduced dimerisation, though differences among mutants were less pronounced than *in vitro*, possibly due to other proteins or factors stabilising dimers. These results support our conclusion that Cys60 and Cys165 mediate ACO dimerisation *in vivo*.

## Discussion

Ethylene biosynthesis is a central process in plant growth, development, and stress adaptation^8–11^. Yet, despite decades of study, the molecular regulation of its terminal enzyme, ACC oxidase (ACO), has remained elusive. Here, we provide structural and mechanistic insights into two distinct regulatory layers: induced-fit catalysis and redox-controlled oligomerisation.

### Substrate binding and bicarbonate anchoring

Our ACC-bound PhACO1 structure reveals bidentate chelation at the active-site metal, adopting an equatorial pose distinct from inhibitor-bound AtACO2 and prior models^19, 14^. Because three coordination sites are available on Fe(II), alternative ACC orientations are plausible. The requirement for bicarbonate supports a model in which the anion anchors ACC to the conserved RXS motif, mimicking carboxylate engagement seen in 2OG binding to 2OGDs^13–16^, thereby positioning the substrate for oxidation. Our structure supports this model and provides a molecular basis for bicarbonate’s role in substrate positioning and reactivity.

### Induced fit and catalytic activation

ACO undergoes a ligand-induced conformational change essential for catalysis. Ligand binding reorganises a network spanning the active site (ligand– RXS–β-sheet) and α3, bending the helix and closing the enzyme. NMR, fluorescence, SAXS and MD converge on this open-to-closed transition, and mutagenesis (R244A, E87D, E87Q) demonstrates that disrupting the network abolishes closure and reduces activity ≥80%, establishing induced fit as a prerequisite for catalysis and resolving previous uncertainties^13, 19, 20^. This closure locks in an optimised geometry for ligand binding, oxygen activation, and reactive intermediates (e.g., Fe-oxo species^14^) consistent with catalytic demands and situates ACO among 2OGDs that employ ligand-induced rearrangements^21–25^.

### A redox gate through disulfide-linked dimerisation

In addition to conformational dynamics, we uncovered a second regulatory layer: redox-controlled oligomerisation. Several ACOs, including PhACO1, have been reported to exist as mixtures of oligomeric states^13, 16, 19^. PhACO1 forms disulfide-linked dimers via Cys165 (primary) and Cys60 (secondary). The dimer is inactive and stabilises an open, rigid state that restricts α3 dynamics; reducing agents restore monomer and activity, whereas oxidants promote dimer formation. More importantly, our data are consistent with *in planta* observations showing that cellular redox environments are tightly coupled to ethylene biosynthesis and signalling^55–57^. In plant lysate, ACO is modified by glutathione or cysteinyl-glycine at Cys165 and remains active, indicating that glutathionylation locks ACO in its monomeric form under physiological reducing conditions. This novel regulatory mechanism highlights an interplay between dimerisation and thiol modification reveals a direct molecular link between redox state and ethylene control in plants^58,59^.

### Physiological integration and implications

Taken together, these findings position ACO as a redox-responsive sensor: active as a monomer in the closed, ligand-stabilised state, and shifted toward inactive dimer under oxidative conditions. This dual control – induced fit for catalysis and redox-mediated dimerisation for activity gating – offers a framework connecting ACO enzyme structure and function with stress-induced modulation of ROS, ascorbate and glutathione signalling in plants, and ethylene production^60–63^.

The ability to precisely control ethylene formation is of enormous significance for plant biology, agriculture and biotechnology industries. Our work reveals the dynamic structural features of ACO and protein-level regulatory mechanisms and suggest practical routes to tune ethylene output through chemical modulators that stabilise or block monomeric closure, and manipulation of redox pathways in agronomic settings^64,65^. These efforts will deepen our understanding of how plants integrate metabolic and oxidative signals to fine-tune hormone biosynthesis and lead to advances in crop resilience and postharvest quality.

## Methods

### Materials

Unless otherwise specified, chemicals were obtained from Sigma-Aldrich/Merck, Thermo Fisher Scientific, AK Scientific, or Bio-Rad. Tris-D11 and D_2_O were purchased from Cambridge Isotope Laboratories or Cortecnet. Chemically competent *E. coli* cells were obtained from Agilent.

### Plasmid preparation

The plasmid pET-28a(+) containing PhACO1 was kindly provided by Dr Zhihong Zhang and Professor Christopher J. Schofield (University of Oxford). PhACO1 variants R244A, E87D and E87Q mutants were generated by site-directed mutagenesis and subcloned into the pNIC28-Bsa4 expression vector. Sequences were verified by in-house sequencing services.

DNA constructs (gBlocks) encoding PhACO1 variants C60S, C165S, C60/165S and AtACO2 and the W89I AtACO2 mutant were synthesised by Integrated DNA Technologies and subcloned into the pNIC28-Bsa4 expression vector^66^.

### Recombinant protein production

Expression plasmids were transformed into *E. coli* BL21 (DE3) competent cells and grown in selective medium (2YT for PhACO1, terrific broth for AtACO2) supplemented with 50 µg/ml kanamycin at 37 °C with shaking. For the production of ^2^H,^15^N-labelled protein, cells were cultured in M9 minimal medium supplemented with ^15^N ammonium chloride (1 g/L) and non-labelled glucose (10 g/L) in D_2_O. At an OD₆₀₀ of 0.6–0.8, protein expression was induced with 0.5 mM isopropyl β-D-1-thiogalactopyranoside (IPTG). Cultures were incubated overnight at 28 °C (PhACO1) or 18 °C (AtACO2). Cells were harvested by centrifugation (30,000 × g, 20 min, 4 °C), resuspended in lysis buffer (50 mM HEPES (pH 7.8), 500 mM NaCl, 5 mM imidazole) and lysed by sonication on ice.

Proteins were purified using immobilised metal affinity chromatography (IMAC) and eluted isocratically with an elution buffer (50 mM HEPES pH 7.8, 500 mM NaCl, 500 mM imidazole). To remove any bound metal ions, IMAC eluates were diluted into 200 mM EDTA containing 15 mM ammonium acetate (pH 7.8) to a protein concentration of ∼1 mg/ml and incubated for 2 hours at 4 °C. Samples were concentrated, desalted using a PD-10 desalting column (Cytiva), and subsequently subjected to size exclusion chromatography (SEC) with isocratic elution in storage buffer (50 mM Tris-HCl pH7.5, 150 mM NaCl). To obtain pure monomer, proteins were either pre-incubated with 2 mM TCEP prior to SEC chromatography or selective monomer fractions were collected from SEC chromatography. Final protein samples were concentrated, flash-cooled in liquid nitrogen, and stored at -80 °C.

### Crystallography

Apo-PhACO1 (20 mg/mL) was incubated with 2 mM NiCl_2_, 1 mM dithiothreitol (DTT), 1 M NaCl in 50 mM Tris (pH 7.5) and 10% glycerol. 1 µL of the protein was mixed with the reservoir solution (1.3 M NaH_2_PO_4_, 0.7M K_2_HPO_4_, 0.3M Li_2_SO_4_, 0.1 M CAPS (pH 10.5)) at a 1:1 volume ratio in a hanging drop vapor diffusion experiment using a siliconised glass cover slip. Irregular shaped and stacked crystals were observed after ∼2 days. Further growth of the crystals was monitored over 2 weeks at 18 °C. Crystals were then picked and transferred to a micro-tube containing fresh reservoir (50 μL) and a seed bead (Hampton Research). The crystals were crushed by vortex mixing (∼2-3 mins) and resting on ice (∼2-3 mins). The resultant seed stock was then used to seed pre-equilibrated (overnight) drops. The trays were then incubated at 18 °C for 2 weeks. The crystals from the microseeding were then soaked in a cryo-solution (crystallisation buffer supplemented with 20% glycerol and 50 mM ACC) for 2 mins before mounting in nylon loops for flash-cooling in liquid nitrogen.

Diffraction data were collected at the Australia Synchrotron (AS) using beamline MX-1. Details of the beamline set up are available from the Australian Synchrotron website, and data were collected using the *Blu-Ice* software^67^.

The diffraction data were indexed and integrated using the auto processing software XDS^68^ while MOSFLM^69^, was used to index and integrate the data. The data was then merged and scaled using AIMLESS^70^. The structures were solved by molecular replacement using PHASER MR and the crystal structure of *hybrida* ACO1 (PDB ID: 1WA6) ^71^. Initial coordinates from molecular replacement were refined using the REFMAC software package^72^; coordinates and electron density maps were subsequently inspected using COOT modelling software^73^. Residual electron density was modelled as ACC, Ni(II) ions, and water molecules as appropriate and structure iteratively refined using either REFMAC or PHENIX^74^. Preliminary coordinates were visualised and figures produced using PyMol^75^. Data collection and refinement statistics table are provided in the Supplementary Information (Supplementary Tables S1 and S2)

### Protein-protein docking and electrostatic analysis

Docking was performed using PatchDock^76^. Structural models were obtained from the Protein Data Bank (PDB) and the AlphaFold Protein Structure Database: PhACO1 (PDB: 5TCV) and PhACO1 (AF-Q08506-F1-v4). For the PDB entry, missing loops and heavy atoms were rebuilt and hydrogen atoms added using PDBFixer^77^. For both structures, flexible N- and C-terminal regions were removed, and residues 3–308 of PhACO1 were used in docking. Docking was performed with default parameters, enforcing a <4 Å distance constraint between the thiol groups of Cys165 from each monomer. Resulting dimer models were clustered with an RMSD cutoff of 2 Å. The lowest-energy dimer was selected and analysed by the Adaptive Poisson–Boltzmann Solver (APBS) algorithm^78^. Electrostatic potentials were calculated for the monomer and dimer, and the monomer potential was subtracted from the dimer to visualize electrostatic changes upon dimerization.

### Molecular dynamics simulations

Atomistic MD simulations were performed using GROMACS version 2021^79^. In the monomeric state, two sets of simulations were performed: (i) Fe(II)-bound AtACO2 and (ii) AtACO2 in complex with Fe(II) and POA, both starting from the ligand-bound AtACO2 structure (PDB: 5GJ9)^19^. To model the dimeric form, MD simulations were performed using the PhACO1 dimer from docking(Template: PDB: 5TCV) as the starting conformation.

All systems were simulated using the CHARMM36m force field combined with the TIP3P water model^80^. After generating the molecular topologies, each system was placed in a cubic box, ensuring a minimum distance of 1.0 nm between the protein and the box boundary, and subsequently solvated with water. Na⁺ and Cl⁻ ions were added to neutralize the system and reach an ionic strength of 0.165 M. Newton’s equations of motion were integrated using the Verlet algorithm with a timestep of 2fs. A cutoff scheme was applied to compute the van der Waals and short-range Coulomb interactions, employing a pair-list cutoff of 1.0 nm. Long-range electrostatics beyond this cutoff were calculated with the Particle Mesh Ewald (PME) method^81^, using fourth-order (cubic) interpolation and a grid spacing of 0.16 nm. Energy and pressure corrections were applied to treat long-range dispersion forces, while bond lengths were fixed using the LINCS algorithm^82^.

To resolve atomic clashes, energy minimization was performed using the steepest descent algorithm. Equilibration of the solvent around the protein was achieved in two steps. First, the system was simulated for 500ps in the NVT ensemble, where a velocity-rescaling thermostat maintained the temperature at 300 K^83^, with velocity rescaling applied every 0.1 ps. In the second step, the velocities from the NVT step were used to run a 1 ns equilibration in the NPT ensemble, with pressure held isotropically at 1 bar using the Parrinello-Rahman barostat^84^. Throughout both equilibration steps, protein atoms were positionally restrained using a force constant of 1000 kJ mol^-1^ nm^-1^. Production simulations were then performed in the NPT ensemble.

Each system was simulated for a total of 2 μs (5 replicates of 400 ns each). Trajectories were collected every 10 ps, and the first 50 ns of each replicate was discarded as equilibration time. All analyses were performed either using tools available in the GROMACS suite, *in-house* scripts or MDAnalysis^85,86^. All distances were calculated considering the minimum image convention. The principal component analysis (PCA) in coordinate space, in the dimer complex was carried out using GROMACS. The covariance matrix of the atomic positional covariance was constructed with *g_covar*, and the corresponding eigenvectors and eigenvalues were obtained. Projections of the trajectories onto the eigenvectors were generated with *g_anaeig*. To visualize the dominant motions, porcupine plots were created in VMD^87^.

### NMR-based ACO activity assay

ACO activity was monitored using a Bruker Avance III HD spectrometer equipped with either a room-temperature BBFO probe or a Prodigy cryoprobe. Reactions were prepared in 5 mm NMR tubes (550 µl, final volume) and analysed at 500 MHz and 298 K.

For time course measurements, all protein samples are in monomeric state unless stated. Reactions contained 30 mM NaHCO₃, 12.5 mM ascorbic acid, 250 µg catalase, 250 µM ACC and 50 mM Tris-D11 (pH 7.5) in 90% H_2_O/10% D_2_O. Blank spectra (without enzyme and Fe(II)) were acquired prior to initiating reactions. Reactions were initiated by addition of 2 µM ACO and 20 µM Fe(II)SO_4_ (freshly prepared from 250 mM stock in 20 mM HCl). Standard proton with water suppression by excitation sculpting were collected starting 2 min 40 sec after mixing. Each assay was repeated at least three times. For endpoint assays, identical mixtures were prepared, but a single spectrum was collected after 2 hours. Enzymatic activity was determined by comparison with blanks.

### Protein NMR-based structural conformational change assay

NMR experiments were performed at 600 MHz on a Bruker Avance spectrometer equipped with an inverse TCI cryoprobe. Samples (550 µl) were loaded into 3-mm MATCH NMR tubes. Experiments were conducted at 298 K using the standard ^1^H-^15^N TROSY pulse sequence. Samples contained 100 µM ^2^H,^15^N-labelled enzyme and 100 µM Ni(II)SO₄, with or without 200 µM POA, in 50 mM Tris-D11 buffer (pH 6.6) in 90% H_2_O and 10% D_2_O.

### Intrinsic tryptophan fluorescence quenching assay

Assays were prepared with 5 µM enzyme, 50 µM NiCl₂, and ligand as indicated (ACC, POA) in 50 mM Tris (pH 7.5). Reactions were performed in triplicate in 96-well black microplates (Eppendorf, 96/UPP). Fluorescence was measured at 28 °C on a PerkinElmer EnSpire multimode plate reader (excitation 295 nm; emission 315–450 nm). Maximum fluorescence intensity at λ_max_ = 350 nm was used for comparison.

### Thermal shift assay

Protein samples (20 µM) and metal ions (100 µM) were prepared in 25 mM Tris (pH 7.5) supplemented with 1 mM DTT and 10% glycerol. The samples were first incubated at 4 °C for 30–40 min. Sypro Orange (1:25 dilution) was then added, and reactions were loaded into 96-well plates. Thermal denaturation was monitored on a Bio-Rad MyiQ real-time PCR system (ver. 2) with a temperature gradient of 25–95°C at 1°C/min.

### SEC-MALS

Experiments were performed on a Wyatt SEC-MALS system comprising a DAWN multi-angle light scattering (MALS) detector and an Optilab refractive index detector, coupled to a Shimadzu HPLC system (SPD-20A UV/Vis detector, FRC-10A fraction collector, CBM-20A communication bus module, CTO-20A column oven, DGU-20A5R degasser, LC-20AD pump, SIL-20ACHT autosampler) equipped with a Superdex 200 Increase 5/150 GL column (Cytiva).

SEC-MALS characterization was carried out at 25 °C with a flow rate of 0.3 mL/min. The column was pre-equilibrated with SEC buffer (50 mM Tris-HCl pH 7.5, 150 mM NaCl) until stable baselines were achieved for UV, MLS, and DLS detectors.

Protein samples (2 µL at 4 mg/mL in SEC buffer) were injected. LabSolutions software (Shimadzu) was used to control the HPLC system, and Astra 7.3.2 software (Wyatt) was used for the MALS system. Blank buffer injections were performed to assess carryover between runs. Data were processed in Astra 7.3.2, and molecular weights were determined using the first-order Zimm fit method, with a refractive index increment (dn/dc) of 0.184 mL/g for proteins.

### Small-angle X-ray scattering (SAXS)

Small-angle X-ray scattering (SAXS) experiments were performed at the Australian Synchrotron SAXS/WAXS Beamline using a coflow setup to minimize radiation damage and allow for higher X-ray flux (11,500 eV)^88^. An in-line SEC system was employed to reduce protein sample dilution^89^. The resulting SAXS data were processed with Scatterbrain^90^ and further analysed using CHROMIXS^91^ and the ATSAS 3.2.1 software suite^92^. Protein samples were freshly prepared by incubating PhACO1 (260 µM) with NiSO_4_ (1.3 mM), in the presence or absence of 1 mM POA, in SEC buffer (50 mM Tris-HCl, pH 7.5, 150 mM NaCl). Full experimental procedures and analysis details are provided in the Supplementary Table S4.

### Intact protein mass spectrometry (LC–MS)

LC–MS analysis was carried out on an Agilent 6545XT QTOF with a PLRP-S column (100 Å, 1.0 × 50 mm, 5 µm). Samples (0.5 µg) were injected at 200 µl/min. Solvent A: 0.1% formic acid in water; Solvent B: 95% acetonitrile/0.1% formic acid. Gradient: 0–3 min, 10% B; 3–7 min, 10–70% B; 7–7.5 min, 70% B; 7.5–8 min, 70–10% B; 8–10 min, 10% B. Between injections, 1 µL trifluoroethanol was used for washing. Data were acquired in positive ESI mode (4 kV, m/z 100– 3200, 2 spectra/s). Raw spectra were analysed with MassHunter Qualitative Analysis; isotopic distributions were deconvoluted over 30–100 kDa at 1 Da increments using proton adducts.

### Analytical size-exclusion chromatography for dimer characterization

Samples (25 µL) containing 250 µM apo-PhACO1, 2.5 mM NiSO_4_, and 500 µM H₂O₂ in 50 mM Tris-HCl (pH 7.5), 150 mM NaCl were incubated on ice overnight. The samples were then diluted to 100 µl, centrifuged (10,000 rpm, 10 min, 4 °C), and analysed on a Superdex 200 Increase 10/300 GL column (Cytiva) using an ÄKTA go™ chromatography system.

### Plant materials and growth conditions

*Arabidopsis thaliana* ET-1 seeds, which contain CRISPR-generated deletions in all five *ACO* genes^50^ (kindly provided by Professor Li-Jia Qu, Cornell University) were verified by PCR and sequencing (Supplementary Table S3). Seeds were surface-sterilized in 30% (v/v) bleach/0.02% (v/v) Triton X-100, rinsed four times in sterile water, and sown on half-strength Murashige & Skoog (½ MS) medium containing 3 mM MES-KOH (pH 5.7) and 0.8% (w/v) Type M agar. Seeds were chilled for 2 days at 4 °C in the dark, then germinated under 12 h light (∼100 µmol m⁻^2^ s⁻¹)/12 h dark at 20 °C. After 7 days, seedlings were transferred to soil and cultivated under 16 h light/8 h dark at 21 °C. *Nicotiana benthamiana* plants were sown directly in soil and grown under a 12 h light/12 h dark cycle at 24/20 °C day/night and 70% relative humidity.

### Split luciferase assay

Coding sequences of PhACO1 and cysteine mutants were amplified by PCR from pET-28a(+) constructs, cloned into pCAMBIA1300-cLUC and -nLUC binary plant expression vectors^54^ and transformed into *Agrobacterium tumefaciens* AGL1. Cultures were grown in LB medium with 50 µg/ml kanamycin and 100 µg/ml ampicillin at 28 °C for 2 days, subcultured overnight, pelleted, and resuspended to OD₆₀₀ = 0.5 in infiltration buffer (10 mM MgCl₂, 1 mM MES (pH 5.6), 0.1 mM acetosyringone). After 2 h incubation in the dark, bacterial suspensions containing expression constructs (e.g. WT-cLUC + WT-nLUC) and a P19 helper plasmid for silencing suppression were combined at equal OD and co-infiltrated into leaves of 4-week-old *N. benthamiana* plants. Plants were incubated under a plastid hood in the dark for 48 h, sprayed with 1 mM D-luciferin K⁺ salt (Promega), and imaged after 5 min using a Retiga LUMO CCD camera (Teledyne Photometrics) and luminescence was quantified in transformed leaf regions using ImageJ.

### Plant lysate incubation

Leaves from 3–4-week-old *A. thaliana* ET-1 plants were snap-frozen, ground, and extracted with 50 mM potassium phosphate buffer (pH 7.4) supplemented with EDTA-free protease inhibitor (1:2 w/v). Lysates were incubated on ice for 30 min and centrifuged (12,000 × g, 10 min, 4 °C). Recombinant apo-PhACO1 or cysteine mutant proteins (500 µg) were incubated with 50 µL plant lysate overnight at 4 °C. Where indicated, 2 mM H₂O₂ was added. Incubated protein samples were first purified using His GraviTrap column (Cytiva), then analysed on a Superdex 200 Increase 10/300 GL column (Cytiva) using an ÄKTA go™ chromatography system.

### Native PAGE

Purified protein samples were adjusted to 2 mg/mL in 50 mM Tris-HCl (pH 7.5) and mixed with non-denaturing loading dye (50 mM Tris-HCl pH 6.8, 40% glycerol [v/v], 0.01 mg/mL bromophenol blue) at a 1:1 ratio. Samples were separated by 10% native polyacrylamide gel electrophoresis at 120 V for 100 min at 4 °C in running buffer (25 mM Tris base, 192 mM glycine; pH 8.8). Gels were stained with Coomassie Brilliant Blue for protein visualisation.

### Protein sequence alignment

ACO sequences (n = 70) from ten plant species (*Arabidopsis thaliana, Malus domestica, Petunia hybrida, Solanum lycopersicum, Nicotiana benthamiana, Glycine max, Triticum aestivum, Zea mays, Amborella trichopoda, Marchantia polymorpha*) were obtained by searching for “ACC oxidase” in the NCBI gene database for each species. Information for each gene is provided in Supplementary Table S5. Sequence alignments were performed with MEGA11 (default parameters)^93^. Sequence conservation around residues Cys60 and Cys165 was analysed with WebLogo^94^.

### Cysteine post translational modification

To ensure protein modification, 2 µM protein was incubated in SEC buffer with freshly prepared 2 mM TCEP, 2 mM GSH, 2 mM GSSH, or 2 mM Cys–Gly (each dissolved in H_2_O) on ice for 45 min prior to further analysis.

### Tandem mass spectrometry

A 0.5 µL aliquot of protein (dimer fraction collected from the size exclusion chromatograph) was diluted in 50 mM ammonium bicarbonate containing 50 mM iodoacetamide, incubated in the dark for 30 minutes, then spun in a Vivaspin concentrator to remove unreacted iodoacetamide before performing a 1 hr microwave digest with sequencing grade trypsin at 45 °C. The resulting digest was then diluted with 0.1% formic acid for LC-MS/MS analysis on a TripleTOF 6600 mass spectrometer. MS/MS spectra were then manually interrogated for fragment ion masses derived from the expected tryptic peptides containing Cys60 and Cys165, using PeakView software, and crosslinked fragment ions were then manually annotated.

### Ligand binding assay by NMR

The binding of POA to the PhACO1 mutants was assessed using a 500 MHz Bruker Avance NMR spectrometer equipped with a cryoprobe. Samples (550 µL) were prepared in standard 5-mm NMR tubes and measured at 298 K. WaterLOGSY experiments were performed using the pulse program *ephogsygpno*, as described by Dalvit et al.^95^, with 512 scans. Water suppression was achieved using excitation sculpting^96^. Each sample contained 400 µM POA and 10 µM ZnCl_2_, with or without 10 µM enzyme, in 50 mM Tris-d_11_ buffer (pH 7.5) prepared in 90% H_2_O and 10% D_2_O.

### ITC assays

The dissociation constant (*K*_D_) of POA binding to PhACO1 and the R244A mutant was determined using isothermal titration calorimetry (ITC) on a MicroCal VP-ITC instrument (Malvern Panalytical). Experiments were performed at 10 °C with a reference power of 8 µcal s⁻¹. Prior to titration, protein samples were extensively buffer-exchanged and diluted to 30 µM in 50 mM Tris-HCl (pH 7.5) containing 100 µM NiCl_2_. The POA stock solution was freshly prepared in the same buffer to minimise buffer mismatch between the syringe and cell solutions.

## Supporting information

Electronic Supplemental Information

## Acknowledgements

We acknowledge the use of the Nuclear Magnetic Resonance Centre and Mass Spectrometry Hub at the University of Auckland, and Melbourne Magnetic Resonance Platform, Melbourne Protein Characterisation Platform, and Mass Spectrometry and Proteomics Facility at the University of Melbourne for this work. The plasmid encoding wildtype PhACO1 was a kind gift from Dr Zhihong Zhang and Professor Christopher J. Schofield (University of Oxford). The *A. thaliana* ET-free seeds were a kind gift from Professor Li-Jia Qu (Peking University). This research was undertaken in part using the MX2 beamline at the Australian Synchrotron, part of ANSTO, and made use of the Australian Cancer Research Foundation (ACRF) detector. D.M.G. and J.A.P.M. were supported by the University of Auckland Doctoral Scholarship. F.K. was supported by the Melbourne Research Scholarship, Dr Jim Desmarchelier Scholarship and the Norma Hilda Schuster (nee Swift) Scholarship. X.C. was supported by the Melbourne Research Scholarship and Rowden White Scholarship. I.K.H.L. acknowledge the University of Auckland, the University of Melbourne and the Royal Society Te Apārangi Marsden Fund (18-UOA-040) for financial support of this project.

## Author contributions

D.M.G., F.K. and S.K. produced and purified proteins. D.M.G., F.K., S.K. and X.C. performed *in vitro* assays. D.M.G. performed crystallography with the help of Y.Y. and C.J.S. F.K. and J.A.P.M. performed docking and molecular dynamics simulations with the help and supervision of D.M. F.K. performed plant-based experiments with supervision of M.J.H. D.M.G., H.Y.H.T., F.K. and A.S. performed SAXS experiments. F.K. and M.J.M. performed protein mass spectrometry experiments. M.J.H., C.J.S. and I.K.H.L. designed the studies. All authors contributed to the writing and reviewing of the paper.

