## Supplementary material for "Redox-controlled dimerisation regulates ethylene biosynthesis": Electronic Supplemental Information

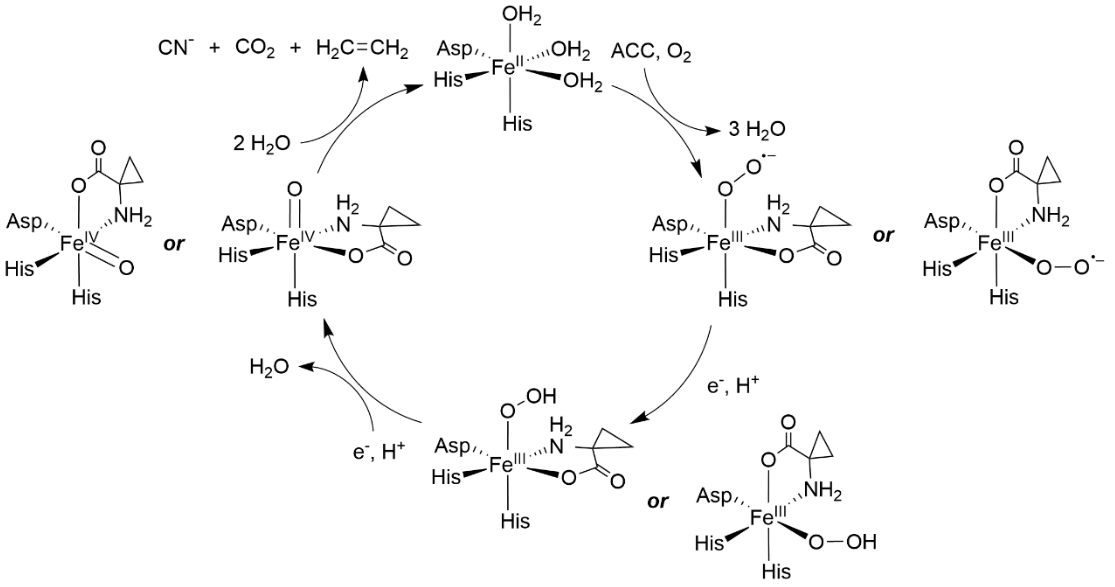

**Supplementary Figure S1 | Catalytic mechanism of ACC oxidase (ACO).** ACC binding displaces three water molecules from the Fe(II) active site, creating a vacant site for $O_2$. Coordination of ACC and $O_2$ to Fe(II) is shown based on current structural analysis; alternative positions ("or") reflect previous inhibitor-mimicking models. Oxidation of ACC is proposed to proceed via a reactive Fe(IV)=O intermediate.

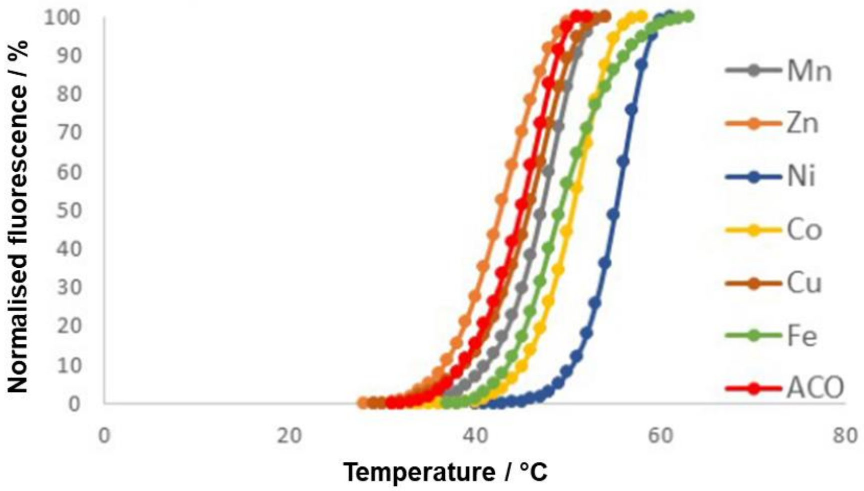

**Supplementary Figure S2 | Melting curves of PhACO1 in the presence of various transition metals.** The metal tested were $Mn^{2+}$, $Ni^{2+}$, $Zn^{2+}$, $Co^{2+}$, $Cu^{2+}$, $Fe^{2+}$ at a 1:5 protein-to-metal concentration ratio. Fluorescence intensity values were averaged and normalised for each dataset to ensure consistency and comparability across the different metal conditions.

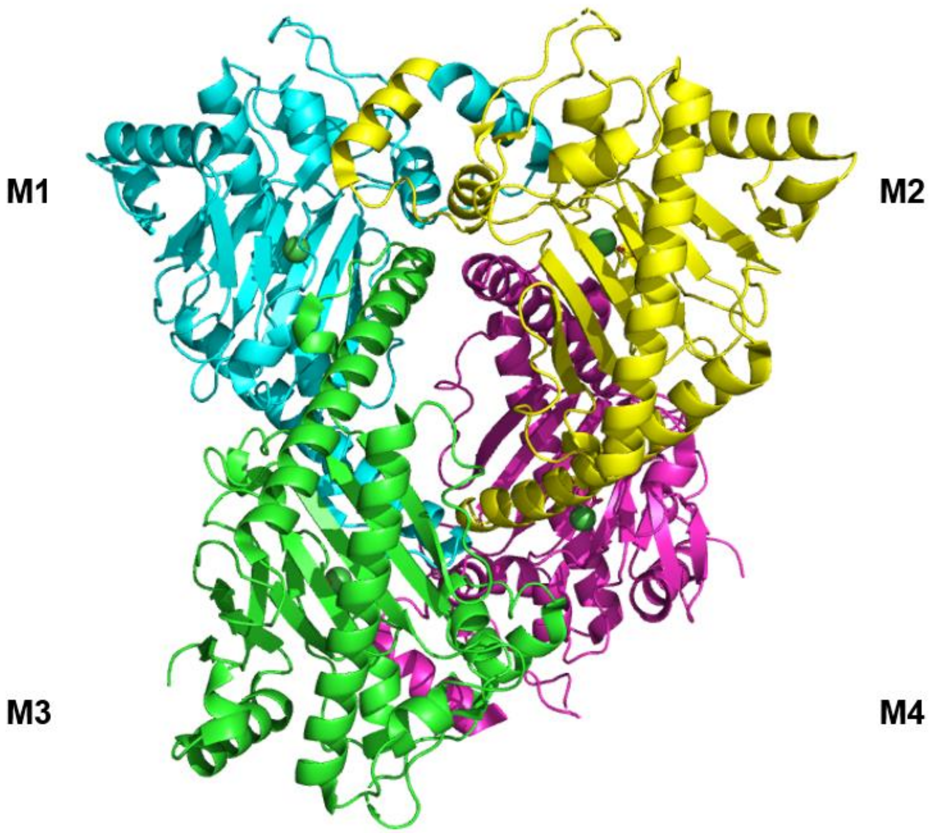

**Supplementary Figure S3 | Crystal structure of PhACO1 in complex with Ni(II) (PDB: 5TCW).** The protein was crystallised as an apparent tetramer.

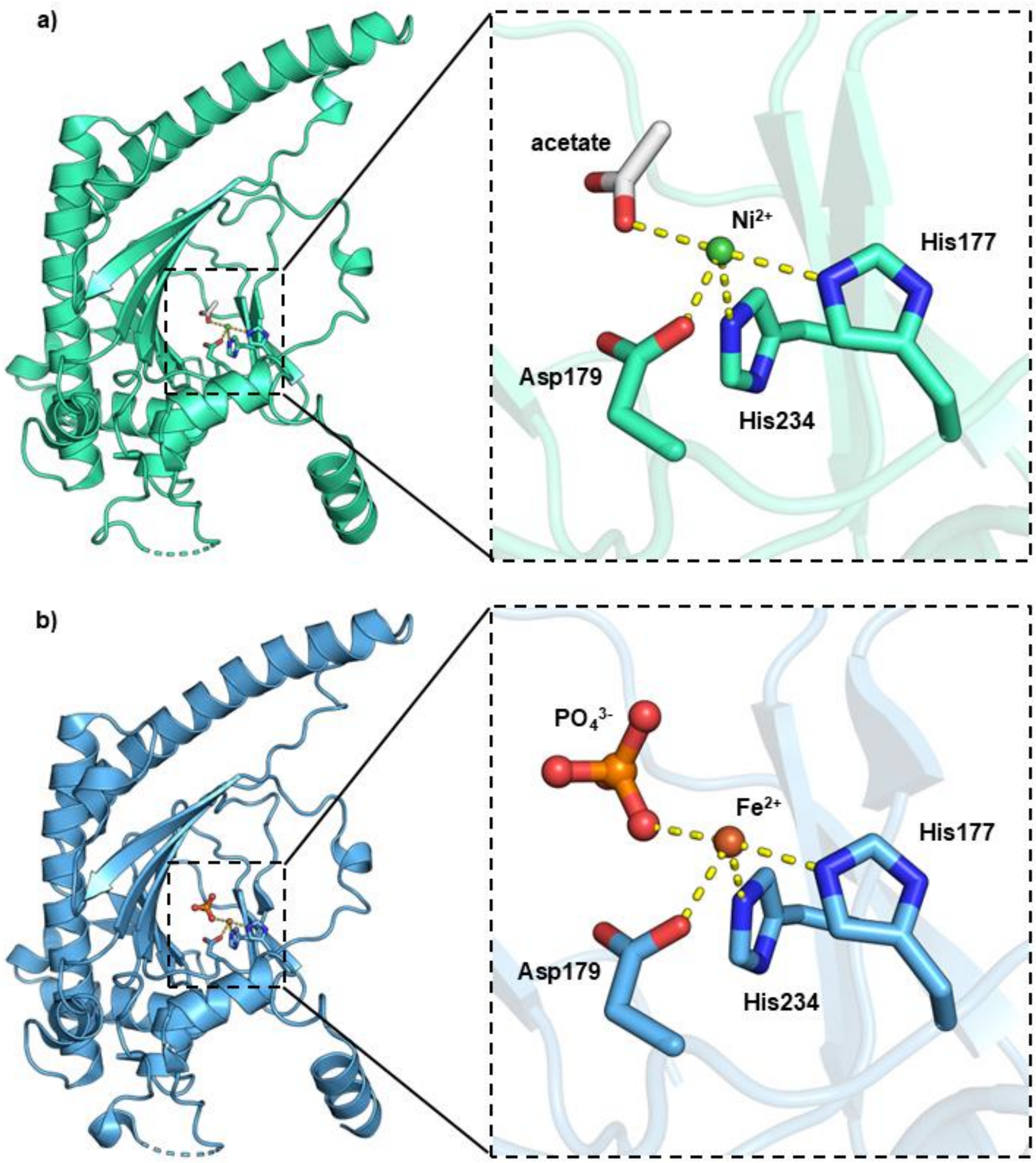

**Supplementary Figure S4 | Metal coordination in PhACO1.** (a) Ni(II) coordination in PhACO1 (PDB: 5TCW) involves His177, Asp179, and His234, forming the canonical “2-His-1-carboxylate” motif of 2OG oxygenases. An acetate occupies the site opposite His177. (b) Fe(II) coordination in PhACO1 (PDB: 1WA6; Zhang et al.) mirrors the Ni(II) arrangement, with a phosphate ion bound opposite His177.

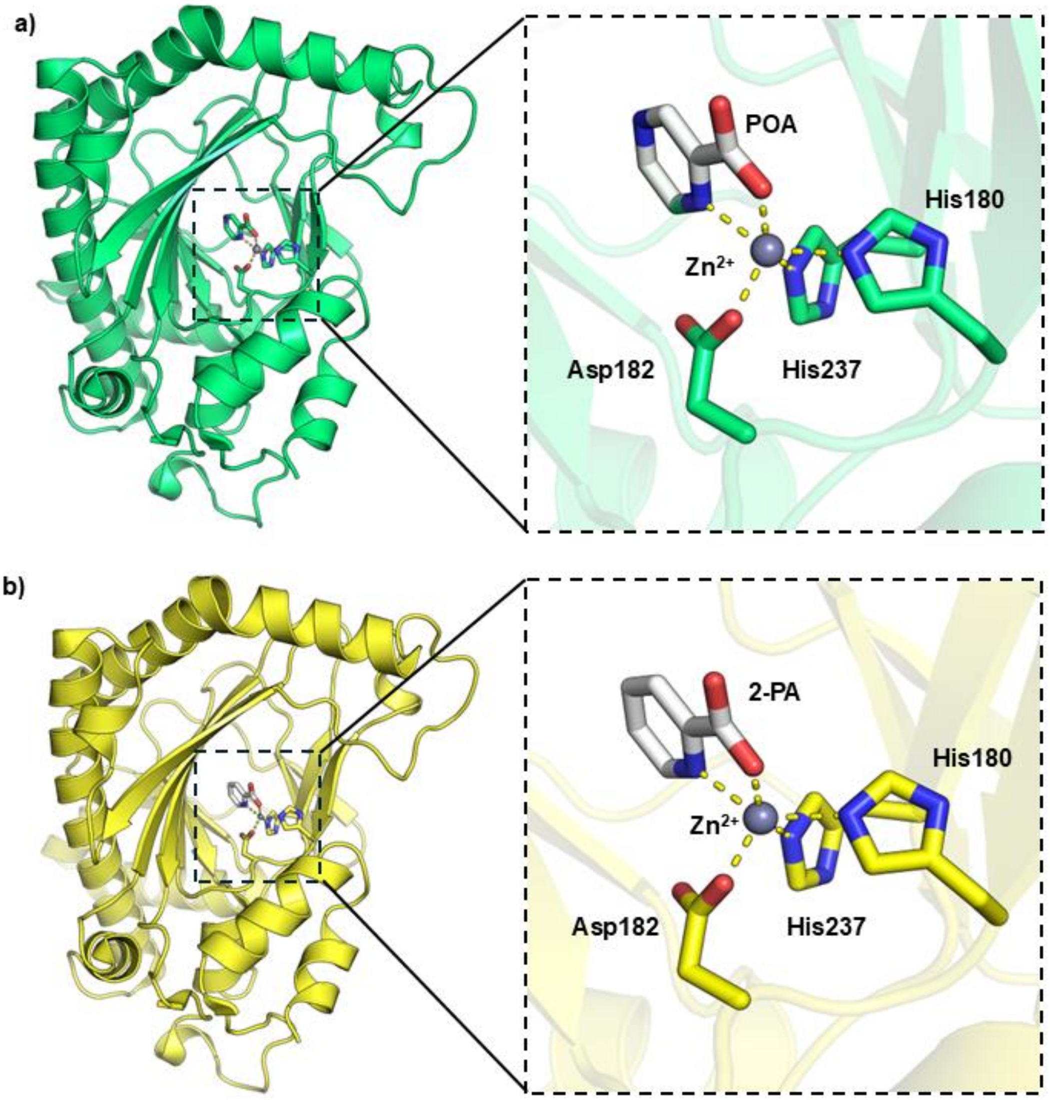

**Supplementary Figure S5 | Crystal structures of AtACO2 with Zn(II) and inhibitors.** (a) AtACO2 bound to Zn(II) and pyrazinecarboxylic acid (POA) (PDB: 5GJ9). (b) AtACO2 bound to Zn(II) and 2-picolinic acid (2-PA) (PDB: 5GJA). Both POA and 2-PA coordinate Zn(II) in a bidentate manner, with the nitrogen occupying the “acetate/phosphate” site.

| Length | Identity | Similarity | Gaps | Score |
|---|---|---|---|---|
| 325 | 220/325 (67.7%) | 267/325 (82.2%) | 11/325 (3.4%) | 1224 |

```
ACCO1_PETHY       1 ME---NFPIISLDKVNGVERAATMEMIKDACENWGFFELVNHGIPREVMD     47
                    ||   .||::.|.|:||.||..||.:|.:|||||||||:||||:|.::||
ACCO2_ARATH       1 MEKNMKFPVVDLSKLNGEERDQTMALINEACENWGFFEIVNHGLPHDLMD     50

ACCO1_PETHY      48 TVEKMTKGHYKKCMEQRFKELVASKALEGVQAEVTDMDWESTFFLKHLPI     97
                    .:|||||.|||.|.||:|.:::.||.|:.::.||.|:||||||:::|||.
ACCO2_ARATH      51 KIEKMTKDHYKTCQEQKFNDMLKSKGLDNLETEVEDVDWESTFYVRHLPQ    100

ACCO1_PETHY      98 SNISEVPDLDEEYREVMRDFAKRLEKLAEELLDLLCENLGLEKGYLKNAF    147
                    ||::::.|:.:|||..|:||.||||.|||:|||||||||||||||||..|
ACCO2_ARATH     101 SNLNDISDVSDEYRTAMKDFGKRLENLAEDLLDLLCENLGLEKGYLKKVF    150

ACCO1_PETHY     148 YGSKGPNFGTKVSNYPPCPKPDLIKGLRAHTDAGGIILLFQDDKVSGLQL    197
                    :|:|||.||||||||||||||::|||||||||||||||||||||||||||
ACCO2_ARATH     151 HGTKGPTFGTKVSNYPPCPKPEMIKGLRAHTDAGGIILLFQDDKVSGLQL    200

ACCO1_PETHY     198 LKDGQWIDVPPMRHSIVVNLGDQLEVITNGKYKSVMHRVIAQKDGARMSL    247
                    ||||.||||||:.||||:|||||||||||||||||:|||:.|::|.|||:
ACCO2_ARATH     201 LKDGDWIDVPPLNHSIVINLGDQLEVITNGKYKSVLHRVVTQQEGNRMSV    250

ACCO1_PETHY     248 ASFYNPGSDAVIYPAPALVEKEAEENKQVYPKFVFDDYMKLYAGLKFQAK    297
                    ||||||||||.|.||.:||||::|     ||.||||||||||||:|||.|
ACCO2_ARATH     251 ASFYNPGSDAEISPATSLVEKDSE-----YPSFVFDDYMKLYAGVKFQPK    295

ACCO1_PETHY     298 EPRFEAMKAMETDVKMDPIATV---    319
                    ||||.|||......:::|.|.|
ACCO2_ARATH     296 EPRFAAMKNASAVTELNPTAAVETF    320
```

**Supplementary Figure S6 | Protein sequence alignment of PhACO1 (ACCO1_PETHY) and AtACO2 (ACCO2_ARATH).** This figure was generated using Clustal Omega at EMBL-EBI with default parameters[1].

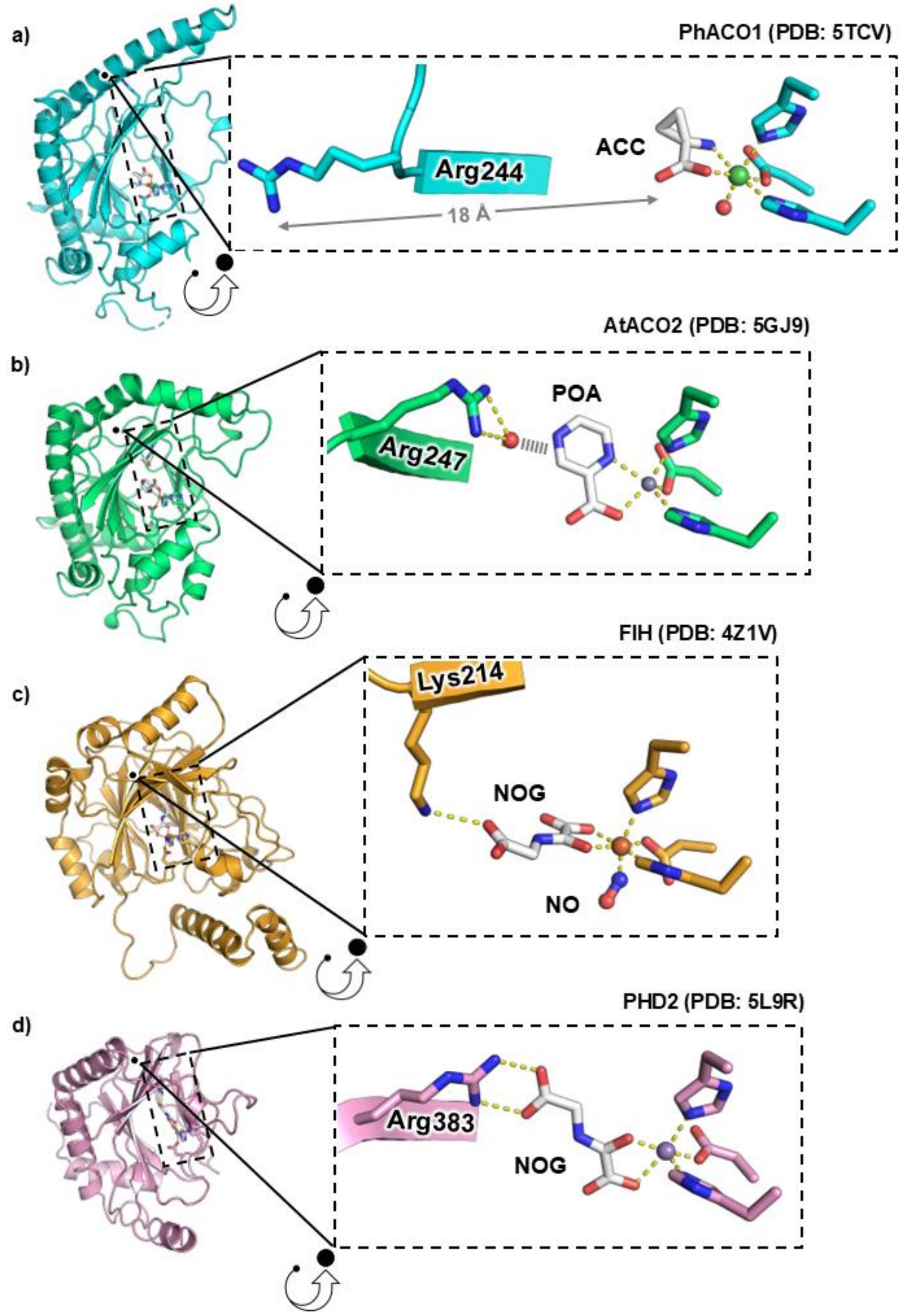

**Supplementary Figure S7 | Comparison of ACC and POA binding in ACO versus NOG binding in 2OG oxygenases.** (a) In PhACO1-Ni(II)-ACC (PDB: 5TCV), Arg244 points away from the metal centre. (b) In AtACO2-Zn(II)-POA (PDB: 5GJ9), Arg247 faces inward and coordinates a water molecule that weakly interacts with POA. (c, d) In the human 2OG oxygenases FIH (PDB: 4Z1V) and PHD2 (PDB: 5L9R), the basic residue (Lys214 or Arg383) interacts directly with the terminal carboxylate of NOG.

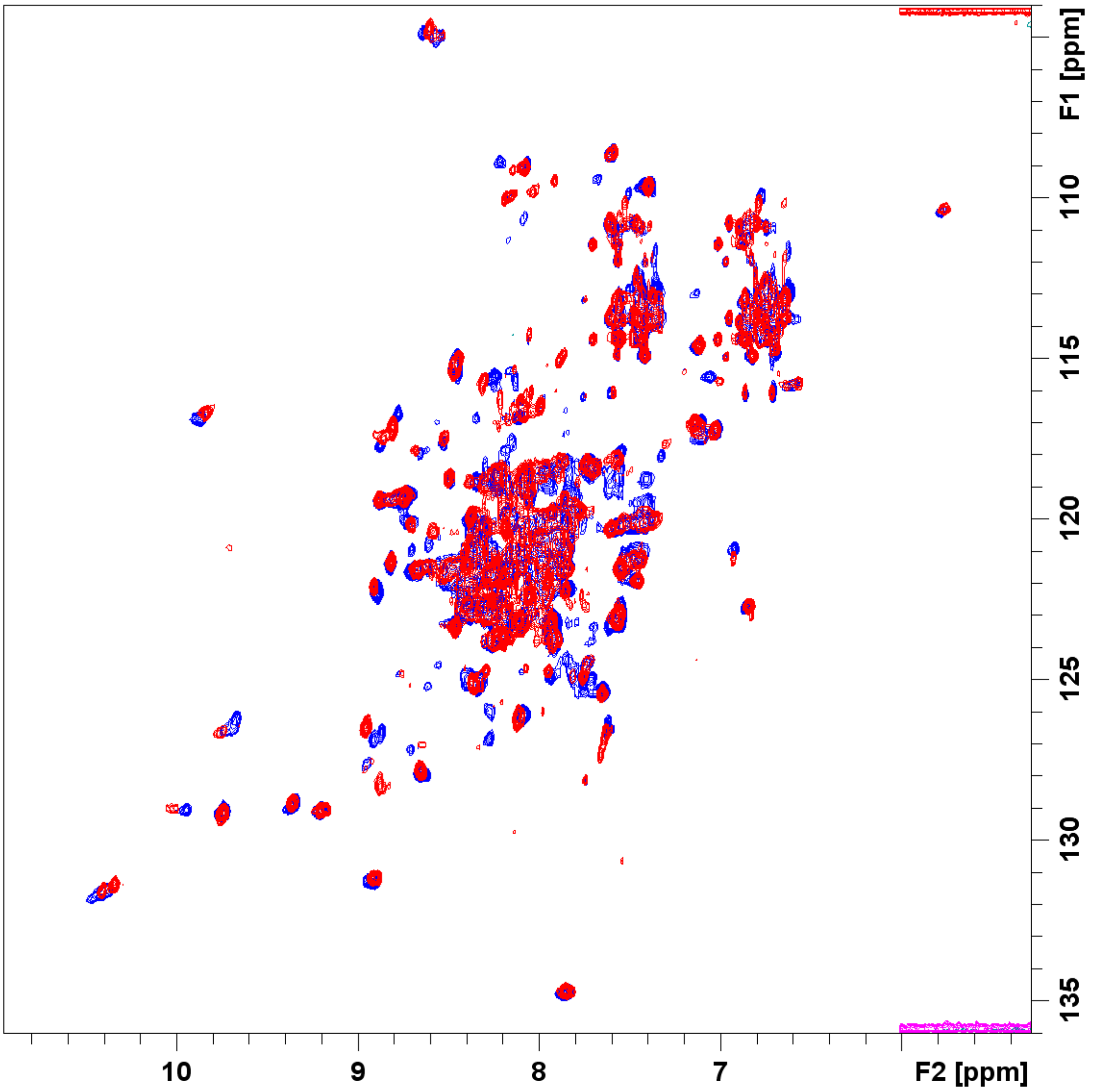

**Supplementary Figure S8 | $^{1}H$-$^{15}N$ TROSY spectra of AtACO2 bound to Ni(II) (blue) and Ni(II)–POA (red).** Samples comprised 100 µM $^{2}H$,$^{15}N$-labelled AtACO2, 150 µM Zn(II), 3 mM POA (where applicable) in 50 mM Tris-D11 (pH 6.6) with 90% $H_2O$/10% $D_2O$. Spectra were recorded at 298 K.

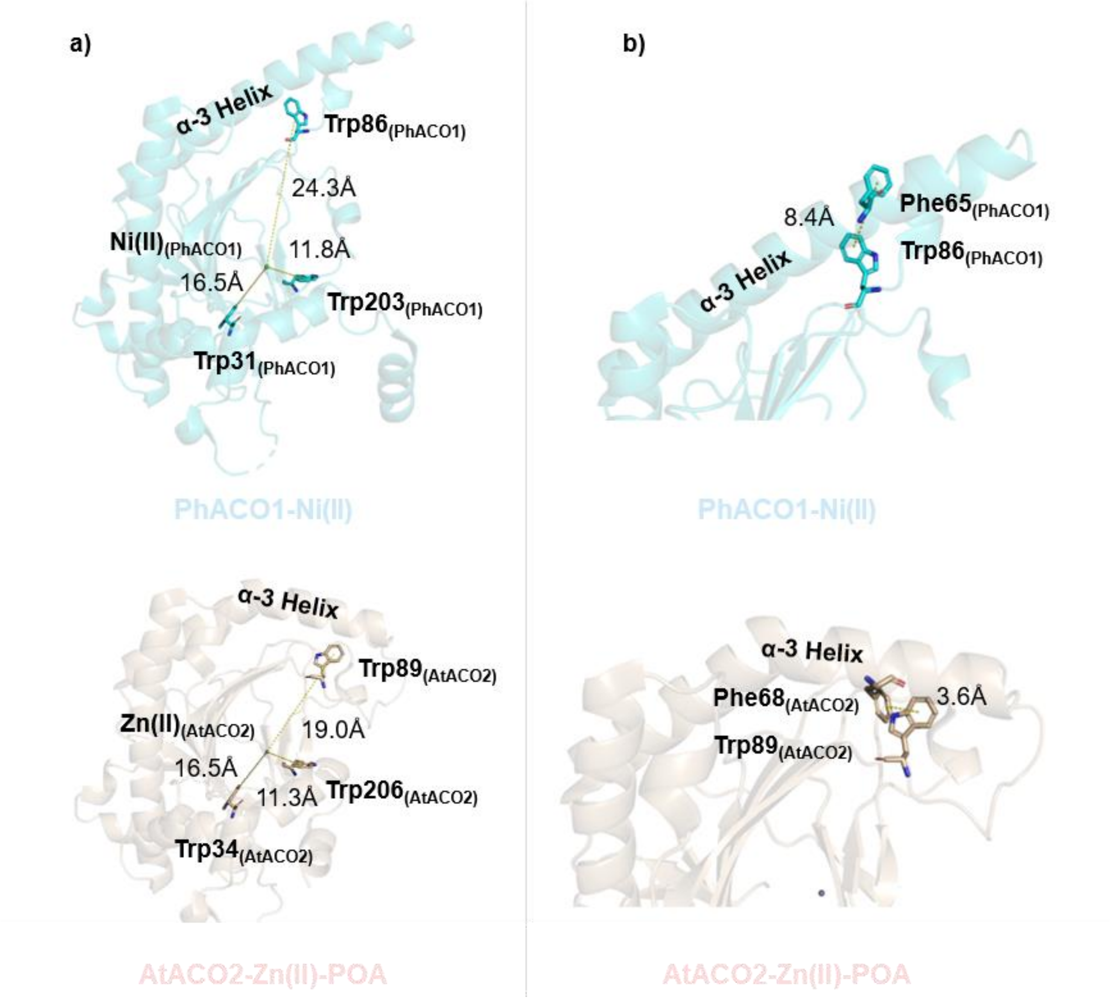

**Supplementary Figure S9 | Intrinsic tryptophan fluorescence as a probe for ACO conformational change.** Both PhACO1 and AtACO2 contain three tryptophan residues in their structures (PhACO1: Trp31, Trp86, Trp203; AtACO2: Trp34, Trp89, Trp207). (a) In both structures, these tryptophan residues are located ⩾10 Å away from the active site, so their fluorescence should not be directly affected by ligand binding. (b) In PhACO1-Ni(II) ("open" conformation), Trp86 is located 8.4 Å away from its nearest aromatic residue (Phe65). In AtACO2-Zn(II)-POA ("closed" conformation), Trp89 is only 3.6 Å from Phe68, suggesting potential fluorescence quenching.

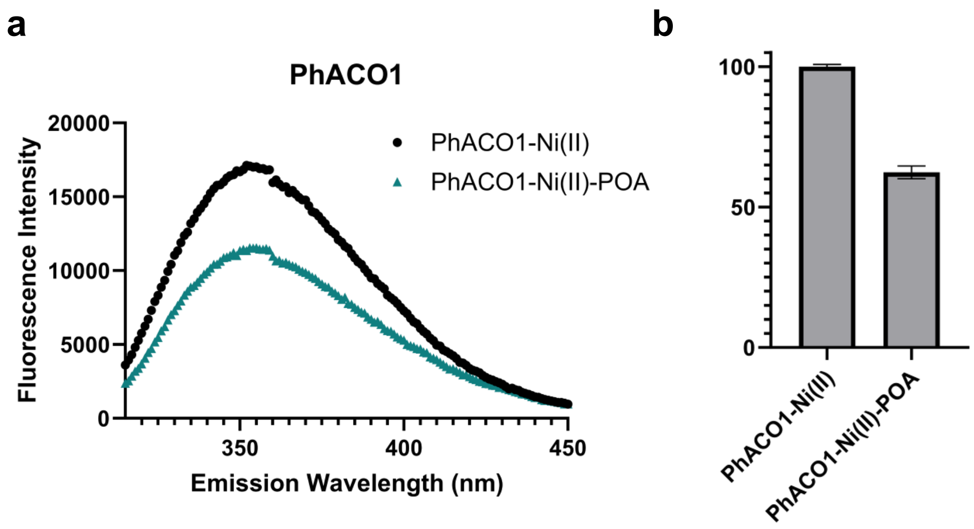

**Supplementary Figure S10 | Intrinsic fluorescence of PhACO1-Ni(II) with and without POA**. (a) Emission spectra PhACO1-Ni(II) alone (black) and with POA (blue). (b) Normalised fluorescence intensity for PhACO1-Ni(II) with and without POA. Samples comprised 2 µM PhACO1 and 20 µM Ni(II), with either 200 µM ACC or 200 µM POA, highlighting spectral changes upon ligand interaction.

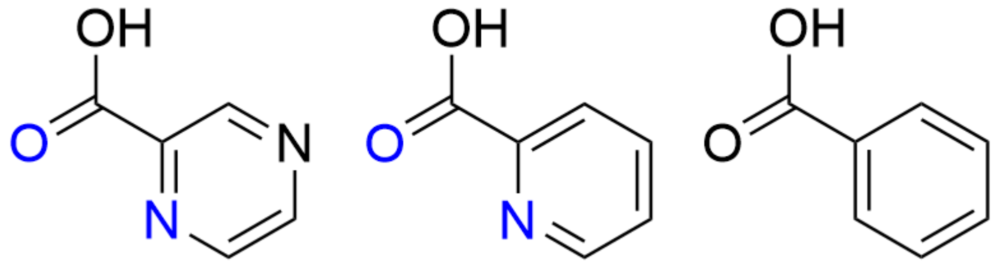

2-Pyrazinecarboxylic acid (POA) 2-Picolinic acid (2-PA) Benzoic Acid

**Supplementary Figure S11 | Chemical structures of POA, 2-PA and benzoic acid.** The atoms that chelate the metal ion within the enzyme complex are highlighted in blue. Both POA and 2-PA coordinate through the carbonyl group and pyridine nitrogen, whereas benzoic acid lacks the pyridine nitrogen and does not bind to ACO.

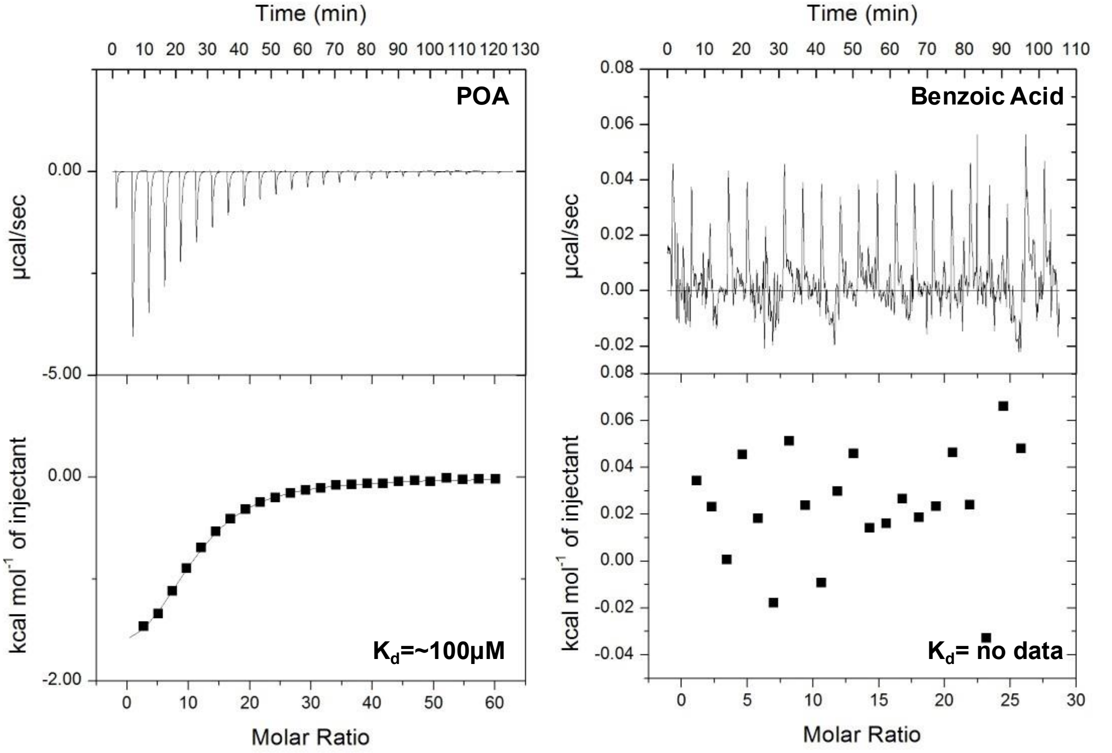

**Supplementary Figure S12 | ITC analysis of POA and benzoic acid binding to AtACO2-Ni(II).** Left: Raw ITC data for 25 injections of POA (first 2 µL, then subsequently 10 µL; 10 mM POA stock; 300 s interval) into a cell containing AtACO2 (30 µM) with 100 µM Ni(II). The integrated titration curve (baseline-corrected) indicates a $K_D$ of ~100 µM. Right: ITC data for benzoic acid under the same conditions. No binding was detected.

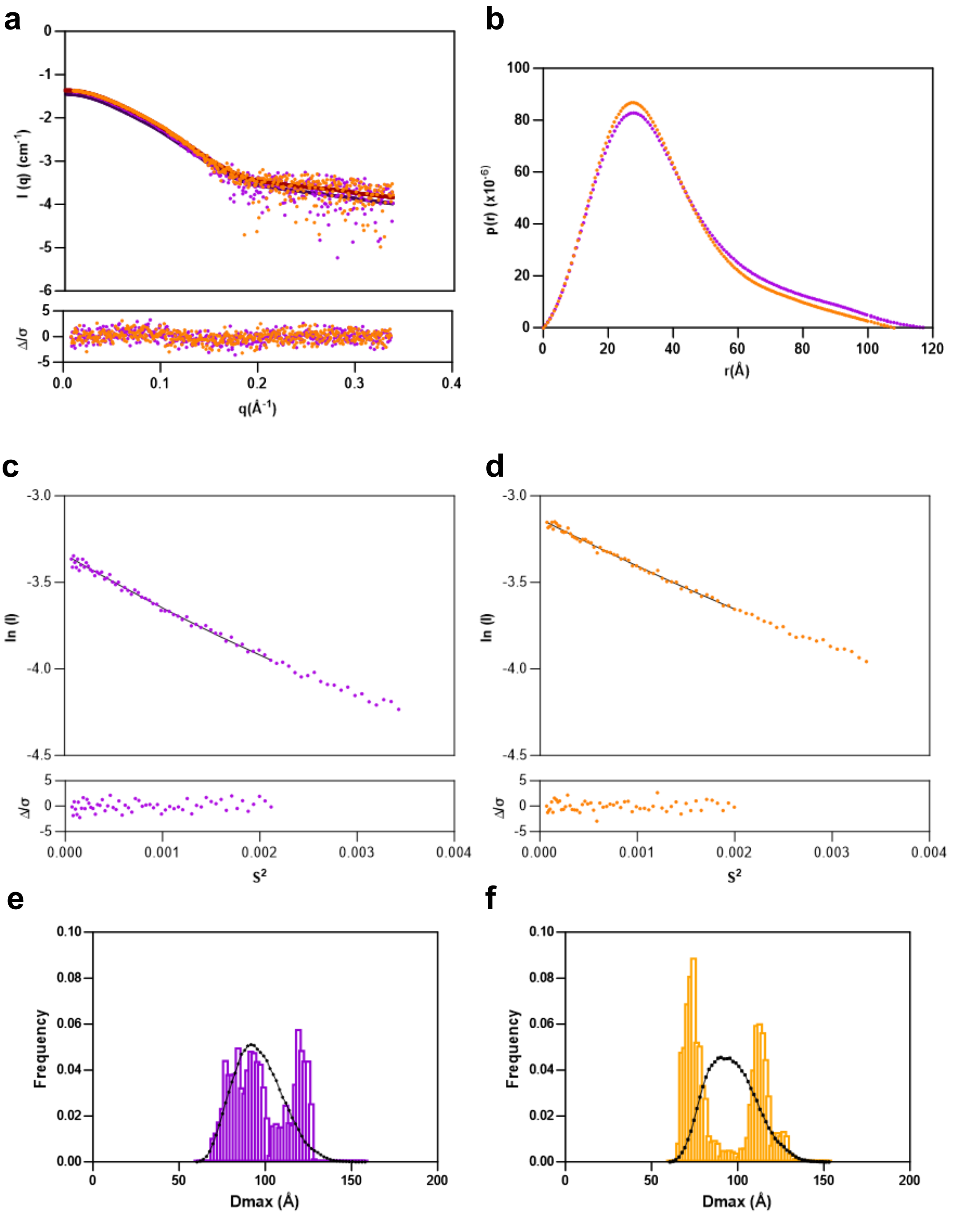

**Supplementary Figure S13 | SAXS analyses of POA-induced conformational changes.** (a) Experimental SAXS profiles for PhACO1–Ni(II) (purple) and PhACO1–Ni(II)–POA (yellow). (b) Pair-distance distribution [*p*(r)] functions showing differences in maximum intraparticle distance (Dmax) between the two states. (c, d) Guinier plots for PhACO1–Ni(II) (purple) and PhACO1–Ni(II)–POA (yellow), confirming structural compaction upon POA binding. (e, f) Ensemble optimization method (EOM) distributions for PhACO1–

Ni(II) (purple) and PhACO1–Ni(II)–POA (yellow), with the pool-predicted distribution shown in black. The shift toward a shorter maximum intraparticle distance (Dmax) upon POA binding indicates a more compact conformational ensemble.

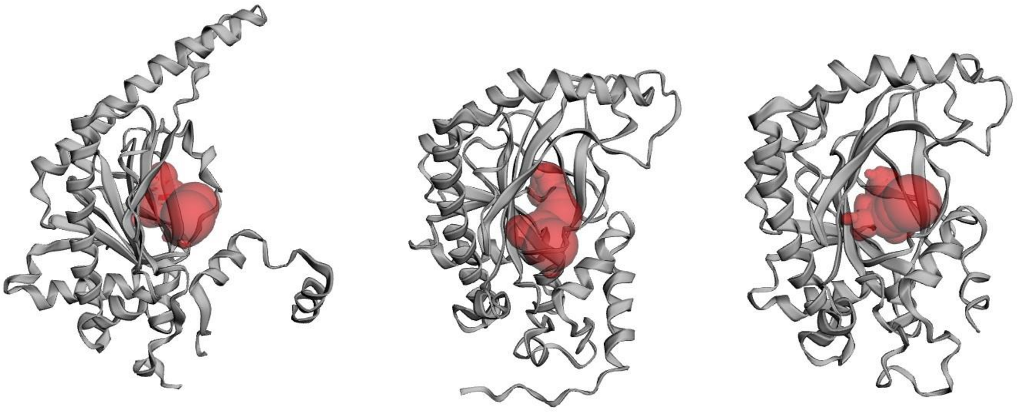

**PhACO1 (PDB: 5TCV)**
**Area (SA) ($Å^2$): 309.382**
**Volume (SA) ($Å^3$): 264.470**

**PhACO1 (AlphaFold)**
**Area (SA) ($Å^2$): 327.367**
**Volume (SA) ($Å^3$): 227.097**

**AtACO2 (PDB:5GJ9)**
**Area (SA) ($Å^2$): 302.189**
**Volume (SA) ($Å^3$): 199.599**

**Supplementary Figure S14 | Active-site pocket area and volume for PhACO1 and AtACO2 in different conformations.** Left: PhACO1–Ni(II) (PDB: 5TCV, open conformation) with pocket area 309.4 $Å^2$ and volume 264.5 $Å^3$. Center: AlphaFold model of PhACO1 (closed conformation) with area 327.4 $Å^2$ and volume 227.1 $Å^3$. Right: AtACO2–Ni(II)–POA (PDB: 5GJ9, closed conformation) with area 302.2 $Å^2$ and volume 199.6 $Å^3$. Analysis used CASTpFold[2].

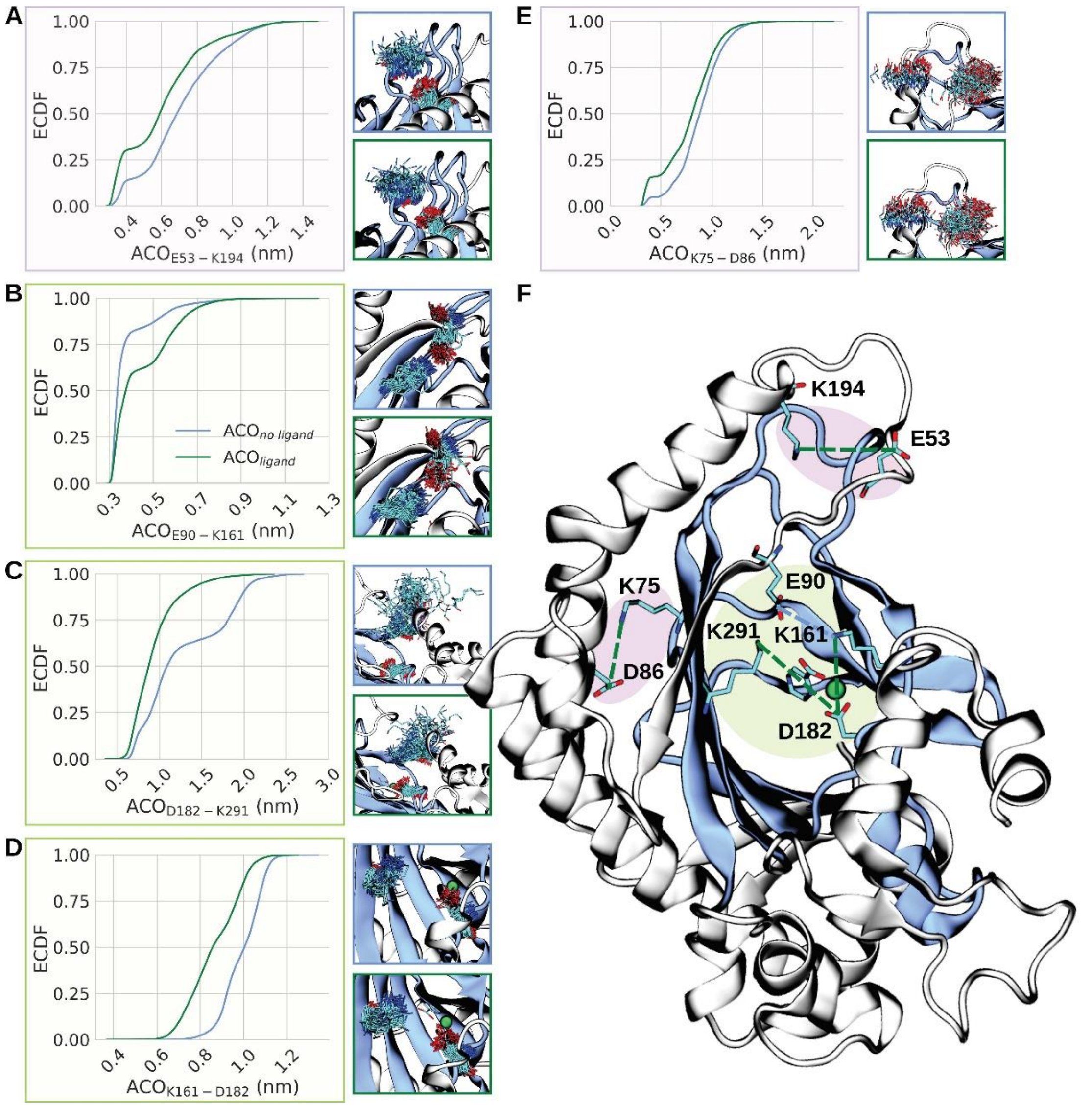

**Supplementary Figure S15 | Effect of POA binding on intramolecular salt-bridge networks in ACO.** Effect of POA binding on intramolecular salt-bridge networks in ACO. (A–E) Empirical cumulative distribution functions (ECDFs) for distances between selected residue pairs: (a) E53–K194, (b) E90–K161, (c) D182–K291, (d) K161–D182, and (e) K75–D86, comparing POA-bound (green) and POA-free (blue) states. Insets show structural ensembles of interacting residues under each condition. (F) Representative ACO monomer conformation highlighting charged residues within or near the Fe–2OG binding domain (light green halo) and the rest of the protein (light purple halo). Secondary structure is shown as cartoons (silver; blue for Fe–2OG domain), with Fe(II) depicted as a green sphere.

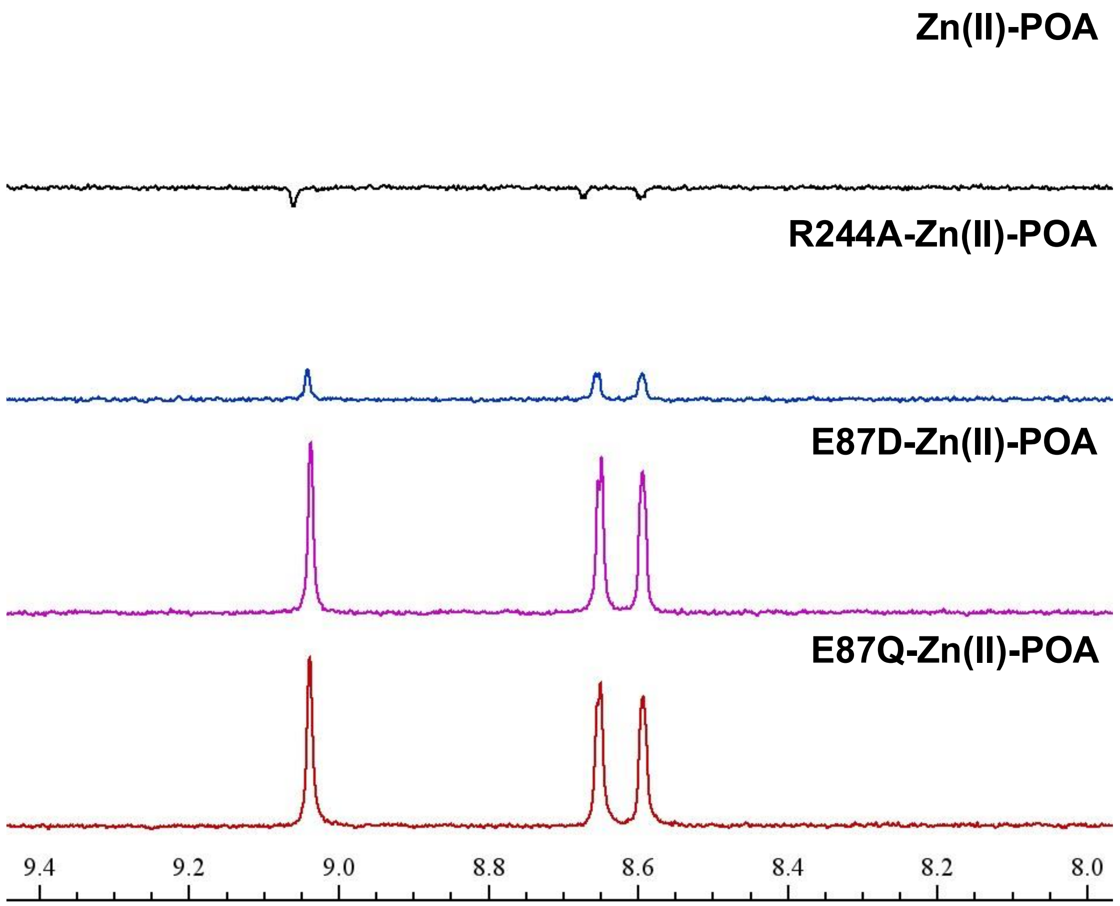

**Supplementary Figure S16 | Proton NMR spectra showing POA interaction with PhACO1 mutants in the presence of $Zn^{2+}$ using the ligand-observed waterLOGSY method.** Spectra are shown for POA with Zn(II) alone (top) and with PhACO1 mutants R244A, E87D, and E87Q (bottom traces). Samples (600 µL) contained 400 µM POA, 10 µM $ZnCl_2$, 50 mM Tris-$d_{11}$ (pH 7.5), and 10% $D_2O$, with or without 10 µM enzyme.

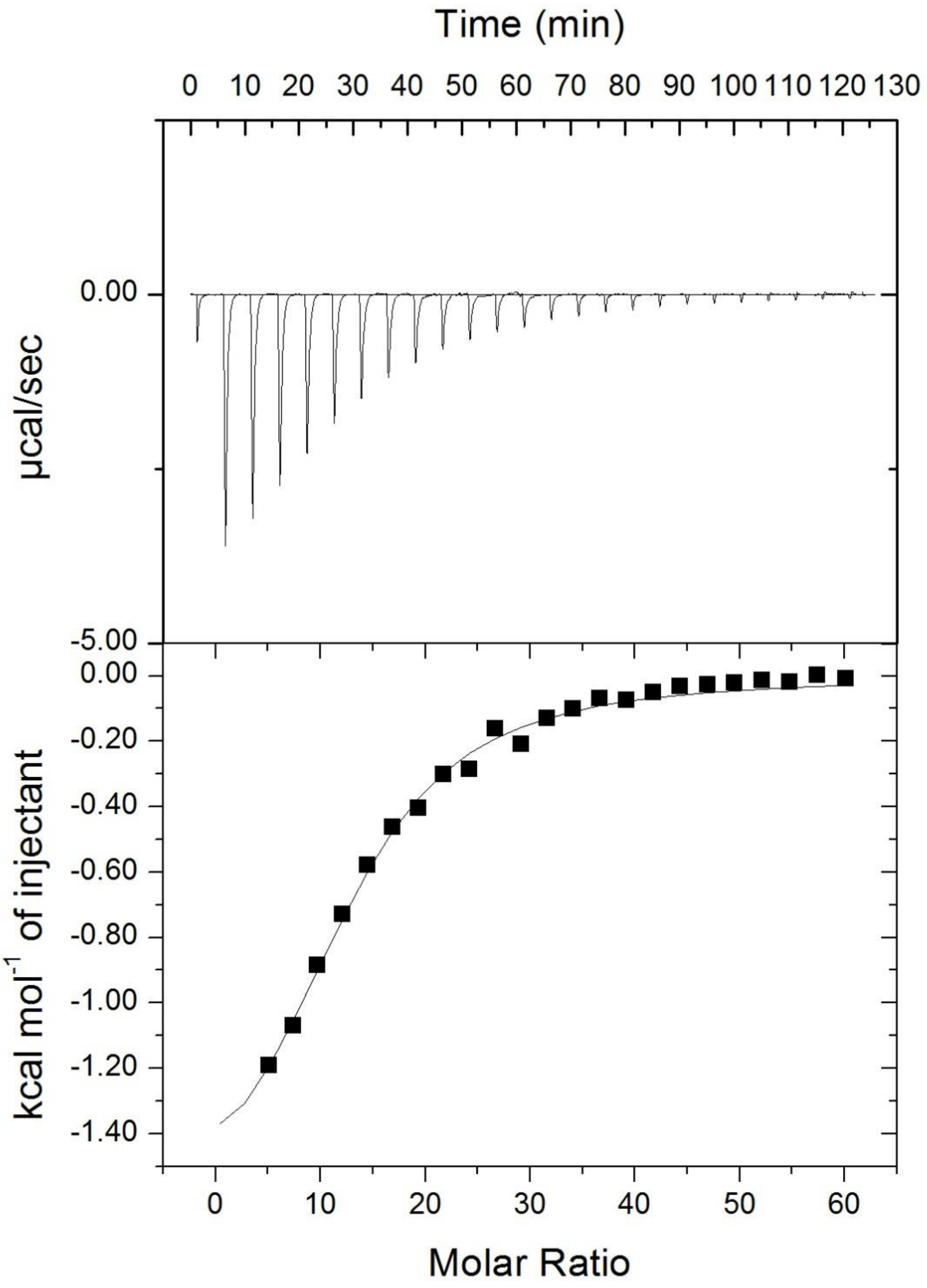

**Supplementary Figure S17 | ITC analysis of POA binding to R244A PhACO1–Ni(II).** Top: Raw ITC data for 25 injections of POA (first 2 µL, then 10 µL; 10 mM POA stock; 300 s interval) into a cell containing R244A PhACO1 (30 µM) with 100 µM Ni(II). Bottom: Baseline-corrected integrated titration curve, yielding a $K_D$ of ~104 µM.

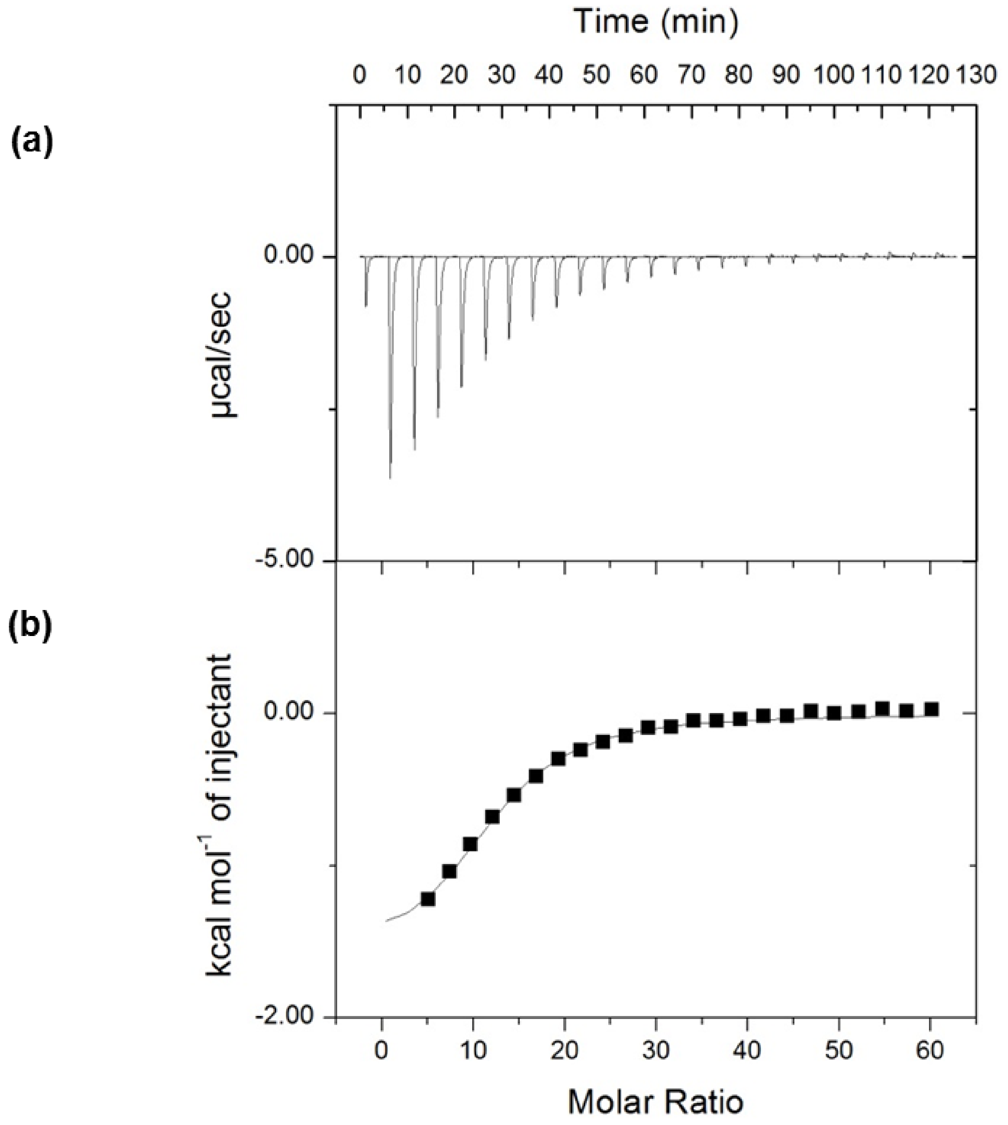

**Supplementary Figure S18 | ITC analysis of POA binding to wild-type PhACO1–Ni(II).** (a) Raw ITC data for 25 injections of POA (first 2 µL, then 10 µL; 10 mM POA stock; 300 s interval) into a cell containing WT PhACO1 (30 µM) with 100 µM Ni(II). (b) Baseline-corrected integrated titration curve, yielding a $K_D$ of ~69.5 µM.

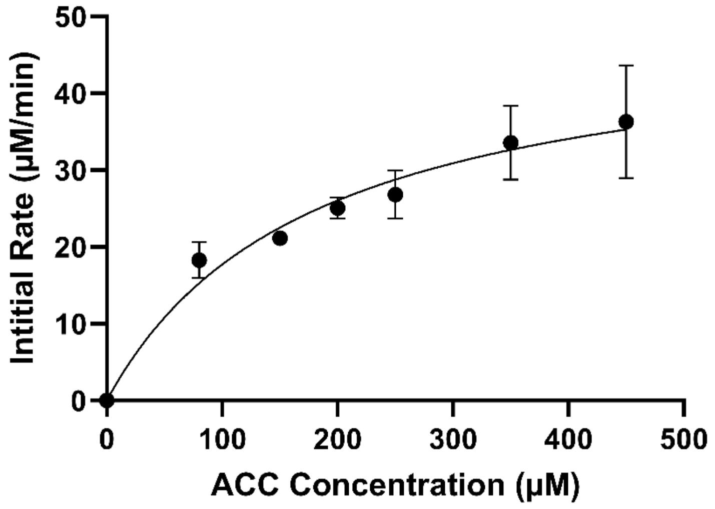

**Supplementary Figure S19 | Michaelis-Menten kinetics analysis of WT PhACO1.** $^{1}H$ NMR assays at ACC concentrations of 80, 150, 200, 250, 350, and 450 µM. The fitted curve, with ($R^2$ = 0.85 ) (n = 3), yielded $V_{max}$ = 49 µM/min and $K_M$ = 178 µM. Reaction mixtures contained 12.5 mM ascorbate, 30 mM $NaHCO_3$, 20 µM Fe(II), 250 µg catalase, and 1 µM PhACO1 in 50 mM Tris-D11 buffer with 10% glycerol-$D_8$, 90% $H_2O$, and 10% $D_2O$. Experiments were conducted at 25 °C.

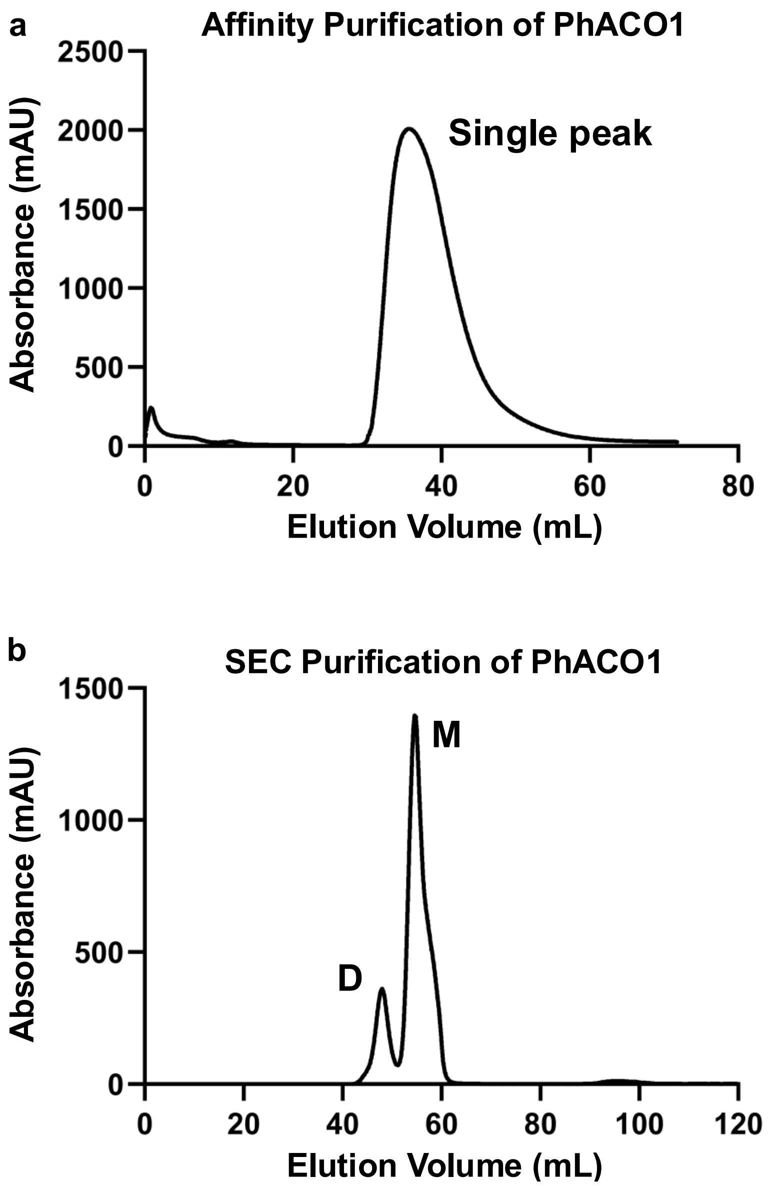

**Supplementary Figure S20 | Purification of PhACO1.** (a) By immobilized metal affinity chromatography (IMAC) and (b) size-exclusion chromatography (SEC). (a) IMAC elution profile showing a single peak after isocratic elution with 500 mM imidazole following a 100 mM imidazole wash. (b) SEC profile of IMAC-purified PhACO1 showing two peaks corresponding to dimer (D) and monomer (M) species. SEC was performed in 50 mM Tris-HCl (pH 7.5) and 150 mM NaCl and used a HiLoad™ Superdex 75 pg.

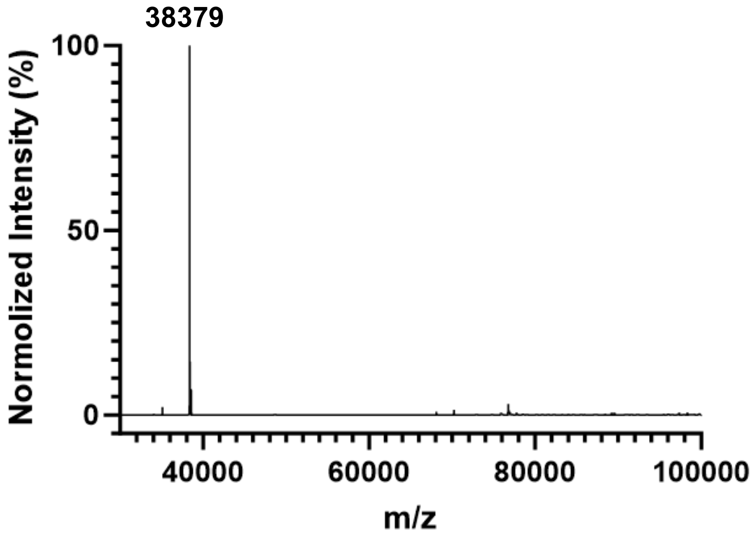

**Supplementary Figure S21 | Intact protein mass spectrum of PhACO1 dimer treated with 2 mM TCEP.** The spectrum shows that the protein was reduced to its monomeric form. Observed molecular weight (MW): 38,379 Da; calculated MW for the monomer after loss of N-terminal methionine: 38,377.03 Da.

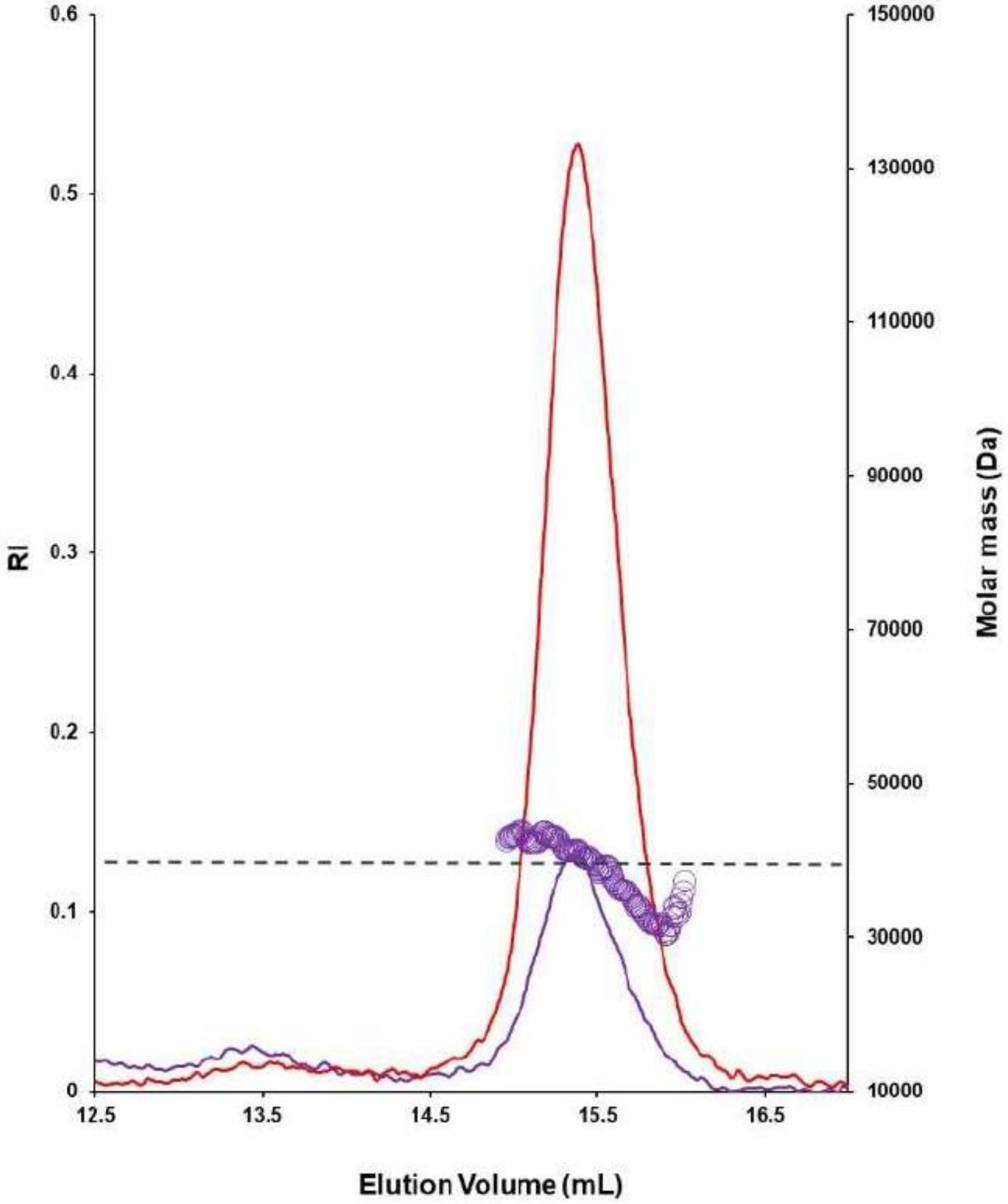

**Supplementary Figure S22 | SEC–MALS chromatogram with refractive index (RI) and molecular mass determination for apo-PhACO1 samples treated with 2 mM TCEP.** The profiles include monomeric apo-PhACO1 (purple) and dimeric apo-PhACO1 (red). Both elution peaks are symmetrical, with an average molar mass of ~40 kDa, indicating that TCEP reduces the dimer to the monomer by breaking a disulfide bond. Differences in peak height reflect varying protein concentrations.

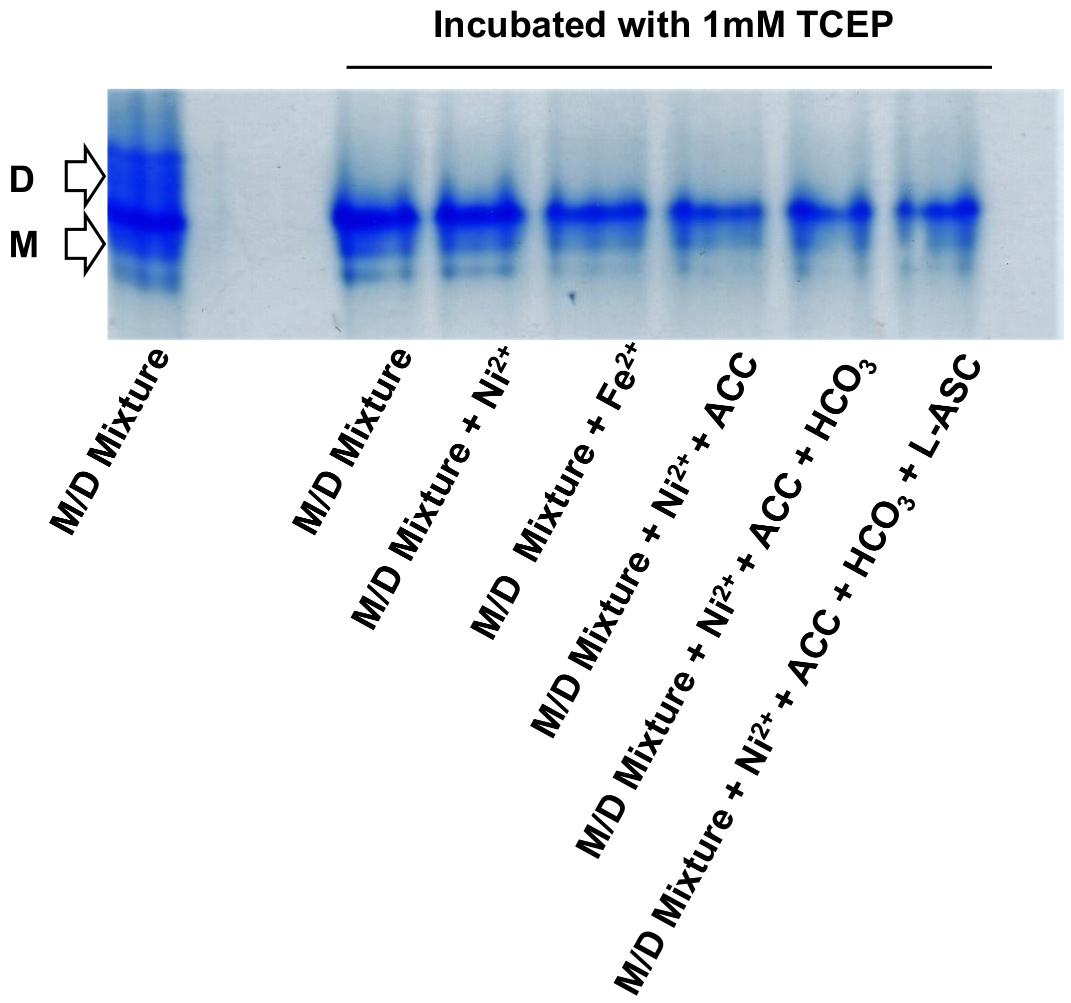

**Supplementary Figure S23 | Native PAGE analysis of PhACO1 showing that the dimer readily reduces to a monomer in the presence of 1 mM TCEP.** Different co-factors ($Ni^{2+}$, $Fe^{2+}$) and co-substrates (ACC, $HCO_3^-$, L-ascorbate) do not affect the reduction. Bands corresponding to dimer (D) and monomer (M) are indicated on the left.

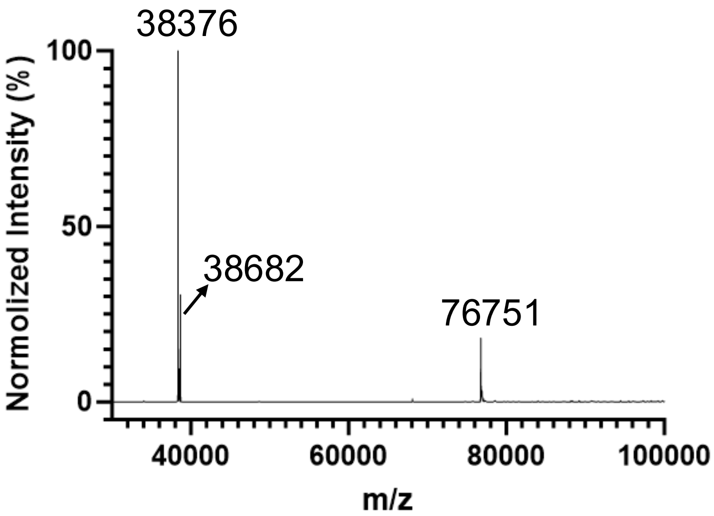

**Supplementary Figure S24 | Intact protein mass spectrum of PhACO1 dimer treated with 2 mM GSH.** The spectrum indicates partial reduction to the monomeric form and glutathionylation of the monomer. Observed monomer MW: 38,376 Da; calculated monomer MW (after loss of N-terminal methionine): 38,377.03 Da; observed glutathionylated monomer MW: 38,682 Da (expected: 38,683 Da); observed dimer MW: 76,751 Da (expected disulfide-linked dimer MW: 76,750 Da).

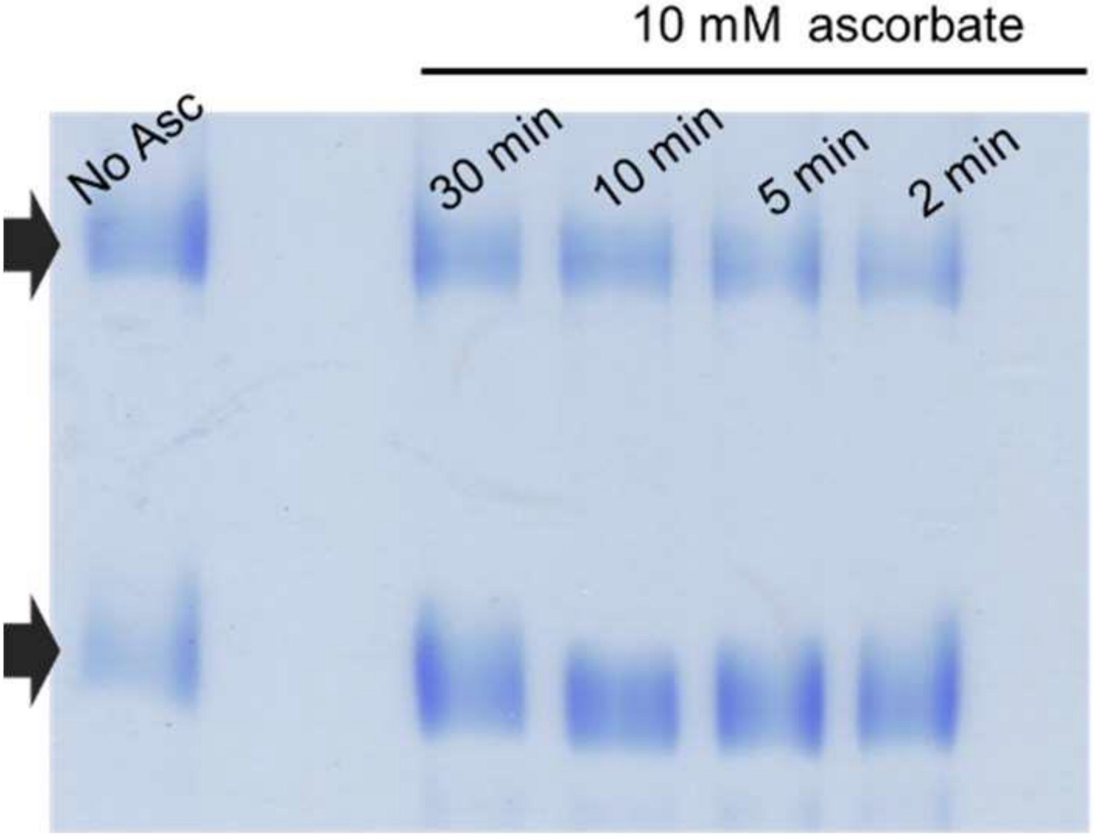

**Supplementary Figure S25 | Native PAGE analysis of PhACO1 with ascorbate (Asc).** A mixture of PhACO1 monomer and dimer (2 mg/mL) with 100 μM Ni(II) was incubated in 10 mM ascorbate for 2, 5, 10, and 30 min. The first lane shows PhACO1 with Ni(II) in the absence of ascorbate as a control.

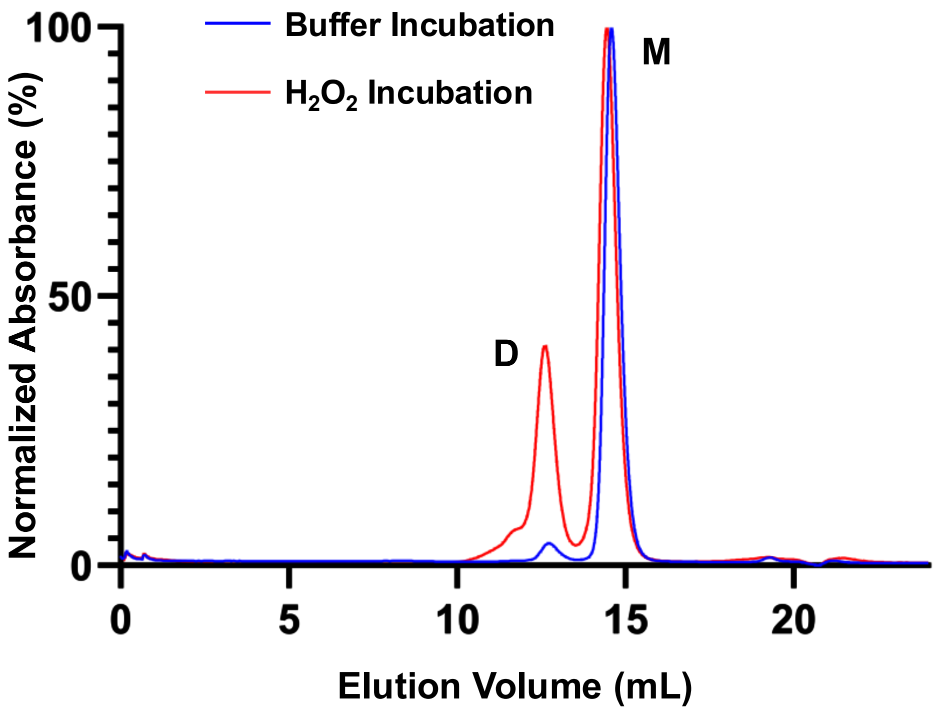

**Supplementary Figure S26 | Size-exclusion chromatography analysis of PhACO1 after incubation with $H_2O_2$.** Samples of 250 µM PhACO1 were incubated with (red) or without (blue) 500 µM $H_2O_2$ in 50 mM Tris-HCl (pH 7.5) and 150 mM NaCl. Chromatography was performed on a Superdex 200 Increase 10/300 GL column, showing elution profiles for dimer (D) and monomer (M) species.

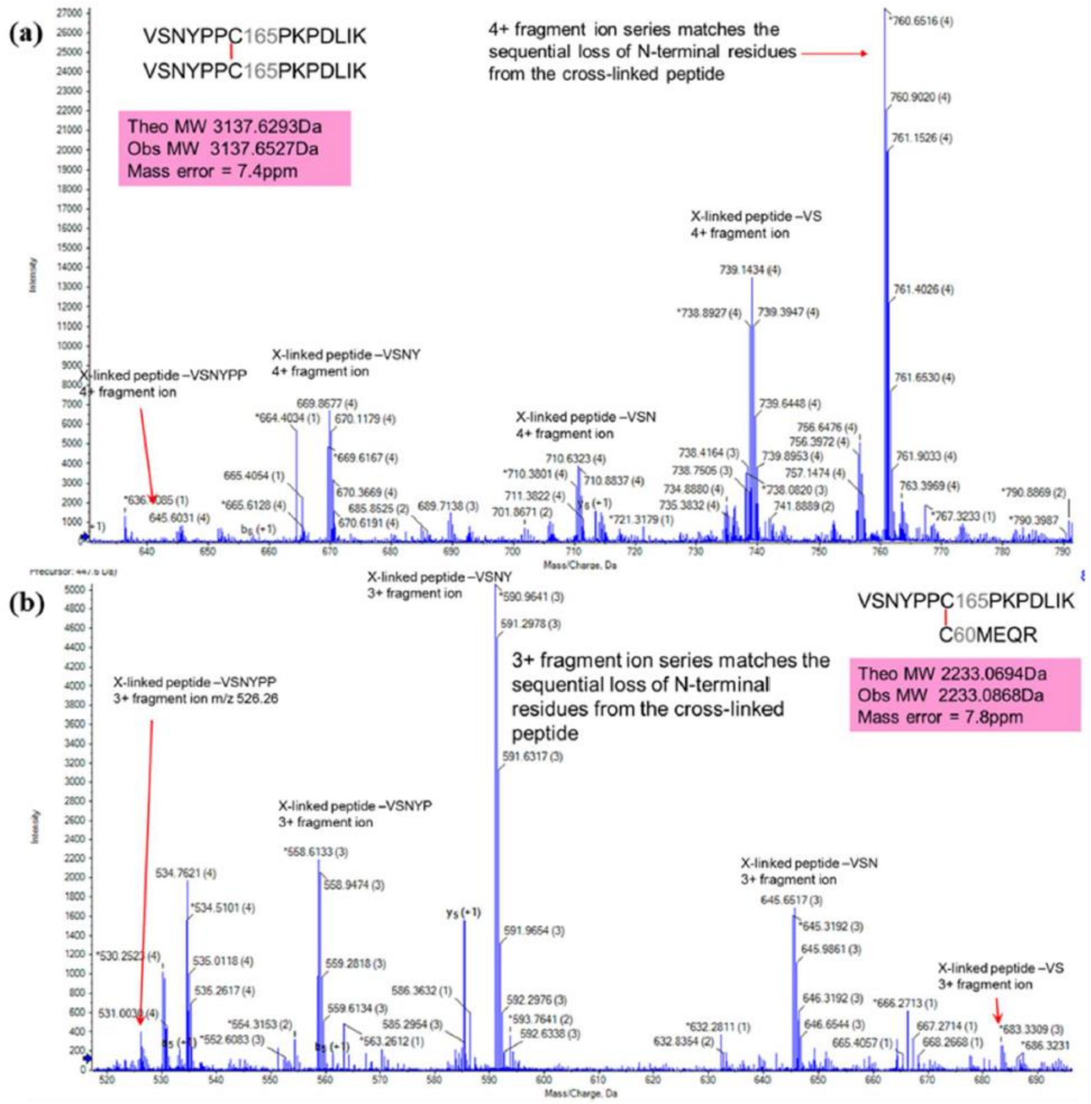

**Supplementary Figure S27 | Tandem MS/MS spectra of disulfide-linked peptides in the PhACO1 dimer.** (a) Major disulfide-linked peptide: homodimer formed via Cys165 residues from each monomer. (b) Minor disulfide-linked peptide: heterodimer formed via Cys60 and Cys165 residues from each monomer.

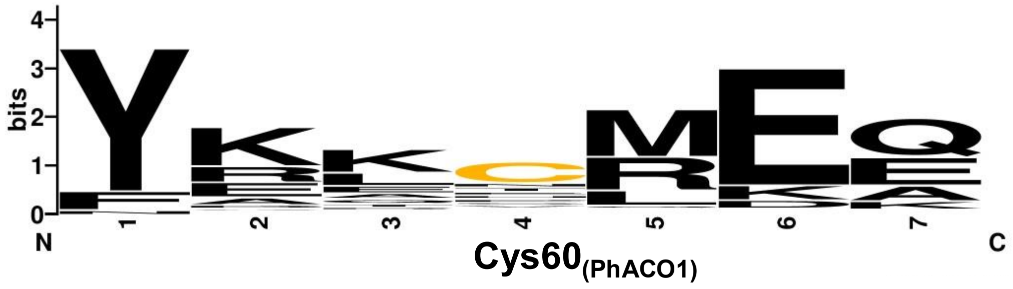

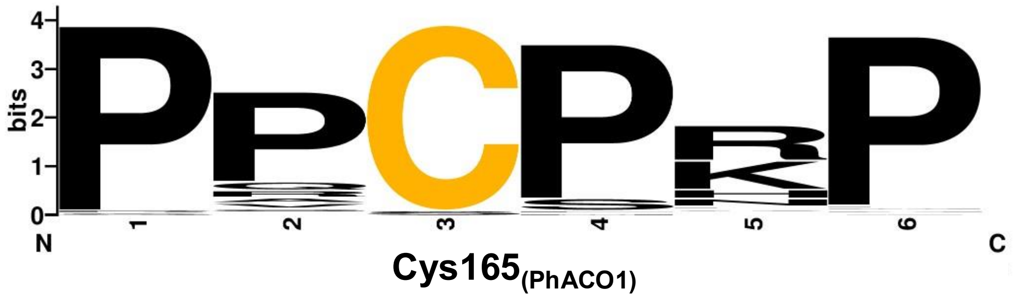

**Supplementary Figure S28 | Protein sequence alignment of 70 ACOs from 12 plant species**. Peptides containing Cys60 and Cys165 were analysed using the WebLogo server. Conservation of each amino acid is represented by letter size, with cysteine residues highlighted in yellow to indicate their significance in the alignment.

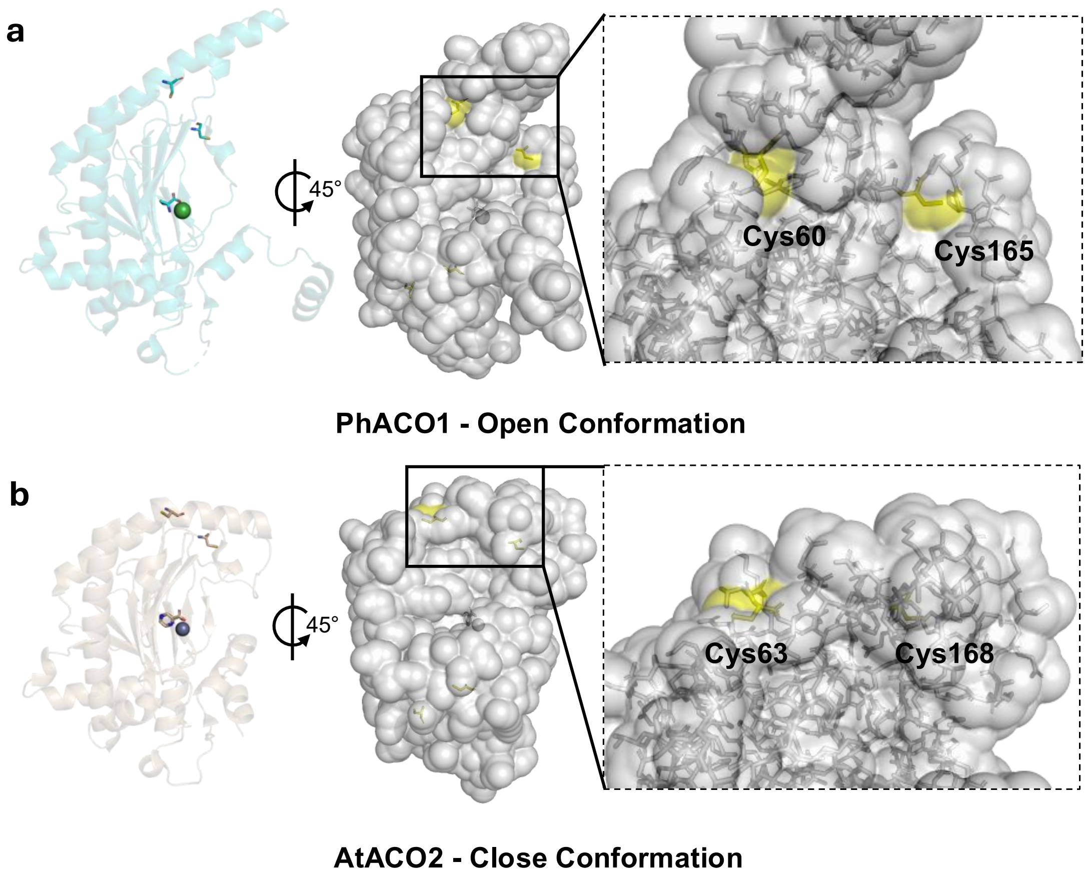

**Supplementary Figure S29 | Surface exposure analysis.** (a) PhACO1 (PDB: 5TCV) and (b) AtACO2 (PDB: 5GJ9) showing that Cys60/Cys63 and Cys165/Cys168 are solvent-accessible in the open conformation but become buried in the closed conformation.

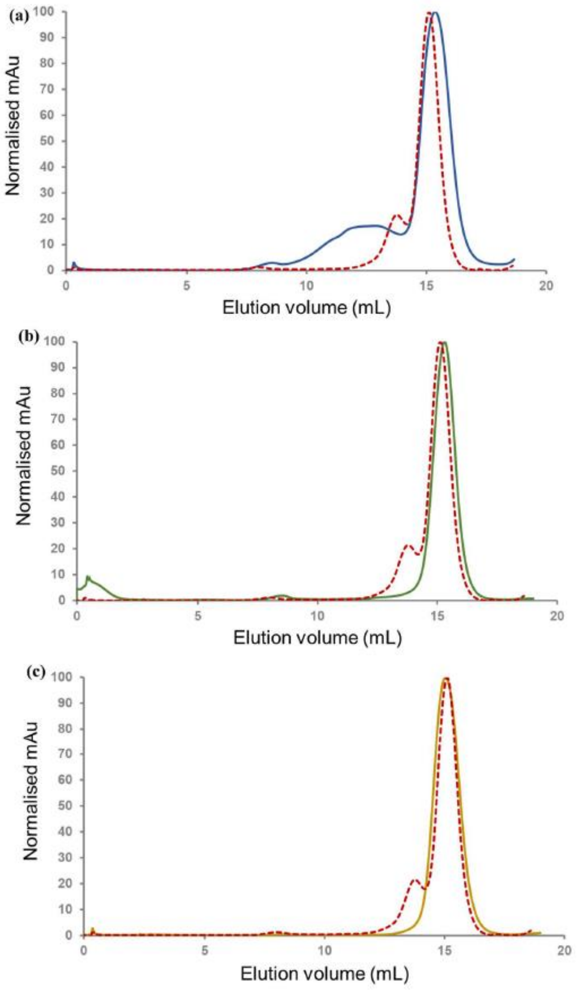

**Supplementary Figure S30 | Analytical size-exclusion chromatograms of PhACO1 mutants compared with wild-type PhACO1 (red dashed lines).** (a) C60S (blue), (b) C165S (green), and (c) C60/165S (yellow). All samples were at 2 mg/mL in 50 mM Tris-HCl (pH 7.5). SEC used a Superdex 200 Increase 10/300 GL column.

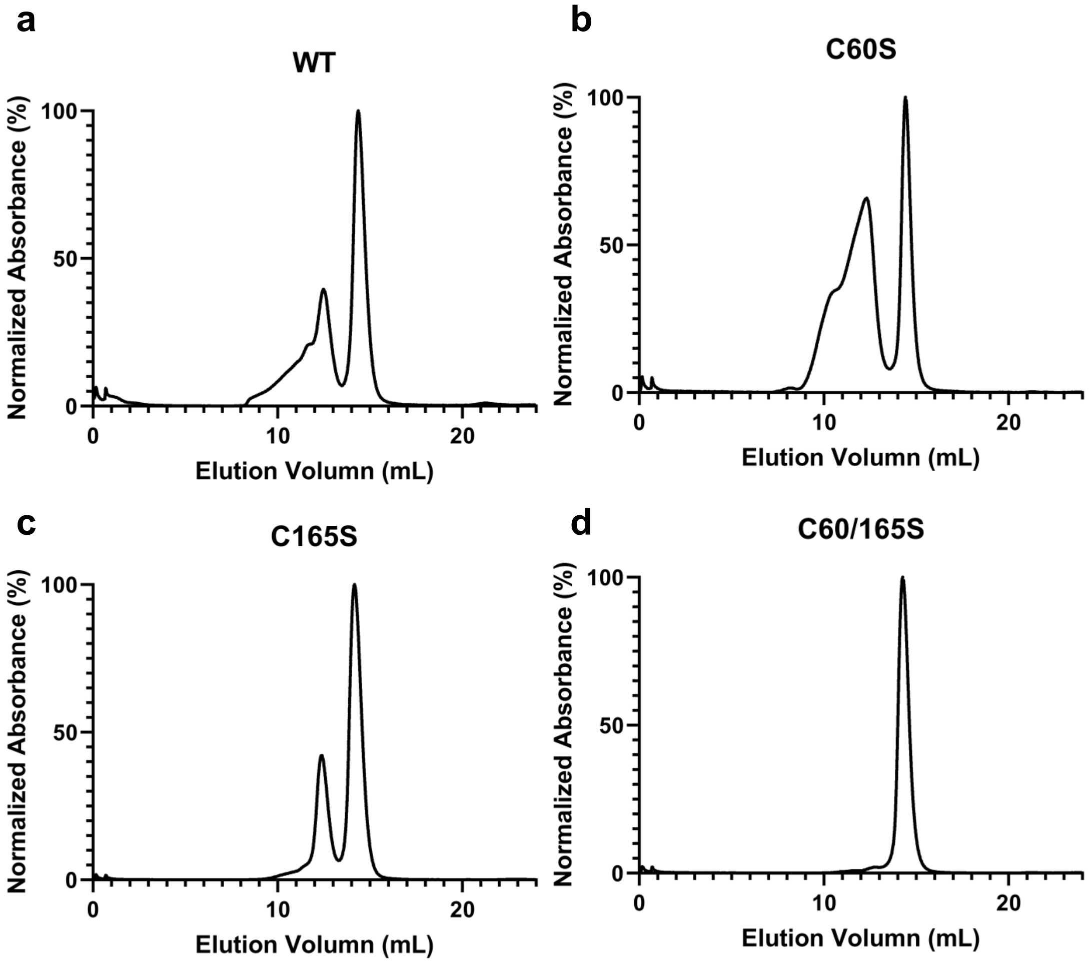

**Supplementary Figure S31 | Size-exclusion chromatography analysis of PhACO1 and its cysteine mutants after incubation with $H_2O_2$.** Samples contained 250 µM protein incubated with 500 µM $H_2O_2$ in 50 mM Tris-HCl (pH 7.5) and 150 mM NaCl. Chromatography used a Superdex 200 Increase 10/300 GL column.

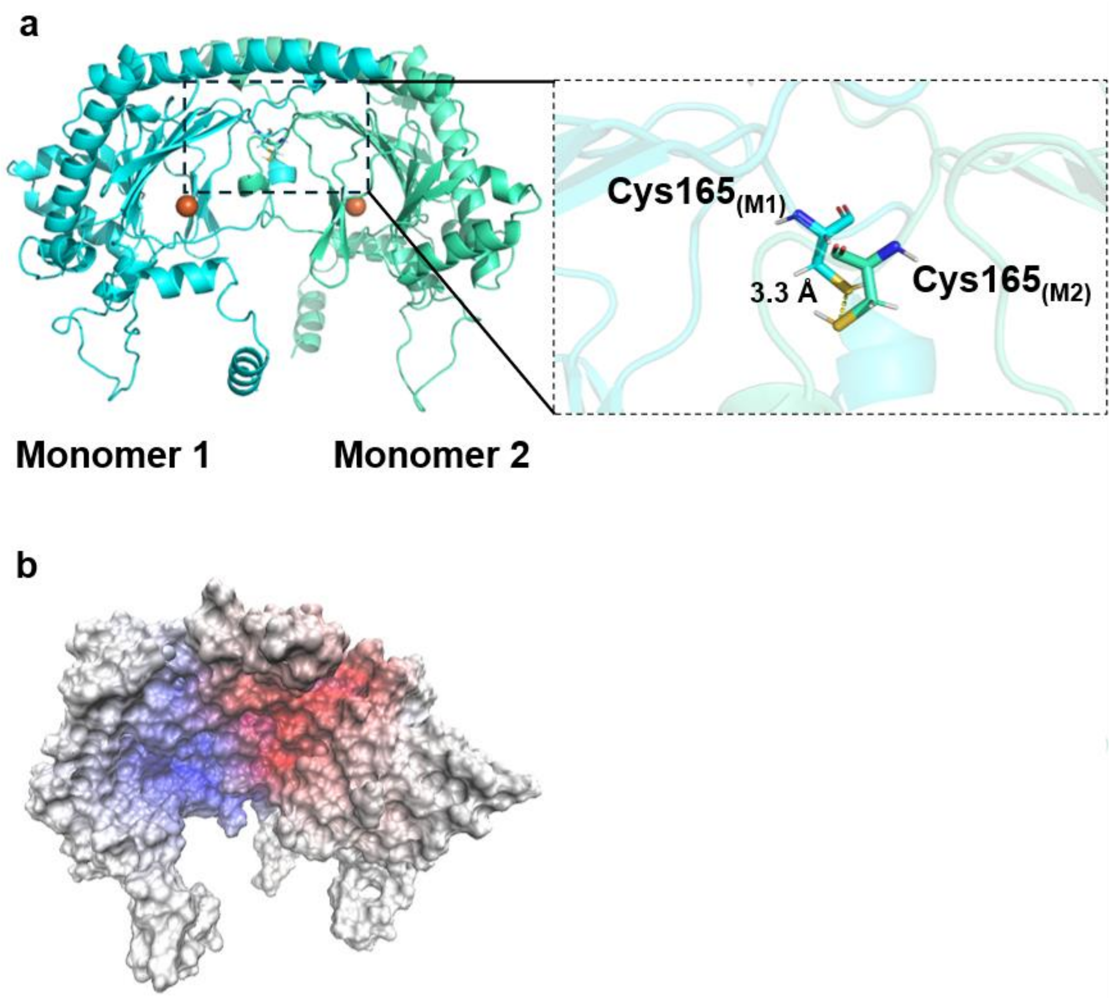

**Supplementary Fig. S32 | Predicted PhACO1 dimer from shape-complemented docking.** Docking was performed with PatchDock using the crystal structure of PhACO1–Ni(II)–ACC (PDB 5TCV) and a distance constraint to position Cys165 residues within disulfide bond range (4 Å). (a) Predicted dimer structure with close-up showing Cys165 residues from each monomer (3.3 Å apart). (b) Electrostatic potential map of the predicted dimer; red indicates negative potential; blue indicates positive potential.

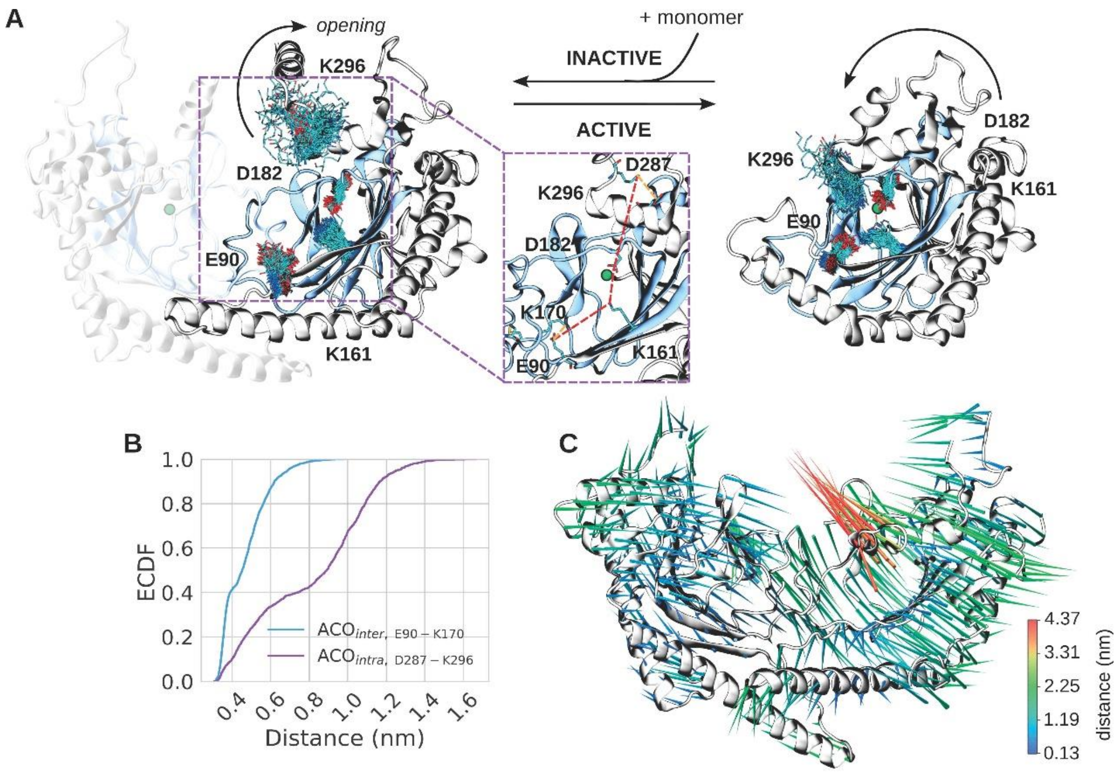

**Supplementary Fig. S33 | MD simulations reveal salt-bridge network formation upon PhACO1 dimerisation.** (a) Representative conformations of charged residues near the Fe–2OG binding domain (blue) for the dimer (left) and monomer (right). Arrows indicate activation state. Protein shown as cartoon (silver), Fe–2OG domain in blue, Fe(II) as green sphere. Inset: salt-bridge network in the dimer; red dashed lines indicate broken bridges, orange indicate newly formed bridges. (b) Empirical cumulative distribution functions for E90–K170 (intermolecular, blue) and D287–K296 (intramolecular, purple) salt-bridge distances. (c) Porcupine plot of ACO dynamics from principal component analysis. Cones represent Cα movements; length and colour scale with magnitude of conformational change.

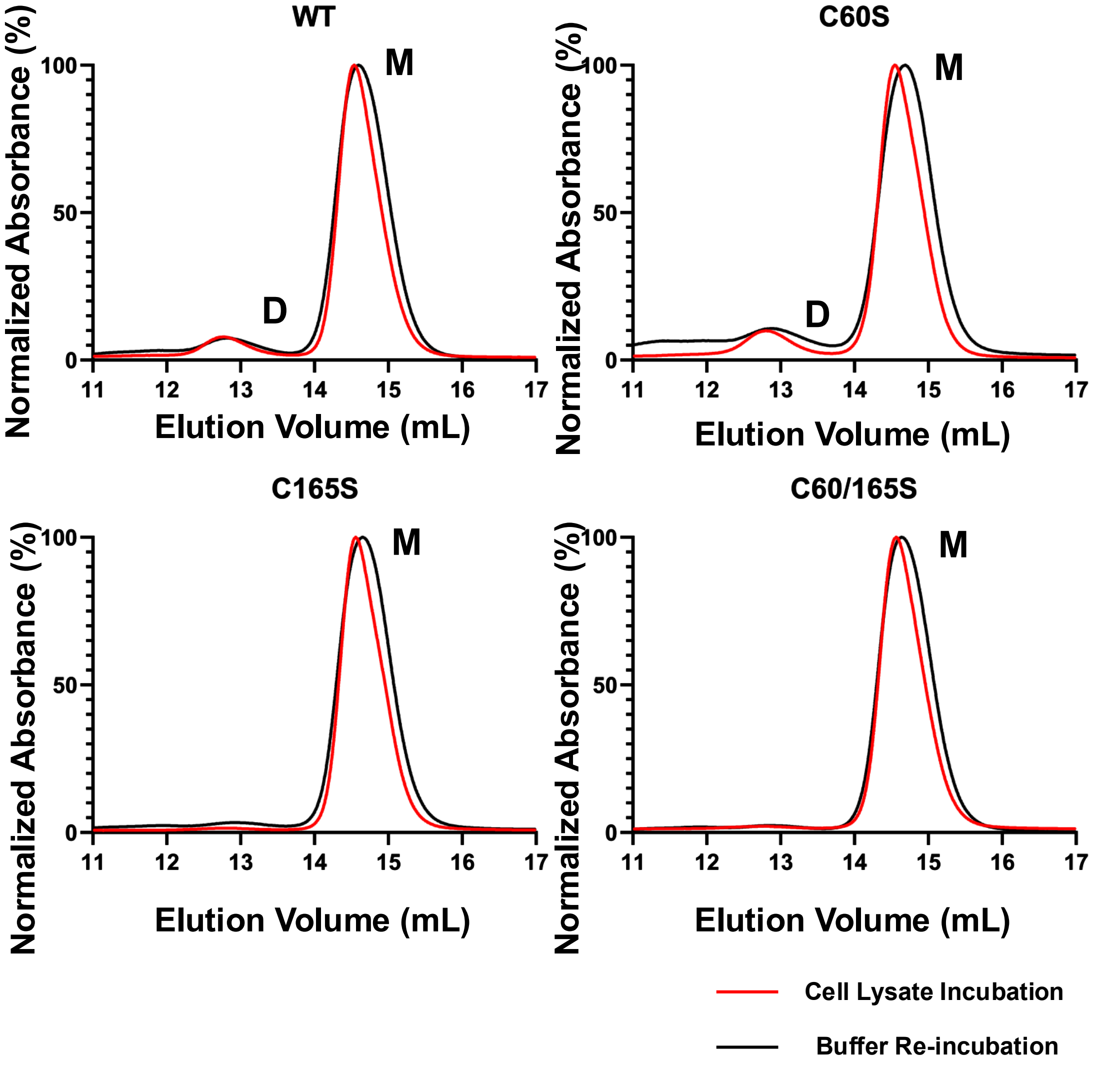

**Supplementary Fig. S34 | Size-exclusion chromatography indicates PhACO1 remains monomeric after plant lysate incubation and $H_2O_2$ treatment.** Monomeric PhACO1 (250 μM) was incubated with 2.5 mM Ni(II) in *A. thaliana* ACO-free plant lysate (red), purified, and re-incubated in 50 mM Tris-HCl (pH 7.5) containing 150 mM NaCl and 500 μM $H_2O_2$ (black). Elution profiles were obtained using a Superdex 200 Increase 10/300 GL column. WT and cysteine mutants (C60S, C165S, C60/165S) show monomer peaks (M) with no evidence of dimer formation (D).

**a** **PhACO1 in Un-treated Plant Lysate**

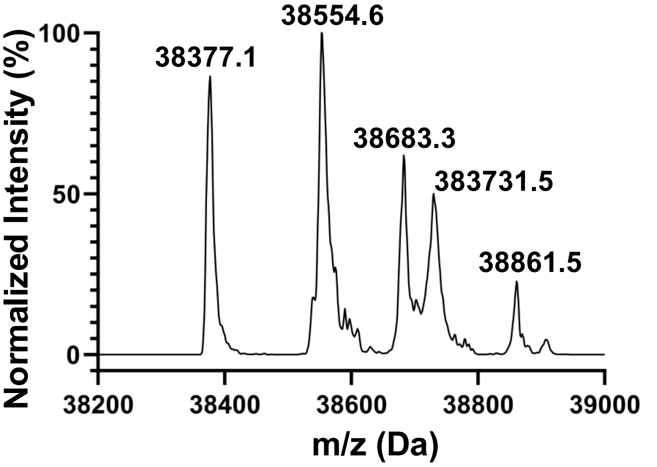

| Observed (Da) | Differences (Da) | Modifications | Expected (Da) |
|---|---|---|---|
| 38377.1 | 0 | Monomer | 38377.0 |
| 38554.6 | 177.5 | 1*Cys-Gly | 38553.2 |
| 38683.3 | 306.2 | 1*GSH | 38682.4 |
| 38731.5 | 354.4 | 2*Cys-Gly | 38729.4 |
| 38861.5 | 484.4 | 1*Cys-Gly + 1*GSH | 38858.6 |

**b** **PhACO1 in Plant Lysate Contained 5mM TCEP**

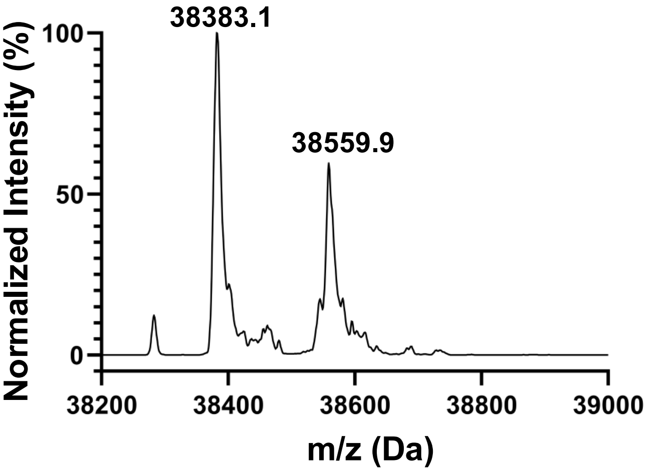

| Observed (Da) | Differences (Da) | Modifications | Expected (Da) |
|---|---|---|---|
| 38383.1 | 0 | Monomer | 38377.0 |
| 38559.9 | 176.8 | 1*Cys-Gly | 38553.2 |

**Supplementary Fig. S35 | Intact protein mass spectra of PhACO1 after plant lysate incubation with or without TCEP.** PhACO1 (500 µg) was incubated in *A. thaliana* ACO-free plant lysate in 50 mM potassium phosphate buffer (pH 7.5) (a) without and (b) with 5 mM TCEP. Addition of TCEP reduced modified species, indicating that glutathione and cysteinyl-glycine adducts were linked via disulfide bonds.

**a**

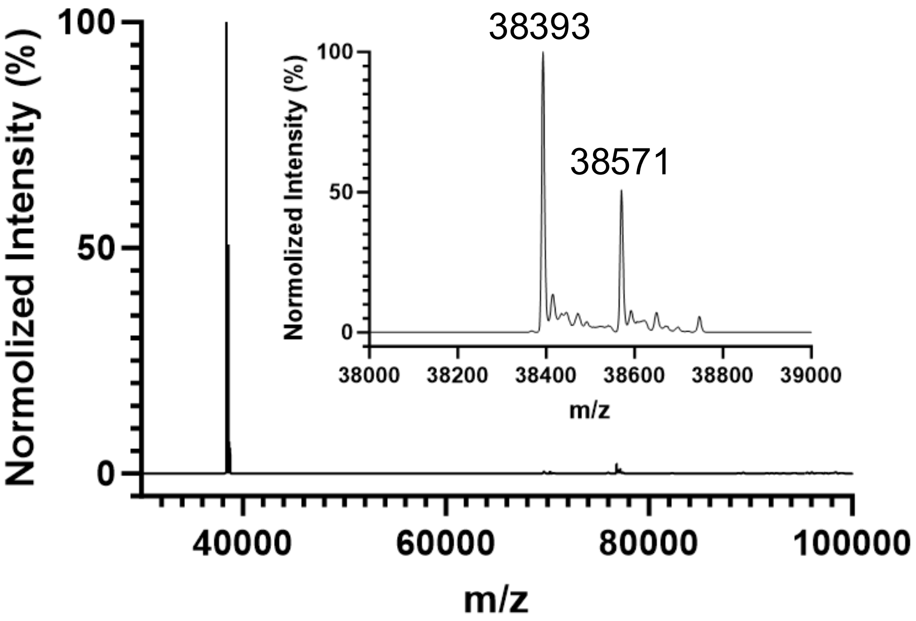

**b**

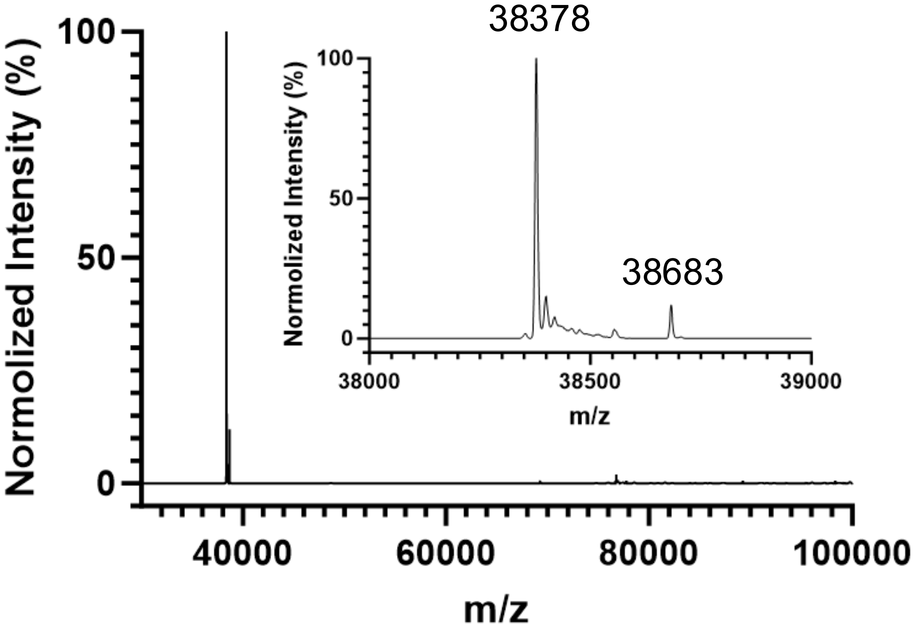

**Supplementary Fig. S36 | Intact protein mass spectra of PhACO1 after incubation with cysteinyl-glycine or oxidised glutathione.** PhACO1 (1 mg/mL) was incubated for 30 min with (a) 2 mM cysteinyl-glycine or (b) 2 mM oxidised glutathione. Insets show zoomed spectra highlighting modified species.

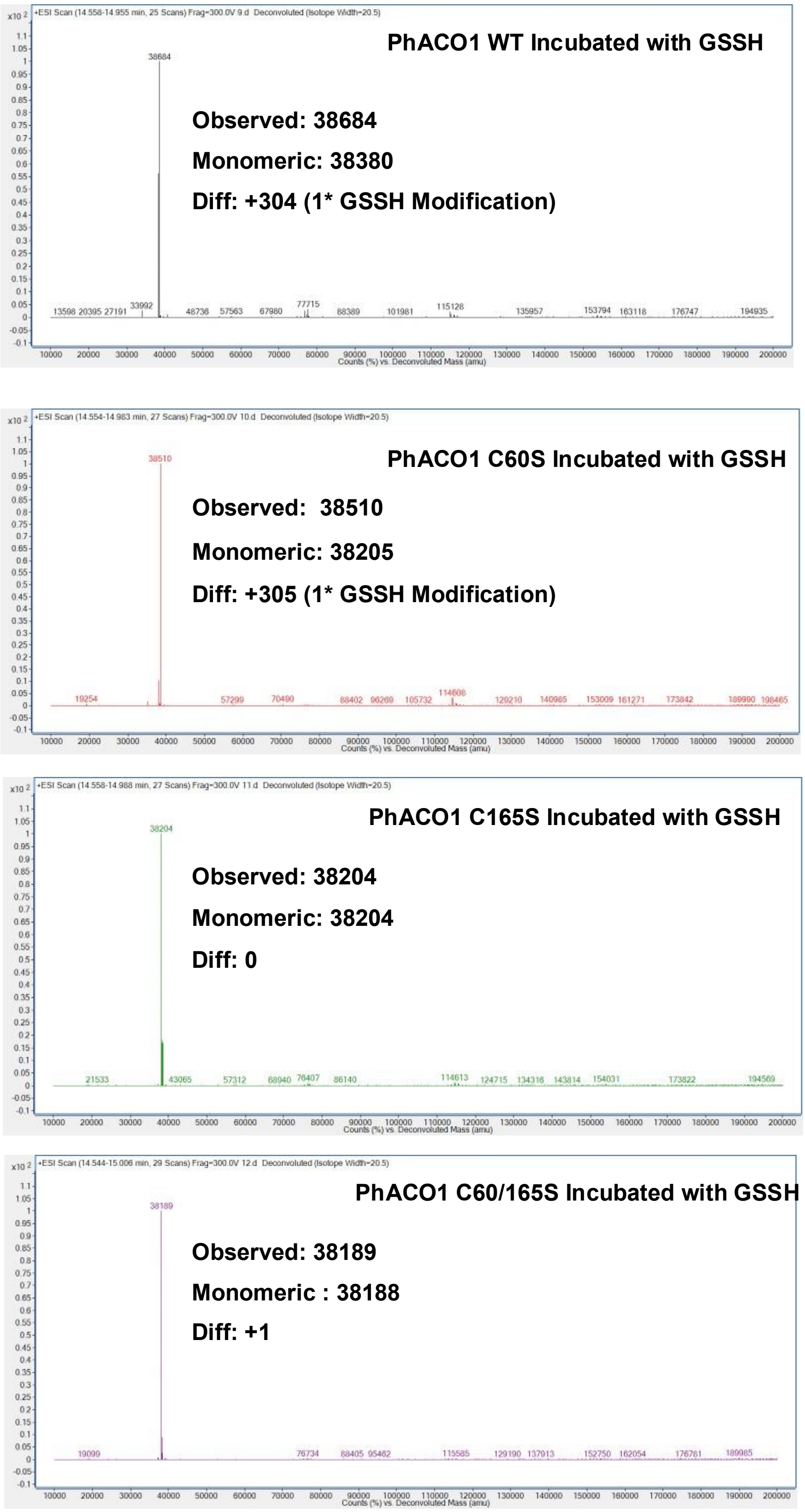

**Supplementary Fig. S37 | Intact protein mass spectrometry of PhACO1 wild type and cysteine mutants after GSSH incubation.** Wild type and mutants (C60S, C165S, C60S/C165S) at 1 mg/mL were incubated with 2 mM oxidised glutathione (GSSH) in Tris-HCl buffer (pH 7.5)

for 30 min. Spectra show mass shifts corresponding to GSSH modification in WT and C60S, while C165S and C60S/C165S remain unmodified.

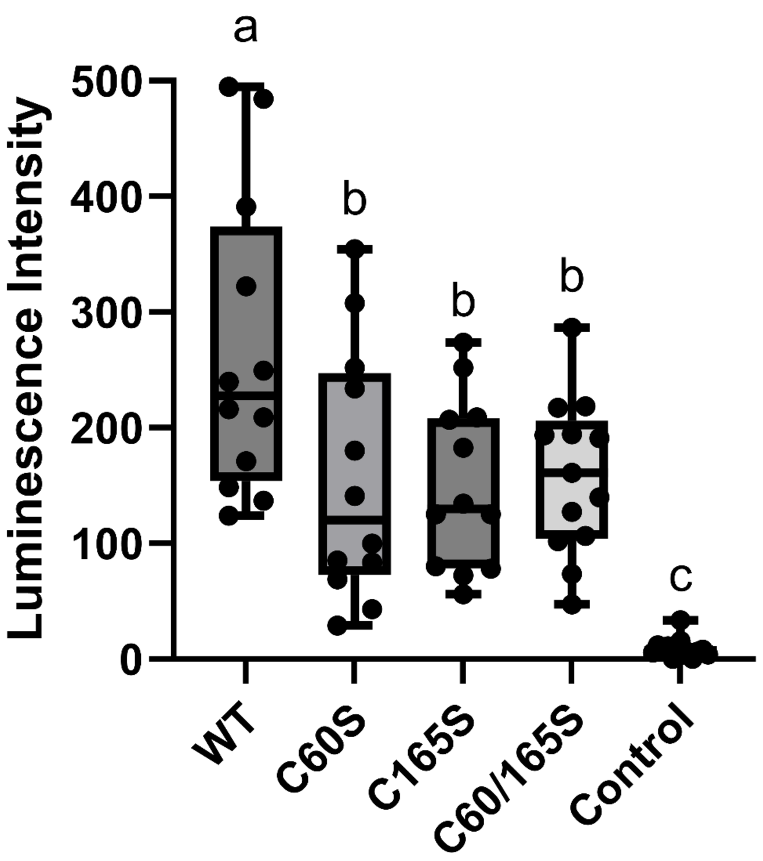

**Supplementary Fig. S38 | Luminescence intensity of PhACO1 wild type and cysteine mutant combinations in split luciferase assay.** Box plots show luminescence intensity for WT and mutants (C60S, C165S, C60S/C165S) compared to control. Each point represents an individual biological replicate ($n \geq 12$). Statistical significance was assessed using Welch's t-test; different letters indicate groups with significant differences.

| | PhACO1-Ni(II)-acetate<br>PDB: 5TCW | PhACO1-Ni(II)-acetate<br>PDB: 5TCW |
|---|---|---|
| X-ray source,<br>wavelength (Å) | Australian Synchrotron<br>MX-1, 0.9537 | Australian Synchrotron<br>MX-1, 0.9537 |
| Detector | ADSC QUANTUM 201r | ADSC QUANTUM 201r |
| Space Group | *I* 2 2 2 | *I* 2 2 2 |
| Unit cell parameters<br>(Å) | a = 70.94<br>b = 108.14<br>c = 108.32 | a = 71.36<br>b = 107.14<br>c = 108.02 |
| Resolution range (Å) | 19.8 – 2.70 (2.83 – 2.70) | 40.0 – 2.60 (2.72 – 2.60) |
| Number of Unique<br>Reflections | 12,054 (1,497) | 13,115 (1,650) |
| Multiplicity | 14.5 (13.6) | 43.9 (44.5) |
| Completeness (%) | 99.5 (98.5) | 100 (100) |
| $\langle I/\sigma(I) \rangle$ | 18.1 (1.0) | 30.0 (1.7) |
| $CC_{1/2}$ | 0.999 (0.454) | 1.000 (0.718) |
| Wilson *B*-factor (Å$^2$) | 70.1 | 66.7 |

* Statistics relating to the high-resolution shell are shown in parentheses.

**Supplementary Table S1 | Crystallography data collection statistics for PhACO1-Ni(II)-acetate (PDB: 5TCW) and PhACO1-Ni(II)-ACC (PDB: 5TCV).**

| | PhACO1-Ni(II)-acetate PDB: 5TCW | **PhACO1-Ni(II)-ACC PDB: 5TCV** |
|---|---|---|
| **Resolution range (Å)*** | **19.8 – 2.70 (2.77 – 2.70)** | **40.0 – 2.60 (2.67 – 2.60)** |
| **$R_{work}$, $R_{free}$*** | **0.225, 0.269 (0.365, 0.380)** | **0.232, 0.291 (0.360, 0.387)** |
| **Average *B* factor: protein (Å$^2$)** | **75.3 (2,335 atoms)** | **76.4 (2359 atoms)** |
| **Average *B* factor: ligands, water (Å$^2$)** | **69.9 (1 $Ni^{2+}$)<br>75.6 (1 acetate)** | **59.2 (1 $Ni^{2+}$)<br>70.7 (7 ACC atoms)<br>59.5 (11 HOH molecules)** |
| **RMSD bond lengths (Å)** | **0.006** | **0.006** |
| **RMSD bond angles (°)** | **1.03** | **1.03** |
| **Ramachandran favoured (%)** | **96.3** | **97.3** |
| **MolProbity Score** | **100$^{th}$ Percentile** | **100$^{th}$ Percentile** |

* Statistics relating to the high-resolution shell are shown in parentheses.

**Supplementary Table S2 | Crystallography refinement statistics for PhACO1-Ni(II)-acetate (PDB: 5TCW) and PhACO1-Ni(II)-ACC (PDB: 5TCV).**

**PhACO1 PCR amplification**

| | |
|---|---|
| PhACO1_KpnI_F | AAGGTACCATGGAGAACTTCCCAATTATCAG |
| PhACO1+_PstI_R | AACTGCAGTTAGACAGTGGCAATTGGATC |
| PhACO1-_SalI_R | AAGTCGACGACAGTGGCAATTGGATCC |
| PhACO1_F | ATGGAGAACTTCCCAATTATCAG |
| PhACO1-_R | GACAGTGGCAATTGGATCC |
| PhACO1+_R | TTAGACAGTGGCAATTGGATC |

**pCAMbia1300 vector sequencing**

| | |
|---|---|
| LUC_1371-F | TTGCTCCAACACCCCAACAT |
| M13(-21)_F | TGTAAAACGACGGCCAGT |
| M13_R | CAGGAAACAGCTATGAC |
| 35Sprom_-71F | ACGCACAATCCCACTATCCT |
| LUC+_68_R | CCAGCGGTTCCATCTTCCAG |

**Supplementary Table S3 | List of primers used in cloning and sequencing.**

| ***SAXS data collection parameters:*** | ***PhACO1-Ni(II)*** | ***PhACO1-Ni(II)-POA*** |
|---|---|---|
| *Instrument/source* | Australian Synchrotron SAXS/WAXS beamline equipped with Pilatus 1M detector and sheath-flow cell for SEC-SAXS. | |
| *Wavelength (Å)* | 1.0332 | |
| *Beam energy (kEv)* | 12 | |
| *Beam size (μm)* | 250 × 130 | |
| *Sample-to-detector distance (mm)* | 3256 | |
| *$q$ (Å$^{-1}$)* | 0.005 – 0.334 | |
| *Absolute scaling method* | Comparison with scattering from 1 mm pure water | |
| *Normalization* | To transmitted intensity from beamstop counter | |
| *Exposure time* | 1 s measurements from SEC-SAXS elution | |
| *Sample temperature (K)* | 295 | |
| ***SEC-SAXS parameters:*** | | |
| *Column* | Superdex S200 5×150 | |
| *Flow rate (mL/min)* | 0.4 | |
| *Concentration (mg/mL)* | 5 | |
| *Injection volume (μL)* | 50 | |
| *Average conc. (mg/mL)* | 3.02 | |
| *Solvent* | 50 mM Tris-HCl, pH 7.5, 150 mM NaCl | |
| ***Software employed:*** | | |
| *SAXS data reduction* | *I(q)* vs *q* using Scatterbrain 2.8.2, SEC-SAXS solvent subtraction using *CHROMIXS* from *ATSAS 3.2.1* | |
| *Basic analysis (Guinier, P(r), molecular mass)* | PRIMUSqt from ATSAS 3.2.1 | |

| | | |
|---|---|---|
| *Ab initio modelling* | *DAMMIN* from *ATSAS 3.2.1* and DENSS using DENSSWeb | |
| *Calculation of theoretical intensities* | CRYSOL from ATSAS 3.2.1 | |
| *Atomic structure (hybrid) modelling* | CORAL from ATSAS 3.2.1 | |
| ***Structural parameters:*** | | |
| *Mass from $V_c$ (kDa) (expected mass, ratio to expected in brackets)* | 45.94 (38.51, 1.2) | 36.56 (38.63, 0.9) |
| ***Guinier analysis:*** | | |
| *$R_g$ (Å)* | 29.37 | 27.55 |
| *I(0) ($cm^{-1}$)* | 0.035±0.0001 | 0.043±0.0001 |
| *$qR_g$ min,max* | 0.24, 1.29 | 0.23, 1.26 |
| ***P(r) analysis:*** | | |
| *$R_g$ (Å)* | 29.81 | 27.59 |
| *I(0) ($cm^{-1}$)* | 0.03±0.0001 | 0.04±0.0001 |
| *$D_{max}$ (Å)* | 116.61 | 107.87 |
| *Porod volume (Å$^3$)* | 88215 | 83456 |
| ***Atomic modelling (DAMMIN):*** | | |
| *Processing mode* | Slow | Slow |
| *Symmetry assumptions* | P1 | P1 |
| *Number of iterations* | 100,000 | 100,000 |
| *Number of annealing steps* | 500 | 500 |
| *$\chi^2$ range* | 1.05 | 1.09 |

| ***Ensemble Optimisation Methodology*** | | |
|---|---|---|
| *Starting Structure* | 5TCV (name) | |
| *Symmetry assumptions* | P1 | |
| *No. of models in the pool* | *RANCH* was used to generate a pool of 10000 models | |
| *Theoretical Intensity* | *Crysol* was used to calculate the theoretical intensity along with $R_g$ and $D_{max}$ | |
| *Genetic Algorithm* | Yes, using GAJOE | |
| *$\chi^2$ range* | 1.5 | 1.2 |
| ***Deposition codes*** | | |
| *SASB Code* | SASDXS9 | SASDXT9 |

**Supplementary Table S4 | SAXS results for PhACO1-Ni(II) and PhACO1-Ni(II)-POA.**

>XP_006837262.2 1-aminocyclopropane-1-carboxylate oxidase [Amborella trichopoda]
MGFSFPVVDLQELEGGERKSAMELINDACENWGFFEVVNHGLSQEFMDQVESLTKEHYRKY
MEKRFKDEVAERVLKKEEEVKDLDWESTFYLRHLPSSNISEIPDLDHEYRRVMKEFAGVIE
KLAEKLLDVLCENLGLEKGYLKKAFQGKNGYPTFGTKVSSYPPCPRPELVKGLRAHTDAGG
LVLLFQDPQVSGLQLLKDGEWVDVPPLRHSIVINIGDQLEVITNGRYKSVMHRVVAQTNGN
RMSIASFYNPGSDAVIFPAPTLLKKETAEYPKFVFEDYMKLYVGQKFQAKEPRFETMKAME
TVSLGPIATA

>XP_006844270.2 1-aminocyclopropane-1-carboxylate oxidase 1 [Amborella trichopoda]
MGVPVIDVMKLRGENREHQMALLGDACENWGFFQIVNHGVEIKVMDRLKGLMNDHYEEYMK
PQFQFHDSHVQNELERSPSYVRSFGDWECSFFVKHMPTSNLTLLPQISDDLKQAIHEYVEE
VKKVAEMMLDLFCENLGLERGYLKKVFLGSDGLPMLGTKLAKYPVCPRPDLVRGLRAHTDA
GGIIFLLQDDQVPGLQFVKDGKWVDVPPLANSLFINIGDQLEVITNGRYKSMLHRVLARED
GSRLSIVTFYNPADDAEIGPAPELTYPSRFRFGEYMNIYASTKFSDKQPRFEAMKKHAVDT
PLLV
>XP_020524572.1 1-aminocyclopropane-1-carboxylate oxidase [Amborella trichopoda]
MTIPVIDMSKLDGEERAKTMAEIANGCEEWGFFQLINHGVPVELLERLKKGCSECYKLERE
EGFNKSVPVKMLDEILVAEKEKSEKKVEEEVKKRVEGVDWEDVFVLQDDNEWPSIPHDFKE
TMGEYRRELKKLAERVMEVMDENLGLEKGYIKRAFGGPEENPFFGTKVSHYPPCPRPDLVA
GLRAHTDAGGVILLFQDDEVGGLQILKDGQWIDVQPIKNAIVINTGDQLEVISNGRYKSVW
HRVLALHEGNRRSIASFYNPSLSATIEPAAKLLMHLEAENETTVDKVKYPRFVFGDYMQVY
QCQKFLPKEPRFQAVGAAAV
>sp|Q9ZUN4|ACCO1_ARATH 1-aminocyclopropane-1-carboxylate oxidase 1 OS=Arabidopsis thaliana OX=3702 GN=ACO1 PE=2 SV=1
MVLIKEREMEIPVIDFAELDGEKRSKTMSLLDHACDKWGFFMVDNHGIDKELMEKVKKMI
NSHYEEHLKEKFYQSEMVKALSEGKTSDADWESSFFISHKPTSNICQIPNISEELSKTMD
EYVCQLHKFAERLSKLMCENLGLDQEDIMNAFSGPKGPAFGTKVAKYPECPRPELMRGLR
EHTDAGGIILLLQDDQVPGLEFFKDGKWVPIPPSKNNTIFVNTGDQLEILSNGRYKSVVH
RVMTVKHGSRLSIATFYNPAGDAIISPAPKLLYPSGYRFQDYLKLYSTTKFGDKGPRLET
MKKMGNADSA
>sp|Q41931|ACCO2_ARATH 1-aminocyclopropane-1-carboxylate oxidase 2 OS=Arabidopsis thaliana OX=3702 GN=ACO2 PE=1 SV=2
MEKNMKFPVVDLSKLNGEERDQTMALINEACENWGFFEIVNHGLPHDLMDKIEKMTKDHY
KTCQEQKFNDMLKSKGLDNLETEVEDVDWESTFYVRHLPQSNLNDISDVSDEYRTAMKDF
GKRLENLAEDLLDLLCENLGLEKGYLKKVFHGTKGPTFGTKVSNYPPCPKPEMIKGLRAH
TDAGGIILLFQDDKVSGLQLLKDGDWIDVPPLNHSIVINLGDQLEVITNGKYKSVLHRVV
TQQEGNRMSVASFYNPGSDAEISPATSLVEKDSEYPSFVFDDYMKLYAGVKFQPKEPRFA
AMKNASAVTELNPTAAVETF
>sp|O65378|ACCO3_ARATH 1-aminocyclopropane-1-carboxylate oxidase 3 OS=Arabidopsis thaliana OX=3702 GN=At1g12010 PE=1 SV=1
MEMNIKFPVIDLSKLNGEERDQTMALIDDACQNWGFFELVNHGLPYDLMDNIERMTKEHY
KKHMEQKFKEMLRSKGLDTLETEVEDVDWESTFYLHHLPQSNLYDIPDMSNEYRLAMKDF
GKRLEILAEELLDLLCENLGLEKGYLKKVFHGTTGPTFATKLSNYPPCPKPEMIKGLRAH
TDAGGLILLFQDDKVSGLQLLKDGDWVDVPPLKHSIVINLGDQLEVITNGKYKSVMHRVM
TQKEGNRMSIASFYNPGSDAEISPATSLVDKDSKYPSFVFDDYMKLYAGLKFQAKEPRFE
AMKNAEAAADLNPVAVVETF
>sp|Q06588|ACCO4_ARATH 1-aminocyclopropane-1-carboxylate oxidase 4 OS=Arabidopsis thaliana OX=3702 GN=ACO4 PE=1 SV=2
MESFPIINLEKLNGEERAITMEKIKDACENWGFFECVNHGISLELLDKVEKMTKEHYKKC
MEERFKESIKNRGLDSLRSEVNDVDWESTFYLKHLPVSNISDVPDLDDDYRTLMKDFAGK
IEKLSEELLDLLCENLGLEKGYLKKVFYGSKRPTFGTKVSNYPPCPNPDLVKGLRAHTDA
GGIILLFQDDKVSGLQLLKDGEWVDVPPVKHSIVVNLGDQLEVITNGKYKSVEHRVLSQT

DGEGRMSIASFYNPGSDSVIFPAPELIGKEAEKEKKENYPRFVFEDYMKLYSAVKFQAKE
PRFEAMKAMETTVANNVGPLATA
>sp|Q0WPW4|ACCO5_ARATH 1-aminocyclopropane-1-carboxylate oxidase 5 OS=Arabidopsis thaliana OX=3702 GN=At1g77330 PE=2 SV=1
MAIPVIDFSKLNGEEREKTLSEIARACEEWGFFQLVNHGIPLELLNKVKKLSSDCYKTER
EEAFKTSNPVKLLNELVQKNSGEKLENVDWEDVFTLLDHNQNEWPSNIKETMGEYREEVR
KLASKMMEVMDENLGLPKGYIKKAFNEGMEDGEETAFFGTKVSHYPPCPHPELVNGLRAH
TDAGGVVLLFQDDEYDGLQVLKDGEWIDVQPLPNAIVINTGDQIEVLSNGRYKSAWHRVL
AREEGNRRSIASFYNPSYKAAIGPAAVAEEEGSEKKYPKFVFGDYMDVYANQKFMPKEPR
FLAVKSL
>XP_003519400.1 1-aminocyclopropane-1-carboxylate oxidase 2 [Glycine max]
MASSDLFGGATSAPPPTPAAHHDNIMSSSDAADALSRLLQRLPPTLSLPTRLPSPSSAAAT
CPPCLSLNDVLSCVSKLGYAQLTDHSVPSELANSAESEALALFDLSQDQKQSLFPKNWPLG
YGNDEDEDEDGVADSFRFDSACSTESSELALFSLRKFARELEKLGLMIVDELTKDLGCENP
LGDDPTRVCSVMWVSESLPGNKSGGFYPFVIGLQYQIRNQKYSLLSDSGWVSVLPHVDSIL
VTFGDIAQVWSNGKLKKVRGRPMATVGDENGSRCITMSLLITLPTDSRVAPLLPKVTCNKD
QKEEEIEGEEEENNDGGDEELEKRVFNSFDFEDYAWRVYHERILFKDPLDRYRVV
>XP_003523026.1 1-aminocyclopropane-1-carboxylate oxidase 1 [Glycine max]
MGEACIPTVDLSPFLREDEDGKKRAIEAITQACSEYGFFQIVNHGVSLDLVKEAMQQSKTF
FDYSDEEKSKSSPSSDAPLPAGYSRQPLHSPDKNEYFLFFSPGSSFNVIPQIPPKFRDVLE
EMFVQMSKMGVLLESIINECLGLPTNFLKEFNHDRSWDFLVALRYFPASNNENNGITEHED
GNIVTFVVQDGVGGLQVLKNGDWVPVVPAEGTIVVNVGDVIQVLSNNKFKSATHRVVRAEG
RSRYSYVFFHNLRGDKWVEPLPQFTSDIGEPPKYRGFLYKEYQELRMRNKSHPPSRPEDEI
HITHYAIDN
>XP_003525310.1 1-aminocyclopropane-1-carboxylate oxidase 1 [Glycine max]
MEIPVIDFSKLNGDKRGDTMALLHEACEKWGCFMVENHEIDTQLMGKVKQLINAYYEENLK
ESFYQSEIAKRLEKQQNTSDIDWESTFFIWHRPTSNINEISNISQELCQTMDEYIAQLLKL
GEKLSELMSENLGLEKDYIKKAFSGNGEGPAVGTKVAKYPQCPRPELVRGLREHTDAGGII
LLLQDDEVPGLEFFKDGKWVEIPPSKNNAIFVNTGDQVEVLSNGLYRSVVHRVMPDNNGSR
ISIATFYNPIGDAIISPAPKLLYPSNFRYGDYLKLYGSTKFGEKAPRFESMKNMINGHKII
PA
>XP_003530544.1 1-aminocyclopropane-1-carboxylate oxidase 1 [Glycine max]
MEIPVIDFSNLNGDKRGDTMALLHEACEKWGCFMVENHEIDTQLMEKLKQLINTYYEEDLK
ESFYQSEIAKRLEKQQNTSDIDWEITFFIWHRPTSNINEIPNISRELCQTMDEYIAQLLKL
GEKLSELMSENLGLEKDYIKKAFSGSGEGPAVGTKVAKYPQCPRPELVRGLREHTDAGGII
LLLQDDKVPGLEFFKDGKWVEIPPPKNNAVFVNTGDQVEVLSNGLYKSVLHRVMPDNSGSR
TSIATFYNPIGDAIISPAPKLLYPSNFRYGDYLKLYGSTKFGEKAPRFECMKNMTNGHKNI
PA
>XP_025985012.1 1-aminocyclopropane-1-carboxylate oxidase 1 [Glycine max]
MAIPVIDFSTLNGDKRGETMALLDEACQKWGFFLIENHEIDKNLMEKVKELINIHYEENLK
EGFYQSEIAKTLEKKQNTSDIDWESAFFIWHRPTSNIKKITNISQELCQTMDQYIDQLVTL
AEKLSELMSENLGLEKNYIKEAFSGTNGPAMGTKVAKYPQCPHPELVRGLREHTDAGGIIL
LLQDDQVPGLEFFKDGKWVEIPPSKNNAIFVNTGDQVEVLSNGFYKSVVHRVMPDKNGSRL
SIASFYNPVGEAIISPANKLLYPSNYRYGDYLELYGNTKFGEKGPRFESIKNMTNGHCNLK
P
>NP_001341820.2 putative 1-aminocyclopropane-1-carboxylate oxidase [Glycine max]
MTNFPVINLEKLNGEERNDTMEKIKDACENWGFFELVNHGIPHDLLDTVERLTKEHYRKCM
EERFKEFMASKGLDAVQTEVKDMDWESTFHLRHLPESNISEIPDLIDEYRKVMKDFALRLE
KLAEQLLDLLCENLGLEKGYLKKAFYGSRGPTFGTKVANYPPCPNPDLVKGLRPHTDAGGI

VLLFQDDKVSGLQLLKDGQWVDVPPMRHSIVVNIGDQLEVITNGKYRSVEHRVIAQTDGTR
MSIASFYNPGSDAVIYPAPELLEKEAEEKSQLYPKFVFEDYMKLYAKLKFQAKEPRFEAFK
ASNFGPIATV
>XP_003519448.1 1-aminocyclopropane-1-carboxylate oxidase [Glycine max]
MTNFPLINLEKLSGEERNDTMEKIKDACENWGFFELVNHGIPHDILDTVERLTKEHYRKCM
EERFKELVASKGLDAVQTEVKDMDWESTFHLRHLPESNISEIPDLIDEYRKVMKDFALRLE
KLAEQLLDLLCENLGLEKGYLKKAFYGSRGPTFGTKVANYPPCPNPELVKGLRPHTDAGGI
ILLFQDDKVSGLQLLKDGQWVDVPPMRHSIVVNIGDQLEVITNGKYKSVEHRVIAQTDGTR
MSIASFYNPGSDAVIYPAPELLEKEAEEKNQLYPKFVFEDYMKLYAKLKFQAKEPRFEAFK
ASNFGPIATV
>XP_003533043.2 1-aminocyclopropane-1-carboxylate oxidase [Glycine max]
MQSPLTPMPDITVDFRAPPPSPVASGRSSTVTNDDVLTEFLEASLRVPDLVLPDKIFPKQN
HLEAPPEVDFVSLCFHCDDALRDIVSDSLARIGCFQLLNHGIPLQLMTAVAEAAAGIFQVP
PANRVSATRSPEKPWGFEEYHAGEEEGDGSEEFVWCNDHELKSKMDGIWPIGYPNFSEKME
KLKSRIEMVGRKMLGVILKKGSIEFVGSGHEVGTLCCVYKHRRDDHSISSSLKYDVIRMLI
RGTDYSHSLCFHVCNGSSQFHVYSKKSWLSFFPQPGALIVTAGDQTQTFSGGEYKHVIGRP
IFKGEKEETISMAFLYSTQTTKDNFQTSRGRTISLGQQAILALILPLVYHVINFFYKKLN

>XP_003545587.1 1-aminocyclopropane-1-carboxylate oxidase [Glycine max]
MENFPVINLENLNGEERKATLNQIEDACQNWGFFELVSHGIPLELLDTVERLTKEHYRKCM
EKRFKEAVSSKGLEAEVKDMDWESTFFLRHLPTSNISEIPDLSQEYRDAMKEFAQKLEKLA
EELLDLLCENLGLEKGYLKNAFYGSRGPNFGTKVANYPACPKPELVKGLRAHTDAGGIILL
LQDDKVSGLQLLKNGQWVDVPPMRHSIVVNLGDQIEVITNGRYKSVEHRVIAQTNGTRMSV
ASFYNPASDALIYPAPVLLEQKAEDTEQVYPKFVFEDYMKLYATLKFQPKEPRFEAMKAIG
SF
>XP_006584893.1 1-aminocyclopropane-1-carboxylate oxidase [Glycine max]
MENFPVINLENLNGEERKTILEQIEDACENWGFFELVNHGIPHELLDIVERLTKEHYRKCM
EQRFKEAVASKGLEGIQAEVKDMNWESTFFLRHLPDSNISQIPDLSEEYRKVMKEFAQKLE
KLAEKLLDLLCENLGLEKGYLKKVFYGSKGPNFGTKVANYPPCPNPELVKGLRAHTDAGGI
ILLLQDDKVSGLQLLKDGHWVDVPPMRHSIVVNLGDQLEVITNGRYKSVELRVIARTDGTR
MSIASFYNPASDAVIYPAPALLDSKAEETDKVYPKFVFEDYMRLYATLKFQPKEPRFQAMK
EVNSF
>XP_003549632.1 1-aminocyclopropane-1-carboxylate oxidase [Glycine max]
MENFPVVDMGNLNNEERSATMEIIKDACENWGFFELVNHGISIELMMDTVERMTKEHYKKC
MEQRFQEMVASKGLESAQSEINDLDWESTFFLRHLPVSNISEIPDLDEDYRKVMKDFAVEL
EKLAELVLELLCENLGLEKGYLKKVFCGSKGPNFGTKVSNYPPCPKPELIKGLRAHTDAGG
IILLFQDHKVSGLQLLKDAHWIDVPPMRHSIVINLGDQLEVITNGKYKSVMHRVITQTDGN
RMSIASFYNPGNDALIAPAPALVKEDETSQVYPKFVFDDYMKLYAGLKFQDKEPRFEAMKA
TESSNINLGPIATV
>XP_003529653.1 1-aminocyclopropane-1-carboxylate oxidase [Glycine max]
MEKFPVVDMGNLNNEERSATMEIIKDACENWGFFELVNHGISIELMDTVERMTKEHYKKCM
EQRFKEMVASKGLESAQSEINDLDWESTFFLRHLPASNISEIPDLDEDYRKVMKDFAVELE
ELAELVLDLLCENLGLEKGYLKKVFYGSKGPNFGTKVSNYPPCPKPELIKGLRAHTDAGGI
ILLFQDHKVSGLQLLKDGHWIDVLPMRHSIVINLGDQLEVITNGKYKSVMHRVITQTDGNR
MSIASFYNPGNDALIAPAPALVKEDETSQVYPKFVFDDYMKLYAGLKFQAKEPRFQAMKAT
ESSNINLGPIATV
>XP_003534723.1 1-aminocyclopropane-1-carboxylate oxidase [Glycine max]
MANFPVVDMGKLNTEERPAAMEIIKDACENWGFFELVNHGISIELMDTVEKLTKEHYKKTM
EQRFKEMVTSKGLESVQSEINDLDWESTFFLRHLPLSNVSDNADLDQDYRKTMKKFALELE

KLAEQLLDLLCENLGLEKGYLKKVFYGSKGPNFGTKVSNYPPCPTPDLIKGLRAHTDAGGI
ILLFQDDKVSGLQLLKDDQWIDVPPMRHSIVINLGDQLEVITNGKYKSVMHRVIAQTDGTR
MSIASFYNPGDDAVISPAPALVKELDETSQVYPKFVFDDYMKLYAGLKFQAKEPRFEAMKA
NASVVDVGAIATV
>lcl|AFJ75134.1 ACC oxidase, partial [Malus domestica]
DLSLVNGEERAATLEKINDACENWGFFELVNHGMSTELLDTVEKMTKDHYKKTMEQRFKEM
VAAKGLDDVQSEIHDLDWESTFFLRHLPSSNISEIPDLEEDYRKTMKEFAVELEKLAEKLL
DLLCENLGLEKGYLKKVFYGSKGPNFGTKVSNYPPCPKPDLIKGLRAHSDAGGIILLFQDD
KVSGLQLLKDGEWVDVPPMHHSIVINLGDQIEVITNGKYKSVMHRVIAQSDGTRMSIASFY
NPGNDAFISPAPAVLEKKTGDAPTYPKFVFDDYMKLYSGLKFQAKEPRFEAMKAKES
>lcl|XP_008336897.3 1-aminocyclopropane-1-carboxylate oxidase
[Malus domestica]
MAIPVIDFSKLDGEERAKTLAEVANGCEDWGFFQLVNHGISEELLERVKKVTGDFFRMERE
ENFKNSTLMKAVNENKKLENVDWEDVFTLLDDNEWPSKTPGFKETMEEYRRELKKLAERVM
EVMDENLGLPKGYMKKAFNGGEEDNAFFGTKVSHYPPCPKPELVTGLRAHTDAGGVILLFQ
DDKVGGLQILKDGQWIEVQPLRNSIVINTGDQIEVLSNGRYKSVWHRVLATPTGNRRSIAS
FYNPSMKATIAPAPELVEKENQEVGQTYPKFVFGDYLSVYAEQKFLPKEPRFQSVRAV
>lcl|XP_008337682.2 1-aminocyclopropane-1-carboxylate oxidase
1 [Malus domestica]
MEIPVIDFNELNGEARSKTMALLDQACEKWGFFQVENHGIDKKLMDKVKQLINEYYEENLR
ASFYKSEIAKSLDQEVTSKVDWESSFFIWHRPTSNIEGIPNFSEDHCKTMNEYIAQLIKMA
EKLSELMCENLGLEKGHIKDAFSGGKGPSVGTKVAKYPQCPQPELVRGLREHTDAGGIILL
LQDDQVPGLEFLKDGEWFAIPPSKNNTMFVNIGDQVEVLSNGRYKSVLHRVLADKNGSRLS
IATFYNPAGDAIISPAPKLLYPNHLRFQDYLKHYATTKFSDKGLRFETMKQMADGHNGFLS

>lcl|NP_001281047.1 1-aminocyclopropane-1-carboxylate oxidase
1 [Malus domestica]
MATFPVVDLSLVNGEERAATLEKINDACENWGFFELVNHGMSTELLDTVEKMTKDHYKKTM
EQRFKEMVAAKGLDDVQSEIHDLDWESTFFLRHLPSSNISEIPDLEEEYRKTMKEFAVELE
KLAEKLLDLLCENLGLEKGYLKKVFYGSKGPNFGTKVSNYPPCPKPDLIKGLRAHSDAGGI
ILLFQDDKVSGLQLLKDGEWVDVPLMHQLTHILGDQIEVITNGKYKSVMHRVIAQSDGTRM
SIASFYNPGNDSFISPAPAVLEKKTEDAPTYPKFVFDDYMKLYSGLKFQAKEPRFEAMKAK
ESTPVATA
>lcl|Q00985.1 RecName: Full=1-aminocyclopropane-1-carboxylate
oxidase 1; Short=ACC oxidase 1; AltName: Full=Ethylene-
forming enzyme; Short=EFE; AltName: Full=PAE12; AltName:
Full=Protein AP4 [Malus domestica]
MATFPVVDLSLVNGEERAATLEKINDACENWGFFELVNHGMSTELLDTVEKMTKDHYKKTM
EQRFKEMVAAKGLDDVQSEIHDLDWESTFFLRHLPSSNISEIPDLEEEYRKTMKEFAVELE
KLAEKLLDLLCENLGLEKGYLKKVFYGSKGPNFGTKVSNYPPCPKPDLIKGLRAHSDAGGI
ILLFQDDKVSGLQLLKDGEWVDVPPMHHSIVINLGDQIEVITNGKYKSVMHRVIAQSDGTR
MSIASFYNPGNDSFISPAPAVLEKKTEDAPTYPKFVFDDYMKLYSGLKFQAKEPRFEAMKA
KESTPVATA
>lcl|XP_008342143.2 1-aminocyclopropane-1-carboxylate oxidase
[Malus domestica] 3
MENFPVINLESLNGEGRKATMEKIKDACENWGFFELVSHGIPTEFLDTVERLTKEHYRQCM
EQRFKELVASKGLDAVQTEVNDMDWESTFYLRHLPQSNISEVPDLKDEYRNVMKEFALKLE
KLAEQLLGLLCENLGLEQGYLTKAFYGTNGPTFGTKVSNYPPCPNNDKIKGLRAHTDAGGL
ILLFQDDKVSGLQLLKDGEWVDVPPMRHSLVINLGDQLEVITNGKYKSVEHRVIAQTDGTR
MSIASFYNPSSDAVIYPAPTLVEKKAVEKNQVYPKFVFEDYMKHYVGVKFQAKEPRFEAMK
AVEVKASFGLSPVKTTA
>lcl|NP_001391910.1 1-aminocyclopropane-1-carboxylate oxidase
2 [Malus domestica] 2
MATFPVVDMDLINGEERAATLEKINDACENWGFFELVNHGISTELLDTVEKMNKDHYKKTM
EQRFKEMVAAKGLEAVQSEIHDLDWESTFFLRHLPSSNISEIPDLEEDYRKTMKEFAVELE

KLAEKLLDLLCENLGLEKGYLKKAFYGSKGPNFGTKVSNYPPCPKPDLIKGLRAHTDAGGI
ILLFQDDKVSGLQLLKDGEWMDVPPVHHSIVINLGDQIEVITNGKYKSIMHRVIAQSDGTR
MSIASFYNPGDDAFISPAPALLEKKSEETPTYPKFLFDDYMKLYSGLKFQAKEPRFEAMKA
RETTPVETARGLRAVRWNTTKRNQN
>PTQ44570.1 hypothetical protein MARPO_0019s0012 [Marchantia polymorpha]
MAAMTMTRPNDPFTSDIARFHEVKGAKDEPFAELTRYHEVRGVREMVESFRVREVPSYYVK
PVNERRFTPPSAAVLSMEQQIPCIDLEALSGQELLSAIANACRDWGFFQVLNHGLPSQLVQ
NMAKQSSEFFAQPLEEKMKCSTPARVSGPVHFGGGGNRDWRDVLKLNCAPASIVAKEYWPQ
RPAGFRDTMEEYSSQQQALAIRLLKLISESLGLESNYLVAACGEPKVVMAINHYPPCPDPS
LTMGIKAHSDPNTITMLLQDDVGGLQVFKEDRWIDVRPLPNALVINVGDQLQILSNGKYSS
CLHRVVNNNRQARTSIATFFSPAHACIIGPAPGLVDEVNPAIYPNIVYADYIKAFYTQALG
PNNKNGGYLAGIELHRRYNCYTSSSSISS
>PTQ39739.1 hypothetical protein MARPO_0043s0013 [Marchantia polymorpha]
MSCPCCRPGGDSSDGEEEEEEEEDVGLLDMSAMKDRVQYLVDSGISQIPAAFIRPVDERPL
GVKGFDRIPVVDMSEVDGKGRERIRAEVGKACEEWGFFQVVNHGVKKSVLDVMRKDGRAFF
QQPMEQKLKYSCTPGVIASEGYGSKMLTKDEQLLDWRDYYDLHTLPMSRRRSGNWPTNPPS
FRDSVVKYSDQMKWLAEKILELISESLGLPGTYLKEALGEVSQNVSVNYYPVCPQPDLTLG
LQAHSDLGAITILMQADIPGLQVKKDDSWIAVEPVEDALVVNLGDMMEVLTNGRYKSVEHR
AVVNSERDRLSIATFLDPAKDRTLSPAKALVDDKHPLLYRDVLFSEFISAWYSKGPDGKRN
IDNLILPL
>BAR72291.1 Aminocyclopropanecarboxylate oxidase [Nicotiana benthamiana]
MESFPVVNMELLNTEQRAATMEKIKDACENWGFFEVVNHGICHELLDTVEKLTKGHYKKCM
EQRFKEMVASKGLEAVETEIKDLDWESTFFLKHLPVSNISEVPDLEDEYRKIMKEFAEKLE
KLAEQLLDLLCENLGLEQGYLKKAFYGSNGPTFGTKVSNYPPCPKPDLIKGLRAHTDAGGI
ILLFQDDKVSGLQLLKDDKWIDVPPMRHSIVINLGDQLEVITNGKYKSVEHRVIAQPDGNR
MSLASFYNPGSDAVIYPAPELLEKENKVIYPKFVFEDYMKLYAGLKFQAKEPRFEAMKAVE
TTVNSAPIATV
>BAR72293.1 Aminocyclopropanecarboxylate oxidase [Nicotiana benthamiana]
MENFPIINLEKLNGSEKAATMEMIKDACENWGFFELVNHGIPHEVMDTVEKLTKRHYKKCM
EQRFKKLVASNGLEGVQAEVTDMDWESTFFLRHLPVSNISEVPDLDDRYREVMRDFAKRLE
NLAEELLDLLCENLGLEKGYLKKVFYGSKGPNFGTKVSNYPPCPKPDLIKGLRAHTDAGGI
ILLFQDDKVSGLQLLKDGQWIDVPPMRHSIVVNLGDQLEVITNGKYKSVMHRVIAQKDGTR
MSLASFYNPGSDAVIYPAPTLVEKDAAESKQVYPKFVFDDYMKLYAGLKFQAKEPRFEAMK
SIESDVKLDPIATA
>BAR72288.1 Aminocyclopropanecarboxylate oxidase [Nicotiana benthamiana]
MENFPIINLEKLNGSEKAATMEMIKDACENWGFFELVNHGIPHEVMDTVEKLTKGHYKKCM
EQRFKELVASKGLEGVQAEVTDMDWESTFFLRHLPVSNISEVPDLDDQYREVMRDFAKRLE
NLAEELLDLLCENLGLEKGYLKKVFYGSKGPNFGTKVSNYPPCPKPDLIKGLRAHTDAGGI
ILLFQDDKVSGLQLLKDGQWIDVPPMRHSIVVNLGDQLEVITNGKYKSVMHRVIAQKDGTR
MSLASFYNPGSDAVIYPAPALVEKEAVESKQVYPKFVFDDYMKLYAGLKFQAKEPRFEAMK
SIESDVKMDAIATA
>sp|Q08506.1|ACCO1_PETHY RecName: Full=1-aminocyclopropane-1-carboxylate oxidase 1; Short=ACC oxidase 1; Short=ACCO; AltName: Full=Ethylene-forming enzyme; Short=EFE
MENFPIISLDKVNGVERAATMEMIKDACENWGFFELVNHGIPREVMDTVEKMTKGHYKKCM
EQRFKELVASKALEGVQAEVTDMDWESTFFLKHLPISNISEVPDLDEEYREVMRDFAKRLE
KLAEELLDLLCENLGLEKGYLKNAFYGSKGPNFGTKVSNYPPCPKPDLIKGLRAHTDAGGI
ILLFQDDKVSGLQLLKDGQWIDVPPMRHSIVVNLGDQLEVITNGKYKSVMHRVIAQKDGAR
MSLASFYNPGSDAVIYPAPALVEKEAEENKQVYPKFVFDDYMKLYAGLKFQAKEPRFEAMK
AMETDVKMDPIATV

>sp|Q08507.1|ACCO3_PETHY RecName: Full=1-aminocyclopropane-1-carboxylate oxidase 3; Short=ACC oxidase 3; AltName: Full=Ethylene-forming enzyme; Short=EFE
MENFPIINLEKLNGSERDATMEMIKDACENWGFFELVNHGIPHEVMDTVEKLTKGHYKKCM
EQRFKELVASKGLEAVQAEVTDLDWESTFFLRHLPVSNISEVPDLDDEYREVMRDFAKRLE
KLAEELLDLLCENLGLEKGYLKKAFYGSKGPNFGTKVSNYPPCPKPDLIKGLRAHTDAGGI
ILLFQDDKVSGLQLLKDGQWIDVPPMRHSIVVNLGDQLEVITNGKYKSVLHRVIAQTDGTR
MSLASFYNPGSDAVIYPAPTLVEKEADQECKQVYPKFVFDDYMKLYAGLKFQAKEPRFEAM
KAREADVKSDPIATA
>sp|Q08508.1|ACCO4_PETHY RecName: Full=1-aminocyclopropane-1-carboxylate oxidase 4; Short=ACC oxidase 4; AltName: Full=Ethylene-forming enzyme; Short=EFE
MENFPIINLENLCGAERDATMEMIKDACENWGFFELVNHGIPHEVMDTVEKFTKGHYKKCM
EQRFKELVASKGLEAVQAEVTDLDWESTFFLRHLPVSNISEVPDLDDEYREVMRDFAKRLE
KLAEELLDLLCENLGLEKGYLKKAFYGSKGPNFGTKVSNYPPCPKPDLIKGLRAHTDAGGI
ILLFQDDKVSGLQLLKDDQWIDVPPMRHSIVINLGDQLEVITNGKYKSVPHRVIAQTDGTR
MSLASFYNPASDAVIYPAPALVERDAEENKQIYPKFVFDDYMKLYARLKFQAKEPRFEAMK
AMEADVKIDPVATV
>NP_001233867.2 1-aminocyclopropane-1-carboxylate oxidase [Solanum lycopersicum]
MESNFPVVDMGLLQTEKRPEAMDKIKDACENWGFFELVNHGISHELLDTVENLTKGHYKKC
MEQRFKEMVASKGLEAVQTEIDDLDWESTFFLKHLPVSNVYEVPDLDDEYRKVMKDFALKL
EKLAENLLDLLCENLGLEKGYLKKAFYGSKGPTFGTKVSNYPPCPKPDLIKGLRAHTDAGG
IILLFQDDKVSGLQLLKDGNWIDVPPMKHSIVINLGDQLEVITNGRYKSIEHRVIAQQDGT
RMSIASFYNPGSDAVIFPAPELIEKTEEDNKLKYPKFVFEDYMKLYAGLKFQAKEPRFEAM
KAVETTVNLGPIETV
>NP_001234638.1 1-aminocyclopropane-1-carboxylate oxidase [Solanum lycopersicum]
METFPVVNMEMLNTEKRAAALEKIKDACENWGFFEVINHGISHELLDTVEKFTKEHYKKCM
EQRFKEMVASKGLEGVQTEIDDLDWESTFFLKHLPVSNISEVPDLEDDYRKIMKEFADKLE
KLAEQLLDLLCENLGLEQGYLKKVFYGSKGPTFGTKVSNYPPCPKPDLIKGLRAHTDAGGI
ILLFQDDKVSGLQLLKDDKWIDVPPMRHSIVINLGDQLEVITNGKYKSVEHRVIAQPDGNR
MSLASFYNPGSDAVIYPAPELLEKEEKENTIMYPKFVFEDYMKLYAGLKFQAKEPRFEAMK
AVETAVNLGPIATV
>NP_001234037.1 1-aminocyclopropane-1-carboxylate oxidase [Solanum lycopersicum]
MEMPVIDFSKLEGEERCATMSLLHQACEKWGFFMIENHGIDSYLMDNVKQFVNQHYEANMK
KRFYESELPMSLEKNGNISNTDWESTFFVWHRPASNIYEIQGLSKELCKAVDDYIDQLIKL
AENLSELMCENLGLAKSYIKEAFSGSKGPSVGTKVAIYPQCPRPDLVRGLREHTDAGGIIL
LLQDEQVPGLEFFKDGHWVNIPPSKNNRLFVNIGDQIEILTNGMYKSIRHRVMAEKDGNRL
SIATFYNPAGEAIISPASKLLYPCHLRFQDYLNLYSKTKFAEKGPRFESAKRLANGH
>NP_001316842.1 1-aminocyclopropane-1-carboxylate oxidase 2 [Solanum lycopersicum]
MENFPIINLEKLNGAERVATMEKINDACENWGFFELVNHGIPHEVMDTVEKLTKGHYKKCM
EQRFKELVAKKGLEGVEVEVTDMDWESTFFLRHLPSSNISQLPDLDDVYREVMRDFAKRLE
KLAEELLDLLCENLGLEKSYLKNTFYGSKGPNFGTKVSNYPPCPKPDLIKGLRAHTDAGGI
ILLFQDDKVSGLQLLKDGRWIDVPPMRHSIVVNLGDQLEVITNGKYKSVMHRVIAQKDGTR
MSLASFYNPGNDALIYPAPALVDKEAEEHNKQVYPKFMFDDYMKLYANLKFQAKEPRFEAM
KAMESDPIAIA
>NP_001234024.2 1-aminocyclopropane-1-carboxylate oxidase 1 [Solanum lycopersicum]
MENFPIINLEKLNGDERANTMEMIKDACENWGFFELVNHGIPHEVMDTVEKMTKGHYKKCM
EQRFKELVASKGLEAVQAEVTDLDWESTFFLRHLPTSNISQVPDLDEEYREVMRDFAKRLE
KLAEELLDLLCENLGLEKGYLKNAFYGSKGPNFGTKVSNYPPCPKPDLIKGLRAHTDAGGI
ILLFQDDKVSGLQLLKDEQWIDVPPMRHSIVVNLGDQLEVITNGKYKSVLHRVIAQTDGTR

MSLASFYNPGSDAVIYPAKTLVEKEAEESTQVYPKFVFDDYMKLYAGLKFQAKEPRFEAMK
AMESDPIASA
>NP_001233928.1 1-aminocyclopropane-1-carboxylate oxidase 4
[Solanum lycopersicum]
MENFPIINLENLNGDERAKTMEMIKDACENWGFFELVNHGIPHEVMDTVEKLTKGHYKKCM
EQRFKELVASKGLEAVQAEVTDLDWESTFFLRHLPTSNISQVPDLDEEYREVMRDFAKRLE
KLAEELLDLLCENLGLEKGYLKNAFYGSKGPNFGTKVSNYPPCPKPDLIKGLRAHTDAGGI
ILLFQDDKVSGLQLLKDEQWIDVPPMRHSIVVNLGDQLEVITNGKYKSVMHRVIAQTDGTR
MSLASFYNPGNDAVIYPAPSLIEESKQVYPKFVFDDYMKLYAGLKFQPKEPRFEAMKAMEA
NVELVDQIASA
>XP_004242046.1 1-aminocyclopropane-1-carboxylate oxidase 5
[Solanum lycopersicum]
MTTIPVIDFSKLYGEERAQTLAQISKGCEEWGFFQLVNHGIPVELLERVKKVGEECFKLER
EEVFNKSRVVNLLNEFVESNKNGKVENVDWEDVFLLTDDNCNQWPSKTPQFKETMKEYRSE
VKKLAERVMEVMDENLGLSKGYIKKAFNNNNNNDAFFGTKVSHYPPCPHPEKVNGLRAHTD
AGGVILLFQDDQVDGLQILKNGQWIDVPPIPNAIVINTGDQIEVLSNGKYKSVWHRVLSKP
DGNRRSIASFYNPSLKATISPAPELLILEEKKVDQLIATYPTFVFGNYMNVYNEQKFLPKE
PRFQAVAAI
>XP_044389578.1 1-aminocyclopropane-1-carboxylate oxidase-
like [Triticum aestivum]
MVVPVIDFSKLDGAERAETMAQIADGCENWGFFQLVNHGIPLELLDRVKKVCSESYRLREA
AFRSSEPVQTLERLVETERRGEAVAPVDDMDWEDIFYLHDDNQWPSDPPAFKETMREYRAE
LKKLAERVMEAMDENLGLDKGRMKAAFTGDGLHAPFFGTKVSHYPPCPRPDLITGLRAHTD
AGGVILLFQDDKVGGLEVLKDGEWLDVQPLPDAIVVNTGDQVEVLSNGRYRSAWHRVLPMR
NGNRRSIASFYNPAFEAAISPAVGEGAAAAYPDYVFGDYMDVYNKQKFDAKEPRFEAVKAP
KAA
>XP_044377701.1 1-aminocyclopropane-1-carboxylate oxidase-
like [Triticum aestivum]
MAIPVIDFSRLDGDERAAALAEIAAGFEQWGFFQLVNTGIPDELLERVKKVCSDCYKLREE
GFNGSNPAVKALAALVEREGEGLAPRKVEGMDWEDVFTLHDDLPWPSIPPTFKETMMEYRT
ELKKLAEKLLGVMEELLGLAEGQITKVFSKDGDFKPFYGTKVSHYPPCPRPEMVDGLRAHT
DAGGLILLFQDDRVGGLQVLGRDGRWADVQPVENAIVINTGDQIEVMSNGRYKSAWHRVLA
TRDGNRRSIASFYNPARAATIAPAIPAAADSGTGGDYPSFSFGDYMEVYIKQKFQDKEPRF
AAAAAIKTTVD
>XP_044404490.1 1-aminocyclopropane-1-carboxylate oxidase-
like isoform X1 [Triticum aestivum]
MALPANAAASLSFPVIHMEKLETGERGATMGVIRDACENWGFFELLNHGISAELMDEVERV
SKAHYAACREEQFKEFAAKTLEAGEKGDDVKDVDWESTFFVRHLPASNLADLPDLDHHYRQ
VMKEFAAEIEKLAEKVLDLLCENLGLEQGYLKRAFAGSRGPTFGTKVSSYPPCPRPDLVAG
LRAHTDAGGVILLFQDDQVSGLQLLKDGAWVDVPPMRHAIVVNIGDQLEVITNGRYKSVMH
RVLTRPDGNRMSLASFYNPGADAVIFPAPALVEELSEEEAERAGSAVYPRFVFEDYMNLYL
LHKFEAKEPRFQAMKADAAPIATA
>XP_044436804.1 1-aminocyclopropane-1-carboxylate oxidase 1-
like [Triticum aestivum]
MEIPVIKMDELHGEKRSETLSLLHDACAQWGFFWLENHGINDDLMDRTKELVNKHYEQNME
KNFYSSEIAKTVGPDKVTSNVDWECSFMYHHQPKSNIHDIPELLRTTVPEYAEEVVKLAEQ
LAEVMSENLGLDKDYLKKAFTKPSIGIKVAKYPRCSHPEVVMGLRGHTDAGGIILLFQDDL
VPGLQFMKDGRWISIPPTKGNRIFVNLGDQIEVISNGIYKSICHQVLPNQNGSRLSIATFY
NPGADAIIFPAPNLTYPSQYRFQDYLNFYSATKFTD KVSRFQTTKMVFK
>XP_044440539.1 1-aminocyclopropane-1-carboxylate oxidase 1-
like [Triticum aestivum]
MDMEIPVIDLQGLTGDASQRSQTMARLHEACKDWGFFWVDSHGVDAALMEEVKRFVYAHYD
EHLKDRFYASDLAKDLQLPAEESKTVSGEVDWETAYFIRHRPANNVADFPEIPPATREMLD
AYIGQMVSLAELLAECMSLNLGLDGGLVRDTFTPPFVGTKVAMYPACPRPDLVWGLRAHTD
AGGIILLLQDDVVGGLEFFRGDREWVPVGPTKGSRIFVNIGDQLEVMSGGAYRSVLHRVAA

VAEGRRLSVATFYNPGADAVVAPAATARQPAAQLYPGPYRFGDYLDYYQGTKFADKAARFQ
AFKELFGSRIRHD
>XP_044381569.1 1-aminocyclopropane-1-carboxylate oxidase 1-like [Triticum aestivum]
MAPASSFPIIDMGLLAGEERPAAMDLLHDACENWGFFQVLDHGISTELMDEVEKMTKGHYK
RVREQRFLEFASKTLQDGGRAAENLDWESTFFVRHLPEPNIAEIPDLDDDYRRVMKQFASE
LERLAERLLDLLCENLGLDKGYLTRAFRGSNGAPTFGTKVSSYPPCPRPDLVKGLRAHTDA
GGIILLFQDDRVGGLQLLRGGEWVDVPPTRHSIVVNLGDQLEVITNGRYKSVVHRVVAQAD
GNRMSIASFYNPAGDAVIFPAPALVEAEAAGGAYPRFVFEDYMKLYVRHKFEDKEPRFEAF
KSMDSHSSNLIATA
>XP_044397656.1 1-aminocyclopropane-1-carboxylate oxidase 1-like [Triticum aestivum]
MASTLSFPIIDMGLLRGEERPAAMNLLHDACQNWGFFQVLDHGISTELMDEVEKMTKEHYK
RVREQRFLEFASKTLQDGDKAAENLDWESTFFVRHLPEPNIAEIPDLDDEYRRVMKQFAAE
LERLAERLLDLLCENLGLEKGYLTRAFRGSKGVPTFGTKVSSYPPCPRPDLVKGLRAHTDA
GGIILLFQDDRVGGLQLLKDGEWVDVPPTRHSIVVNLGDQLEVITNGRYKSVLHRVVAQAD
GNRMSIASFYNPASDAVIFPAPALAEAAGGAYPRFVFEDYMKLYVRHKFEDKEPRFEAFKS
MGSQTSNPIATA
>XP_044397657.1 1-aminocyclopropane-1-carboxylate oxidase 1-like [Triticum aestivum]
MAPASSFPIIDMGLLAGEERPAAMDLLHDACENWGFFQVLDHGISTELMDEVEKMTKEHYK
RVREQRFLEFASKTLEDGGRAAENLDWESTFFVRHLPEPNIAEIPDLDEEYRRVMKQFAAE
LERLAERLLDLLCENLGLEKGYLTRAFRGSKGVPTFGTKVSSYPPCPRPDLVKGLRAHTDA
GGIILLFQDDRVGGLQLLRDGEWVDVPPTRHSIVVNLGDQLEVITNGRYKSVLHRVVAQAD
GNRMSIASFYNPASDAVIFPAPALAEAEAAGGAYPRFVFEDYMKLYVRHKFEDKEPRFEAF
KSMESQSTKLIATA
>XP_044404493.1 1-aminocyclopropane-1-carboxylate oxidase 3-like [Triticum aestivum]
MAIPADAAVSLNFPVINMEKLETGERGAAMEVIRDACENWGFFELLNHGISHELMDEVERV
SKANYTACREGQFKEFAARTLEAGEKGADVKDVDWESTFFVRHLPASNLADLPDLDHHYRQ
LIKEFASEIEKLAEKVLDLLCENLGLEQGYLKRAFAGSRGPTFGTKVSSYPPCPRPDLVDG
LRAHTDAGGVILLFQDDQVSGLQLLKDGAWVEVPPMRHAIVVNIGDQLEVITNGRYKSVMH
RVLTRADGNRMSIASFYNPGADAVIFPAPALVAAAGATEKNEGEEGGAVYPKFVFEDYMNL
YVRHKFEAKEPRFKAMKADAAPIATA
>XP_044409773.1 1-aminocyclopropane-1-carboxylate oxidase 1-like isoform X2 [Triticum aestivum]
MAIPANAAASLSFPVINMEKLETEERGAAMGVIGDACENWGFFEVPLPPTHIEFSSAGISH
ELMDEVERVSKAHYAACWEQQFKEFAARTLEAGEKGADVKDVDWESTFFVRHLPSSNLADL
PNLDHHYRQVMKEFASEIEKLAEKVLDLLCENLGLELGYLKRAFAGSRGPTFGTKVSSYPP
CPRPDLVDGLRAHTDAGGVILLFQDDQVSGLQLLKDGAWVDVSPMRHAIVVNIGDQLEVIT
NGRYKSVMHRVLTRPDGNRMSLASFYNPGADAVIFPAPALVEELSEEEAERAGSAVYPRFV
FEDYMNLYLRHKFESKEPRFEAMKADAAPIATA
>XP_044421053.1 1-aminocyclopropane-1-carboxylate oxidase 1-like [Triticum aestivum]
MGIPANATASFSFPVINMEKLETQERGAAMGVIGDACENWGFFELLNHGISHELLDEVERA
SKAHYAACREEQFKEFAAKTLEAGEKGADVKDVDWESTFFVRHLPASNLADLPDLDHHYRQ
VMKEFAAEIEKLAEKVLDLLCENLGLELGYLKQAFAGSWGPTFGTKVSSYPPCPRPDMVDG
LRAHTDAGGVIMLFQDDQVSGLQLLKDGAWVDVPPMRHAIVVNIGDQLEVITNGRYKSVMH
RVLTRPDGNRMSIASFYNPGADAVIFPATALVEEPSEEAERAGSAVYPRFVFEDYMNLYMR
HKFEAKEPRFEAMKADAAPIATA
>NP_001149209.1 1-aminocyclopropane-1-carboxylate oxidase [Zea mays]
MAPTAAKDSGSADRWREVQAFEDSKLGVKGLVDSGVKSIPAMFHHPPESLKDVISPPALPS
SDDAPAIPVVDLSVARREDLVAQVKHAAGTVGFFWVVNHGVPEELMASMLSGVRQFNEGSL
EAKQALYSRDPARNVRFASNFDLFESAAADWRDTLYCKIAPDPAPRELVPEPLRNVMMEYG
EELTKLARSMFELLSESLGMPSDHLHKMECMQQLHIVCQYYPPCPEPHLTIGVRKHSDTGF

FTILLQDGMGGLQVLVDRGGGRQTWVDVTPRPGALMVNMGSFLQLVTNDRYKSVDHRVPAN
KSSDTARVSVAAFFNPDEKRTERLYGPIPDPSKPPLYRSVTFPDFIAKFNSIGLDGRVLDH
FRLEDDGPTRLAAPAHHV
>NP_001151658.2 1-aminocyclopropane-1-carboxylate oxidase
[Zea mays]
MASSSLPAPAAGRAELLKAFDDARTGVRGLVESGVSSVPELFRHADPYASIPLAPPGVSIP
VVDLSLPPHLAAAAAASAARTWGFFHLVNHHHALPAAAAAADDDYPERAFAAVRAFNELPA
HERAPHYSRAVDGGVNYSSNVDLYNSPAASWRDTIQILLGPNRHPDLADRIPAACRAEVLE
WEVRATAVARALLRLLSQGLGLRPEALEDASCADGKLMVCHYYPHCPEPERTMGIVPHTDP
GVLTVLAQDGVGGLQVKHQDEDGKISWVDVKPVPGALVINVGDLLQIMSNDKYTSVEHRVV
MNTREEPRVSIGIFFSPGKRGDSVFYGPLPELVSSENPPKYRNFTMSEFYGTFFSRDLASK
ALLDNFKLSP
>XP_008669786.2 1-aminocyclopropane-1-carboxylate oxidase
[Zea mays]
MAAIPVIDFSKLEGSERAETMAAIAAGFEHVGFFQLVNTGIPDELLERVKKVCSDCYKLRD
EAFMDSNLAVKALAELVDKESEGGAPMRKIEGMDWEDVFTLHDDLPWPSNPPAFKETMMEY
RKELRKLAEKMLGVMEELLGLEEGHIRKAFTNDGELEPFYGTKVSHYPPCPRPDLVDGLRA
HTDAGGLILLFQDDRFGGLQAQLPDGSWVDVQPLDNAIVVNTGDQIEVLSNGRYKSAWHRI
LATRDGNRRSVASFYNPARLATIAPAIPAADNYPSFVFGDYMQVYVKQKFQAKTSRFAAMA
TTTTK
>NP_001130227.1 1-aminocyclopropane-1-carboxylate oxidase
1Acc oxidase [Zea mays]
MAATVSSFPVVNMEKLETEERATAMEVIRDGCENWGFFELLNHGISHELMDEVERLTKAHY
ATFREAKFQEFAARTLEAGEKGADVKDVDWESTFFVRHLPASNLADLPDVDDRYRQVMEQF
ASEIRKLSERLLDLLCENLGLEPGYLKAAFAGSDGPTFGTKVSAYPPCPRPDLVDGLRAHT
DAGGIVLLFQDDQVSGLQLLRGGEWVDVPPMRHAIVANVGDQLEVITNGRYKSVMHRVLTR
PDGNRMSVASFYNPGADAVIFPAPALVGAAEEDRAEAAYPSFVFEDYMNLYVRHKFEAKEP
RFEAMKSAIATA
>NP_001141367.2 1-aminocyclopropane-1-carboxylate oxidase
[Zea mays]
MVVPVIDFSKLDGAERAETLAQIANGCEEWGFFQLVNHGIPLELLERVKKVSSDCYRLREA
GFKASEPVRTLEALVDAERRGEVVAPVDDLDWEDIFYIHDGCQWPSEPPAFKETMREYRAE
LRKLAERVMEAMDENLGLARGTIKDAFSSGGRHEPFFGTKVSHYPPCPRPDLITGLRAHTD
AGGVILLFQDDRVGGLEVLKDGQWTDVQPLAGAIVVNTGDQIEVLSNGRYRSAWHRVLPMR
DGNRRSIASFYNPANEATISPAAVQASGGDAYPKYVFGDYMDVYAKHKFQAKEPRFEAVKV
AAPKSSPAA
>XP_008662969.1 1-aminocyclopropane-1-carboxylate oxidase
[Zea mays]
MVVPVIDFSKLDGAERTETLAQIANGCEEWGFFQLVNHGIPLELLERVKKVCSDCYRLREA
GFKASEPVRTLEALVDAERRGEEVAPVDDLDWEDIFFIHDGCQWPSDPSAFKETMREYRAE
LRKLAERVMEAMDENLGLTKGTIKDAFSAGGRHEPFFGTKVSHYPPCPRPDLITGLRAHTD
AGGVILLFQDDRVGGLEVLKDGQWIDVQPLAGAIVINTGDQIEVLSNGRYRSAWHRVLPMR
DGNRRSIASFYNPANEATISPAAVQGSGGGETYPKYVFGDYMDVYVKQKFQAKEPRFEAVK
AAAPKSSPAA
>XP_008662968.1 1-aminocyclopropane-1-carboxylate oxidase
[Zea mays]
MVVPVIDFSKLDGAERTETLAQIANGCEEWGFFQLVNHGIPLELLERVKKVCSDCYRLREA
GFKVSEPVRTLEALVDAERRGEEVAPVDDLDWEDIFFIHDGCQWPSDPSAFKKTIREYRAE
LRKLAERVMEAMDENLGLTKGTIKDAFSGGGRHEPFFGTKVSHYPPCPRPDLITGLRAHTD
AGGVILLFQDDRVGGLEVLKDGQWIDVQPLAGAIVINTGDQIEVLSNGRYRSAWHRVLPMR
DGNRRSIASFYNPANEATISPAAVQGSSGGETYPKYVFGDYMDVYVKQKFQAKEPRFEAVK
AAAPKSSPAA
>NP_001395046.1 1-aminocyclopropane-1-carboxylate oxidase
isoform 2 [Zea mays]
MAIPVIDFSKLDGPERAETMAALAAGFEHVGFFQLVNTGISDDLLERVKKVCSDSYKLRDE
AFKDSNPAVKALTELVDKEIEDGLPARKIKDMDWEDVFTLHDDLPWPSNPPAFKETMMEYR

RELKKLAEKMLGVMEELLGLEEGHIRKAFSNDGEFEPFYGTKDDRFGGLQAQLPDGSWVDV
QPLENAIVINTGDQIEVLSNGRYKSAWHRILATRDGNRRSIASFYNPARLATIAPAIPAAG
VGDDDYPSFVFGNYMEVYVKQKFQPKAPRFEAMATTTTK
>NP_001146957.1 1-aminocyclopropane-1-carboxylate oxidase
isoform 1 [Zea mays]
MAIPVIDFSKLDGPERAETMAALAAGFEHVGFFQLVNTGISDDLLERVKKVCSDSYKLRDE
AFKDSNPAVKALTELVDKEIEDGLPARKIKDMDWEDVFTLHDDLPWPSNPPAFKETMMEYR
RELKKLAEKMLGVMEELLGLEEGHIRKAFSNDGEFEPFYGTKVSHYPPCPRPDLIDGLRAH
TDAGGLILLFQDDRFGGLQAQLPDGSWVDVQPLENAIVINTGDQIEVLSNGRYKSAWHRIL
ATRDGNRRSIASFYNPARLATIAPAIPAAGVGDDDYPSFVFGNYMEVYVKQKFQPKAPRFE
AMATTTTK
>XP_008646855.1 1-aminocyclopropane-1-carboxylate oxidase 1
[Zea mays]
MPSPRNARPRKPSRNARAPARVPHPRASPSPPIIPRLDCILRHPSPPHRGRSPFLAPPPSR
LRHPCRPRPPANLAAVKVESPRPSPSRVDPSPRLHPNDIHLHLIATSMACRAWPRLRPAHL
AVVKMESPRHPCRPRPPPNLTAVEVESPRPSPSRADPISRTPPSRLDCHAAASISTPRSRR
CRYRRTGRRTRLGPPMGGATGRRTRPGVGKCFVGLMEKNFYSSENAKILGCEKVPSNVDWE
CSFMYRHQPESNSHDIPELLRAMVSEYAEEVIKLAEQLAAAMSENLGLDKGYIEKEFSKPF
VGVKVAKYPRCSHPELVMGLREHTDAGGIILLFQDELIPGLEFLKDGRWMAVPPTQGNRIL
VNLGDQIEVITNGTYKSICHRVLPNKNGSRLSIATFYNPGADAIICPASKLTYPSQYRFQD
YLDFYSTTKFTDKVFRFQTTKAILK
>NP_001149427.2 1-aminocyclopropane-1-carboxylate oxidase 1
[Zea mays]
MTGPMEIPVIDLGGLNGGGEERSRTLAELHDACKDWGFFWVENHGVDAPLMDEVKRFVYGH
YEEHLEAKFYASALAMDLEAATRGDTDEKPSDEVDWESTYFIQHHPKTNVADFPEITPPTR
ETLDAYVAQMVSLAERLAECMSLNLGLPGAHVAATFAPPFVGTKFAMYPSCPRPELVWGLR
AHTDAGGIILLLQDDVVGGLEFLRAGAHWVPVGPTKGGRLFVNIGDQIEVLSAGAYRSVLH
RVAAGDQGRRLSVATFYNPGTDAVVAPAPRRDQDAGAAAYPGPYRFGDYLDYYQGTKFGDK
DARFQAVKKLLG
>XP_008661019.1 1-aminocyclopropane-1-carboxylate oxidase 1
[Zea mays]
MASPDLLFNLRNLFYLGAYQAAINNIDIPGLDAAAAAERDAIVFRSYIALGSYQVRTHPAS
GASAATSLQVVKLLALYLTGDKRRYVMKKINISKQNDKFQQTAYQEVTHVLLTHEDKKTLL
ERYTAFSKPSVGVKVAKYPRCSHPELVMGLREHTDAGGIILLFQDELIPGLEFLKDGRWMA
VPPTQGNRIFVNLGDQIEVITNGTYKSICHRVLPNKNGSRLSIATFYNPGADAIICPASKR
TYPSQYRFQDYLDFYSTTKFTDKVFRFQTTKAILK
>XP_008650459.1 1-aminocyclopropane-1-carboxylate oxidase 1
[Zea mays]
MEIPMIKMDQLHGEKRSETLSLLHNACAQWGFFWLENHGVDEDLMSKMKGLVNKHYEQDLE
KNFYSSENAKILGCEKVPSNVDWECSFMYRHQPESNSHDIPELLRAMVSEYAEEVIKLAEQ
LAAAMSENLGLDKGYIEKAFSKPSVGVKVAKYPRCSHPELVMGLREHTDAGGIILLFQDEL
IPGLEFLKDGRWMAVPPTQGNRILVNLGDQIEVITNGTYKSICHRVLPNKNGSRLSIATFY
NPGADAIICPASKLTYPSQYRFQDYLDFYSTAKFTDKVFRFQTTKAILK
>NP_001105234.1 acc oxidase [Zea mays]
MVVPVIDFSKLDGAERAETLAQIANGCEEWGFFQLVNHGIPLELLERVKKVCSDCYRLREA
GFKASEPVRTLEALVDAERRGEVVAPVDDLDWEDIFYIHDGCQWPSDPPAFKETMREYRAE
LRKLAERVMEAMDENLGLARGTIKDAFSGGGRHDPFFGTKVSHYPPCPRPDLITGLRAHTD
AGGVILLFQDDKVGGLEVLKDGEWTDVQPLEGAIVVNTGDQIEVLSNGLYRSAWHRVLPMR
DGNRRSIASFYNPANEATISPAAVQASGGDAYPKYLFGDYMDVYVKQKFQAKEPRFEAVKT
GAPKSSPAA
>XP_020401392.1 1-aminocyclopropane-1-carboxylate oxidase
[Zea mays]
MVVPVIDFSKLDGAERTETLAQIANGCEEWGFFQLVNHGIPLELLERVKKVCSDCYRLREA
GFKVSEPVRTLEALVDAERRGEEVAPVDDLDWEDIFFIHDGCQWPSDPSAFKKTIREYRAE
LRKLAERVMEAMDENLGLTKGTIKDAFSGGGRHEPFFGTKVSHYPPCPRPDLITGLRAHTD

AGGVILLFQDDRVGGLEVLKDGQWIDVQPLAGAIVINTGDQIEVLSNGRYRSAWHRVLPMR
DGNRRSIASFYNPANEATISPAAVQGSSGGETYPKYVFGDYMDVYVKQKFQAKEPRFEAVK
AAAPKSSPAA

>NP_001105235.1 ACC oxidase20 [Zea mays]
MAATVSFPVVNMEKLETEERDTAMAVIRDACENWGFFELLNHGISHELMDEVERLTKAHYA
TFREAKFQEFAARTLAAAGDEGADVSDVDWESTFFVRHLPASNLADLPDVDDHYRQVMKQF
ASEVQKLSEKVLDLLCENLGLEPGYLKAAFAGSDGGPTFGTKVSAYPPCPRPDLVAGLRAH
TDAGGLILLLQDDQVSGLQLLRGGDGGEWVDVPPLRHAIVANVGDQLEVVTNGRYKSAVHR
VLARPDGNRMSVASFYNPGADAVIFPAPALVGEEERAEKKATTYPRFVFEDYMNLYARHKF
EAKEPRFEAMKSSAIATA

**Supplementary Table S5 | List of protein sequence (n=70) from ten plant species used protein multiple sequencer alignment.** Sequence included are from *Arabidopsis thaliana, Malus domestica, Petunia hybrida, Solanum lycopersicum, Nicotiana benthamiana, Glycine max, Triticum aestivum, Zea mays, Amborella trichopoda, Marchantia polymorpha*).
